# Initial amino acid:codon assignments and strength of codon:anticodon binding

**DOI:** 10.1101/2023.12.18.572171

**Authors:** Meng Su, Samuel J. Roberts, John D. Sutherland

## Abstract

The ribosome brings 3′-aminoacyl-tRNA and 3′-peptidyl-tRNAs together to enable peptidyl transfer by binding them in two major ways. Firstly, their anticodon loops are bound to mRNA, itself anchored at the ribosomal subunit interface, by contiguous anticodon:codon pairing augmented by interactions with the decoding centre of the small ribosomal subunit. Secondly, their acceptor stems are bound by the peptidyl transferase centre which aligns the 3′-aminoacyl- and 3′-peptidyl-termini for optimal interaction of the nucleophilic amino group and electrophilic ester carbonyl group. Reasoning that intrinsic codon:anticodon binding might have been a major contributor to bringing tRNA 3′-termini into proximity at an early stage of ribosomal peptide synthesis, we wondered if primordial amino acids might have been assigned to those codons that bind the corresponding anticodon loops most tightly. By measuring the binding of anticodon stem loops to short oligonucleotides, we determined that family box codon:anticodon pairings are typically tighter than split box codon:anticodon pairings. Furthermore, we find that two family box anticodon stem loops can tightly bind a pair of contiguous codons simultaneously whereas two split box anticodon stem loops cannot. The amino acids assigned to family boxes correspond to those accessible by what has been termed cyanosulfidic chemistry, supporting the contention that these limited amino acids might have been the first used in primordial coded peptide synthesis.

**Table of Contents (TOC) graphic:** 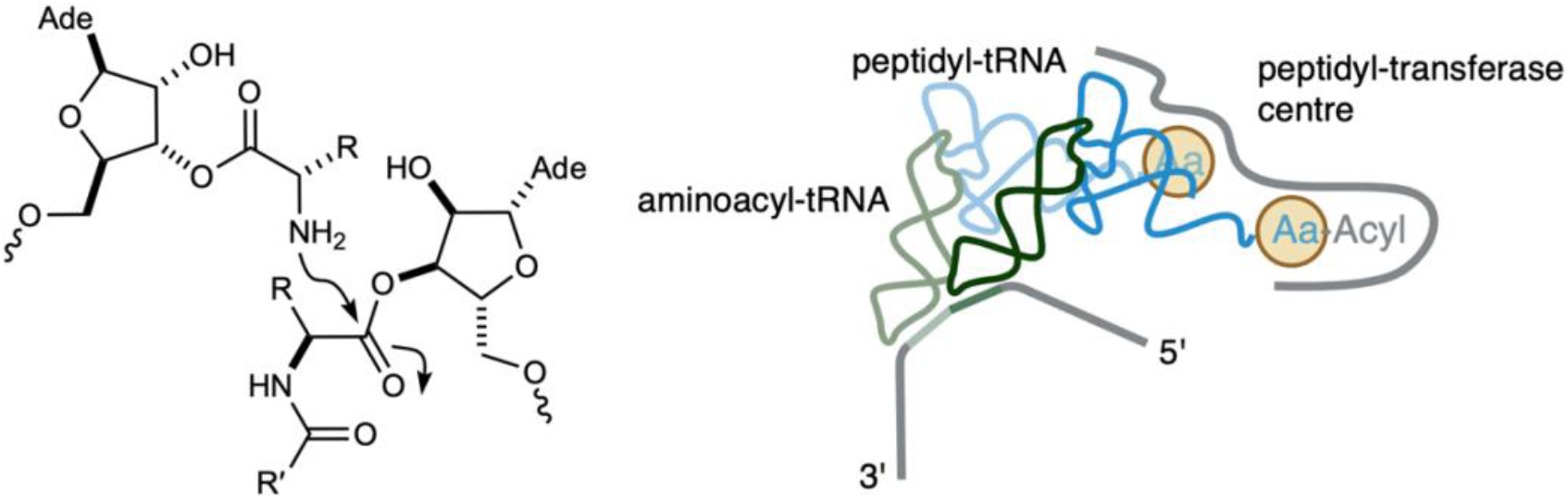

## Introduction

Molecular recognition events control the specificity of the writing and reading steps of translation according to the genetic code. The code is written by aminoacyl-tRNA synthetases recognising specific tRNAs and attaching cognate amino acids to their 3′-termini. The code is read by sequential anticodon:codon recognition in the decoding centre of the ribosome. How such a process could have evolved is an intriguing question with the mechanism of individual amino acid-codon assignments at its heart. Before the advent of aminoacyl-tRNA synthetases – be they ribozymes or enzymes – it is difficult to imagine selective tRNA aminoacylation unless molecular recognition between the amino acid residue and the tRNA itself was responsible for this. If such recognition took place, parsimony suggests that it should have been between the amino acid residue and the anticodon, but this is countered by the fact that the 3′-terminal aminoacylation site is some 75Å distant from the anticodon. Having a trinucleotide sequence elsewhere in the tRNA act as the aminoacylation specificity control element alleviates this distance constraint,^1^ but decouples the nature of the aminoacyl residue and the sequence of the anticodon, necessitating an alternate mechanism for amino acid-codon assignments.

Our recent finding that the terminal trinucleotide sequence of a tRNA acceptor stem analogue influences the specificity and stereochemistry of 3′-terminal aminoacylation suggests that loosely coded aminoacyl-tRNA acceptor stem-overhang domains could have been produced by prebiotic chemistry.^1,2^ However, the degree of intrinsic coding is low, and we suspect that higher fidelity coding is unlikely without extrinsic auxiliary control elements such as ribozymes. To us, this implies that if the initial loosely coded aminoacyl-tRNAs had participated in the early stages of ribosomal peptide synthesis, the short peptides thus produced would not have been advantageous to the system per se. We have thus started to investigate the possibility that the primary function of early peptide synthesis was not the produced oligopeptide. Rather, that in a scenario of adventitiously produced aminoacyl-RNA molecules, peptide bond formation would have provided a mechanism to deblock RNA 3′-termini so that they were rendered extension/ligation competent. For example, a self-priming substrate for chemical or in *trans* ribozymic copying in which a duplex 3′-overhang primes strand displacement synthesis via a folded-back conformation – such overhangs also being susceptible to aminoacylation by a variety of mechanisms.^2,3^ According to this idea, the selective pressure behind the development of transpeptidation would have been to allow more efficient RNA replication – peptides produced by the process would simply have been waste products. Second-order transpeptidation would have had to be faster than pseudo-first-order hydrolysis of aminoacyl- and peptidyl-RNAs to be selected for this purpose. One can envisage that the proto-peptidyl transferase centre of the ribosome initially catalyzed transpeptidation between acceptor stem-overhangs simply by binding them to increase their effective molarities. Subsequent fusion of the acceptor stem-overhang domains with anticodon stem-loop domains could have facilitated this process if it further contributed to the proximity required for transpeptidation (Fig. 1a).^4^ One way in which this could have occurred would have been through anticodon binding of two such fusions to contiguous codons on a short piece of single-stranded RNA (ssRNA).^5^ In investigations of extant biochemistry, it has been shown that two tRNA^Phe^(UUC) molecules do not appreciably bind contiguous codons on a short oligonucleotide simultaneously absent the ribosomal decoding centre.^6^ However, we reasoned that this might be a specific case of weak binding between Phe-codon (UUC) and anticodon (GAA) loops. Accordingly, we decided to investigate the effective molarity of aminoacyl/peptidyl-RNAs required to favour uncatalyzed transpeptidation relative to hydrolysis and to study the energetics of codon:anticodon loop binding across the whole genetic code.

**Fig. 1.**
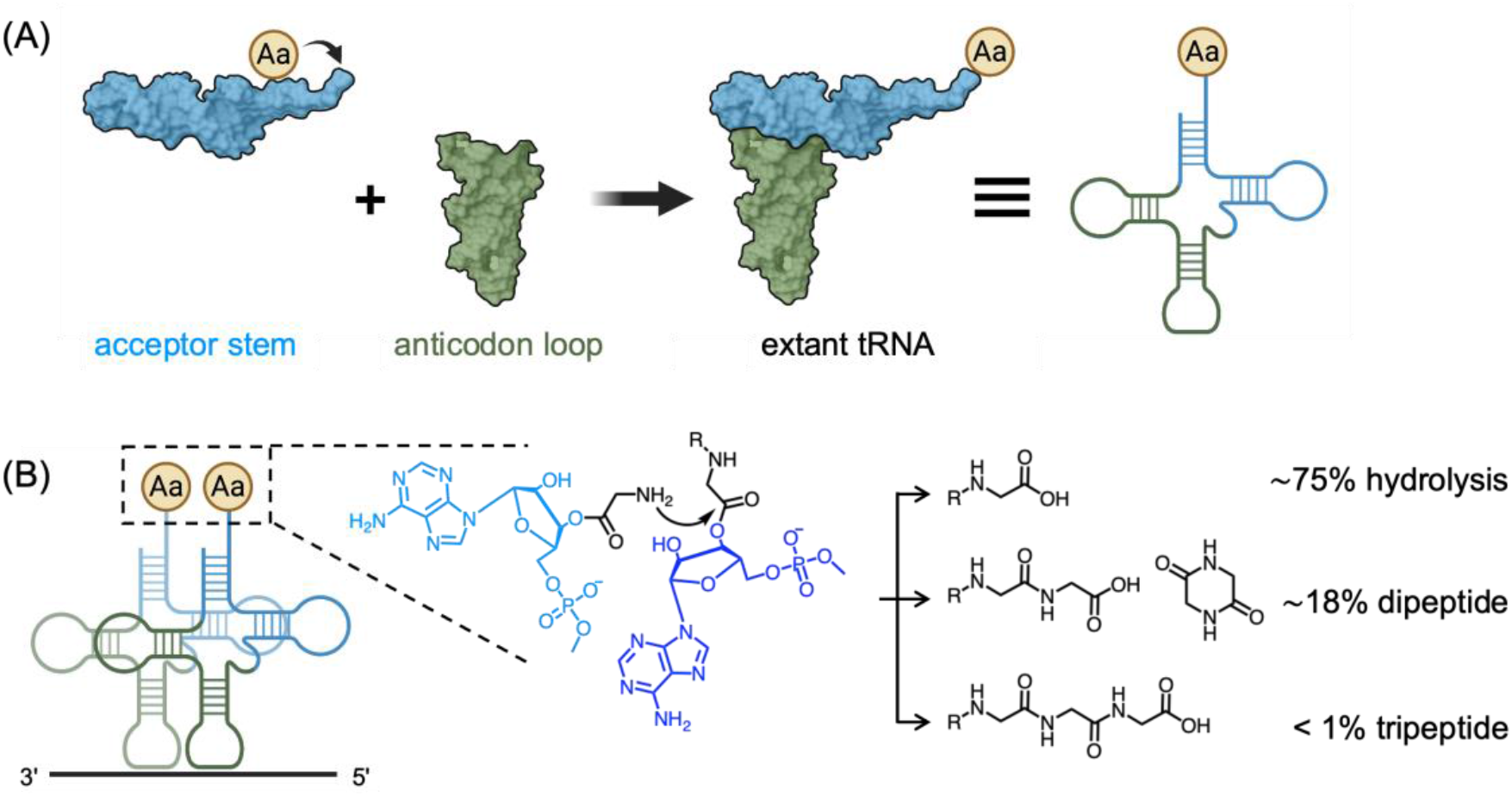
Evolution of tRNA and the mechanism of transpeptidation. (A) Schematic representation of aminoacyl-transfer on the acceptor stem-overhang^1^ and the fusion of the acceptor stem domain and anticodon stem-loop domain; (B) The high effective concentration of aminoacylated tRNAs and the power of exclusion of water by the ribosome during translation is demonstrated by the incubation of 0.4 M 2′/3′-glycyl-adenosine-5′-*O*-methylphosphate in solution producing a minority of di- and tri-peptide species compared to free glycine. Created with BioRender.com, tRNA PDB ID: 4TNA. R = H or formyl group.

## Results and Discussion

The first assessment of the effective molarity of tRNA termini required to ensure that transpeptidation can compete with hydrolysis by 55 M water, was published by Weber and Orgel.^7,8^ These authors found that 0.4 M 2′/3′-glycyl-adenosine-5′-(O-methylphosphate) gave rise to diglycine and its diketopiperazine (DKP) in ∼15% combined yield along with a trace of triglycine after 2 hours at pH 8.2 or 9. We confirmed their results by ^1^H-NMR analysis of similar reactions (415 mM, pH 8.2), observing yields of 6% diglycine, 12% DKP, and <0.5% triglycine (Fig. 1b, Fig. S1). Incubation of equimolar glycine with 2′/3′-glycyl-adenosine-5′-*O*-methylphosphate demonstrated a substantial reaction between free amino acids and aminoacyl-nucleotides (Fig. S2). This indicates that a translational system in the presence of free amino acids – either initially endogenous, or those produced by substantial aminoacyl ester hydrolysis – would produce high yields of uncoded peptides. This data was supported by functionalization of the crude products of our reactions with dansyl groups and analysis by HPLC, which demonstrated similar yields for diglycine and triglycine (Table S1, S2, and Fig. S3). The vast majority of the aminoacyl esters, however, underwent hydrolysis to free glycine as also reported by Orgel. Dipeptidyl species were hardly detectable by HPLC or NMR when the aminoacyl ester concentration was reduced by more than an order of magnitude (35 mM, Fig. S4).

Diketopiperazine formation is a known hurdle to be overcome for the production of oligopeptides.^9^ Incubating conventionally synthesized 2′/3′-diglycyl-adenosine-5′-*O*-methyl-phosphate at a low concentration at pH 8.2 gave DKPs and linear dipeptides in a 3:1 ratio (Fig. S6). One of the ways biology overcomes the DKP problem is by initiating peptide synthesis with an *N-*formyl amino acid. To reduce DKP formation in our model system and to investigate whether transpeptidation is more efficient when one reaction partner is *N*-acylated, we incubated a 1:1 mixture of 2′/3′-glycyl-adenosine-5′-*O*-methylphosphate with 2′/3′-*N*-formyl-glycyl-adenosine-5′-*O*-methylphosphate at a combined concentration of 415 mM. Whilst this reduced DKP formation and consequently improved the yields of linear peptides the reaction still suffered from similar low total peptide yields (1.5% diglycine, 5.4% *N*-formyl-diglycine, 5.2% DKP, ∼0.1% tripeptides, Fig. S7). We thus conclude that uncatalyzed transpeptidation, even using a combination of a free nucleotide aminoacyl ester and an *N*-acylated one, is an intrinsically inefficient process in water. Any early transpeptidation catalyst would therefore have had to bring about mutual proximity of the reaction partners equivalent to an effective molarity of both species of the order of hundreds of mM.

There have been several recent attempts to recapitulate the function of the extant ribosomal peptidyl-transferase centre using heavily truncated variants of the large ribosomal subunit, but thus far only trace transpeptidation activity has been detected.^10,11^ We have also started efforts to (re)produce a minimal peptidyl-transferase centre capable of inducing proximity between a free aminoacyl-nucleotide and an *N*-blocked one. Given the high proximity requirement for transpeptidation to outcompete hydrolysis, we anticipate that any primitive entropic catalyst will benefit from other contributions to the proximity of the reaction partners. As mentioned above, one such additional contribution could be afforded by fusing the acceptor stem-overhang domains to anticodon domains. Simultaneous binding of two such fusions to contiguous codons on a short oligonucleotide would augment the proximity induced by the minimal peptidyl-transferase centre. Production of intact tRNAs by this sort of domain fusion has been suggested previously,^4,12^ but the potential benefit afforded to transpeptidation caused by contiguous anticodon binding to ssRNA has not been alluded to. It is reasonable to expect this benefit to scale with the strength of binding between tRNAs codons and anticodons.^5^

The energies of the codon:anticodon trinucleotide minihelices have been predicted on the basis of the nearest neighbour model.^13-15^ However, since there is limited experimental data regarding the binding of tRNA anticodon loops to codons, we investigated their binding affinity using biolayer interferometry (BLI).

First, we prepared biotin-labeled ssRNAs containing our desired codon sequences. We also prepared RNA hairpins with an identical 5-base pair stem and differing 7-mer loops containing 64 anticodon triplets (Table S3-5). The remaining four nucleotides of the loop were limited to bases equivalent to the anticodon loop residues 32C, 33U, 37A, and 38A of a tRNA based on the consensus sequences of existing tRNAs.^16^ However, it is known that variations at these positions can alter the geometry of the anticodon loops and subsequently influence the binding strength (Fig. S8).^17^ Using these synthetic hairpins, we determined the association (*k*_*on*_) and dissociation rate constants (*k*_*off*_) for binding of the anticodon loop to the corresponding immobilized ssRNA using BLI, thus enabling calculation of the equilibrium dissociation constant (*K*_*D*_) (Fig. 2, Fig. S9, Table S6). For all the measurable *K*_*D*_s, the average value of (four-fold degenerate) family-box^18^ codon:anticodon binding is 100 μM, while the average of split-box codon:anticodon interaction is 460 μM (P<0.0001, Fig. 2A). As we had anticipated, the previously studied UUC codon:GAA anticodon pairing showed weak binding as evidenced by a high *K*_*D*_ (470 μM). The disparity between the two classes of codons is most substantial when codons end with cytosine (average 49 μM vs 486 μM, P<0.005, Fig. 2B, C, Fig. S10). With this terminal codon base, a threshold of 90 μM can be set to divide the two classes of codons. The precise rules for codons ending with bases other than C are less clear. Further data gained using isothermal titration calorimetry (ITC), with all oligonucleotides in free solution, substantially agree with the BLI results (Fig. S11, Table S7-9). Stronger codon:anticodon binding pairs would have provided a more pronounced effect on the catalysis of early transpeptidation. Weaker pairs would have been less useful until auxiliary contributors, for example, nucleobase modifications, emerged to compensate for their lower binding affinity. We note that the third position of the codon has been hypothesized to be either promiscuous (through various wobble interactions) or inoperative in early genetic codes.^18-19^

**Fig. 2.**
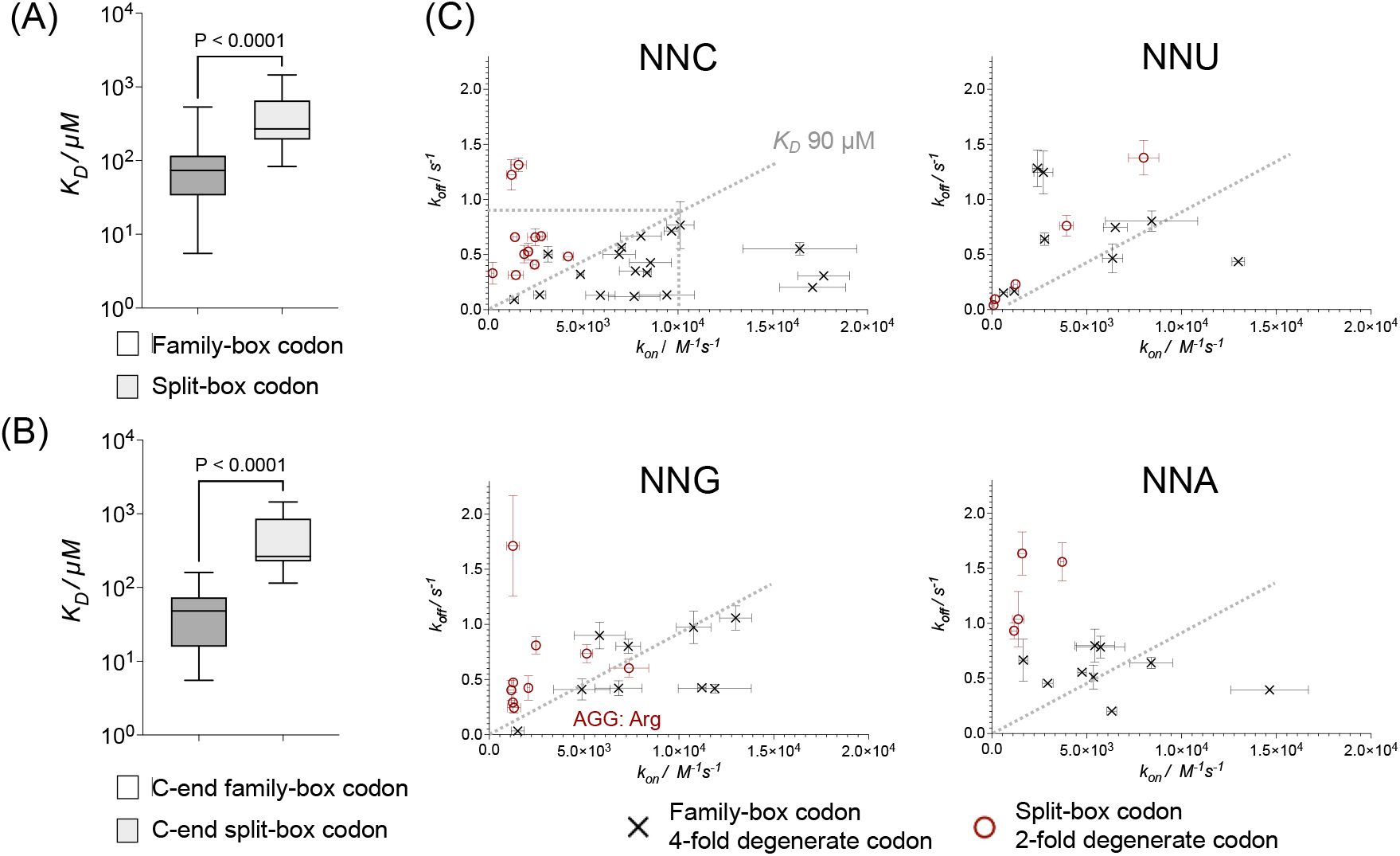
Equilibrium dissociation constant (*K*_*D*_) of codons;anticodon loop binding as determined by BLI. Box and whisker plots of *K*_*D*_ of (A) all family-box codons and split-box codons and (B) C-ending family-box codons and split-box codons; (C) plots of association constants and dissociation constants of measured anti-codon loop:codon ending with C/U/G/A in two codon classes.

Some of our earlier research has focused on the prebiotic synthesis of Strecker amino acid precursors, which are products of reductive homologation of hydrogen cyanide.^20^ The precursors of Asp, Asn, Glu and Gln derived from cyanoacetylene which we originally sourced by in situ stoichiometric oxidative coupling of acetylene and hydrogen cyanide by Cu(II). However, we now favour a model in which atmospherically-derived cyanoacetylene is stored as a dicyanoimidazole adduct for exclusive incorporation into pyrimidine ribonucleotides,^21^ so our current view is that these amino acid precursors are not part of the cyanosulfidic set. The remaining Strecker precursors correspond precisely with the amino acids encoded by family box codons in extant biology. Hence we suggest that the current assignment of amino acids to codons is in part due to this correlation between the prebiotically accessible early amino acids and the stronger family box codon:anticodon interactions. There are two split box examples where codon:anticodon binding is similar strength to those of the family boxes; AGG (85 μM) and GAG (140 μM) encoding Arg and Glu. Arg is also a family box amino acid, but Glu is only encoded by a split box in extant biochemistry. We thus contend that this suggests that the family box amino acids were used in the earliest forms of coded peptide synthesis and that Glu was either post-biosynthetically assigned to GAG, or, if a plausible cyanosulfidic synthesis is uncovered, was also one of the initially assigned prebiotic amino acids.

Transpeptidation requires the proximity of two stem-overhang domains. This proximity would be most enhanced if both overhangs were co-immobilised, much as they are in modern biology when two tRNAs are bound to contiguous codons. Again using BLI, we demonstrated that two anticodon loop domains can bind to an ssRNA simultaneously without additional partners, but that the binding is typically only strong when both pairings involve family box codons (Fig. 3A).

**Fig. 3.**
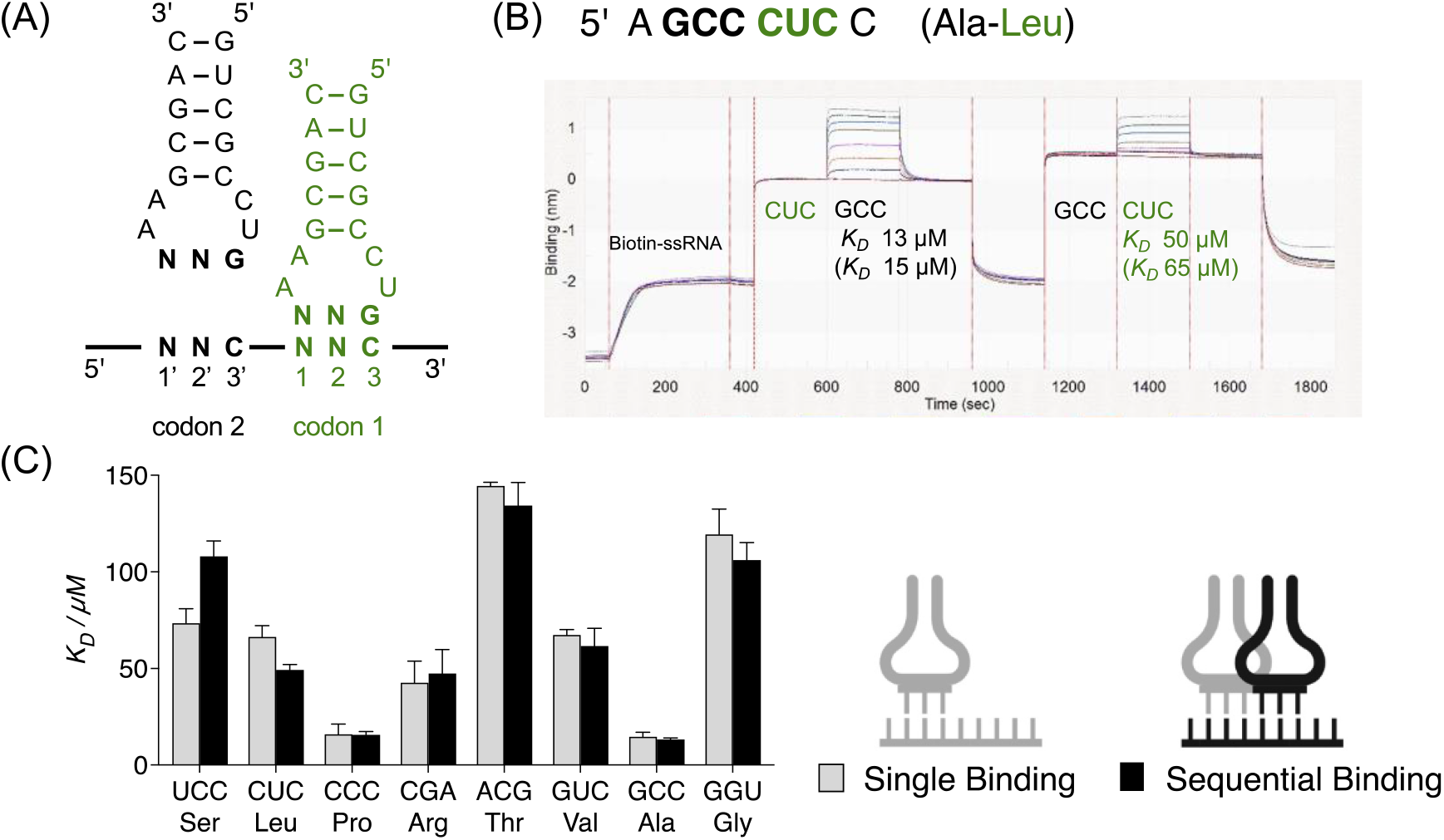
Sequential binding of two anticodon loops to ssRNA. (A) Schematic representation of anticodon loops binding to a biotin-labeled ssRNA containing the respective codons; (B) typical BLI curves of ssRNA containing two family-box codons. Equilibrium dissociation constants of the anticodon:codon complex after the first loop binding to the ssRNA (i.e., sequential binding) are annotated beneath their respective codons. Bracketed equilibrium dissociation constants are those single anticodon loop:ssRNA binding; (C) bar chart comparing the *K*_*D*_ of single binding and sequential binding of one orthogonal anticodon loop subset (i.e. a subset of anticodon loops which show significant (*K*_*D*_ <130 μM) binding to ssRNA and are not able to bind each other by loop-loop interactions). Conditions: 200 μM for the 1st anticodon loop, 0–400 μM for the 2nd anticodon loop, 100 mM NaCl, 100 mM MgCl_2_, 50 mM HEPES, pH 7.2, 20 ºC. Btn, Biotin.

We immobilised a biotinylated oligoribonucleotide 5’-AGCCCUCC containing two contiguous family-box codons ending in C (GCC and CUC, Fig. 3B, Table S10). We saturated the CUC codon with a corresponding hairpin RNA to give a complex which was subsequently incubated with a second hairpin designed to bind the GCC codon. The *K*_*D*_ for the second binding (GCC) was found to be slightly lower (13 μM) than when binding to ssRNA alone (15 μM), suggesting there was no significant synergy in the termolecular interaction. We were also able to reverse the order of hairpin addition and saw a slightly stronger binding for the second hairpin (CUC *K*_*D*_ = 50 μM, compared to 65 μM for when binding singly to ssRNA).

It has previously been observed that two complementary anticodon loops can bind each other tightly forming a loop-loop complex.^22^ We also found that specific pairs of anticodons can bind each other with affinities higher than we had seen for codon:anticodon binding (e.g. GCC:GGC 1.0 μM; GCC:GGU 1.7 μM; Fig. S12, Table S7). To avoid loop-loop complexes, we hypothesized that only a subset of non-interacting anticodon loops could have been used at the origin of genetic coding. Through exhaustive search, we identified 14 such subsets containing 8 anticodons each encoding a different family box amino acid of sufficiently high binding affinity (<130 μM), that cannot bind any of the other anticodons in the same subset significantly (Table S11, S12). We selected one subset and, using four different ssRNAs, measured the binding affinity to matching contiguous codons from that subset (Fig. 3C, Fig. S13, Table S10). The affinities for sequential binding remained largely similar to those of single binding. Those showing a slight weakening in binding affinity could be accounted for by steric clash between the first and second anticodon loop. A strengthening in binding affinity of the second hairpin could be accounted for by an energetic reorganisation of the ssRNA template, much like the kinking of messenger RNA in the ribosome.^23^

We then tested an immobilised ssRNA containing two contiguous split-box codons UUC and CAU (Table S10, Fig. S14). The weak ssRNA binding affinity of the corresponding split-box anticodon loops and the limit of their solubility meant pre-saturation by one hairpin was not possible. Instead, both hairpins were incubated simultaneously affording an apparent *K*_*D*_ (190 μM) nearly identical to the single site binding for the stronger codon (180 μM). This suggests only single-site occupation could be observed, consistent with the observations of Labuda *et al*.^5^. We found that similar experiments using ITC qualitatively complemented the results seen with BLI (Fig. S11, Table S7 and S9). Furthermore, when a split-box anticodon loop was added to a preformed complex of family-box anticodon bound to an ssRNA (5’-AGAC<u>GCC</u>C; family box codon underlined), we observed reduced binding of the split-box anticodon loop (sequential binding *K*_*D*_ = 580 μM vs single binding *K*_*D*_ = 240 μM (Fig. S13, Table S10). When the order of addition was reversed, family-box binding also decreased by a factor 2.4 (sequential binding *K*_*D*_ = 34 μM vs single binding *K*_*D*_ = 14 μM). A second example, this time using a ssRNA containing a lower-affinity split-box codon (5’-A<u>GUC</u>UACC; family box codon underlined), also demonstrated reduced binding. Preincubation of an anticodon loop to UAC reduced the subsequent binding affinity to GUC by a factor of three, from 80 μM to 240 μM, whilst reversing the order made the split-box binding event unmeasurable. This case is remarkable in that the binding to GUC is reduced even though the affinity between the split box codon UAC and its anticodon complement is extremely low (3400 μM). In both these examples, it is plausible for partial anticodon:anticodon interactions between the different hairpins to occur (<u>G</u>*G*<u>C</u>:<u>G</u>*U*<u>C</u> and G<u>AC</u>:<u>GU</u>A; Watson-Crick binding underlined, wobble italicised). The weakened binding demonstrates there would exist an evolutionary driving force to restrict to an orthogonal set of family box anticodon loops in a background lacking significantly complementary split-box anticodon loops, thereby avoiding inhibition of the former’s binding to ssRNAs.

Coded peptide synthesis by translation requires two tRNAs to be bound adjacently to a ssRNA. Our results (albeit from a limited number of pairwise combinations) suggests that the earliest forms of translation could only utilise subsets of orthogonal family box codons. Incorporation of split-box codons would have required further stabilisation, for instance by the evolution of the decoding centre in the rRNA small subunit.^24^ Furthermore, as more codons were assigned, nascent biology would have had to evolve strategies to avoid the inhibition of codon binding by anticodon:anticodon complementarity.

## Conclusion

Coded peptide synthesis by translation is ubiquitous in extant biology and its emergence would have constituted an extremely beneficial evolutionary innovation. Key to the mechanism of translation is spatial restriction of two aminoacyl residues to enable transpeptidation. Complementing previous work, we have shown that an effective concentration of aminoacyl (oligo-) nucleotides of hundreds of mM is required to form peptide bonds at a synthetically useful level in water.

The ribosome constrains transpeptidation substrates, in part, by localising aminoacyl-tRNA and peptidyl-tRNA to a ssRNA template through codon:anticodon binding. A proto-translational mechanism could have benefited greatly from such an increase in effective molarity in proportion to the binding strength. The interaction of codons with anticodon loops has not been studied previously in the absence of ribosomal architecture, but various relationships have been hypothesised.

Herein, we present experimental data demonstrating the binding strength between anticodon loop domains and their complementary codons on ssRNA. In general, our findings indicate that family-box codons exhibit stronger binding compared to codons from split-boxes. This is most evident for codons ending in base C. Furthermore, contrary to previous reports,^5^ two distinct anticodon loops can simultaneously bind to an RNA strand containing contiguous family-box codons without auxiliary components like the ribosome. 14 different subsets of orthogonal, strong binding affinity, family-box codons encoding all eight prebiotic amino acids can be envisaged, which avoid the problems of loop-loop complexation whilst maintaining high binding affinity to ssRNA. The binding of the second hairpin is generally slightly weakened, suggesting that inherent strong binding affinity of each individual hairpin is the essential property required for binding two hairpins contiguously. ssRNA containing weak-binding split-box codons failed to simultaneously bind two respective anticodon loops.

These findings provide a strong argument for the selection of strong binding family box codons over split-box codons at the origin of translation. Expansion of the early genetic code would have required additional components to stabilize weaker binding loops. Combining this selection mechanism with the prebiotic availability of amino acids produced by the cyanosulfidic scenario,^20^ would explain why glycine, alanine, valine, leucine, serine, threonine, arginine, and proline are all assigned to family box codons in modern biology. In light of our recent work on coded aminoacylation,^1^ which family box codons code for each amino acid would be dependent on the fusion of acceptor stems and anticodon domains. It is possible that the precise relationship between any particular family box codon and its encoded amino acid is a ‘frozen accident’.^25^

Since the evolution of functional coded peptides by transpeptidation of (oligo)nucleotide aminoacyl esters would have faced extreme challenges, we propose that transpeptidation was initially developed instead as an RNA deblocking mechanism. Activation of RNA in the presence of (*N*-acyl-)amino acids would also lead to RNA diol termini becoming (*N*-acyl-)aminoacylated.^1,26^ Acylated RNA termini would prevent extension/ligation chemistries, preventing RNA replication until the (*N*-acyl-)aminoacyl group was removed. Development of a catalyst which freed RNA termini by transpeptidation would have been evolutionarily favoured because it would have increased the efficiency of RNA replication. The initial catalyst might have acted on (*N*-acyl-)aminoacyl RNA domains in free solution, but would have been greatly improved upon by conformational restriction of the two hairpin RNAs. Fusing aminoacyl stem overhangs to family box anticodon-containing hairpins would have enabled ssRNA chains to act as a template, increasing the effective concentration of (*N*-acyl-)aminoacyl groups. The strength of template binding could have been additionally improved by later emerging auxiliary molecules. These auxiliaries could also develop further, along with anticodon loop nucleotide modifications, to stabilise split-box codon:anticodon interactions thus enabling the use of the full set of 64 possible codons. The production of waste peptides with partial coding from various mechanisms, could later undergo exaptation to produce useful peptides, hence building the modern translation system. Understanding the nature of the initial transpeptidation catalyst is essential and a current focus in our laboratory.

## ASSOCIATED CONTENT

### Supporting Information

The data that support the findings of this study are available within its Supporting Information.

The Supporting Information is available free of charge at Materials and methods, supplementary data and figures (PDF)

All the codes used in this study are available in a GitHub respository at: https://github.com/nieseln/Codon-select.git

## Notes

The authors declare no competing financial interest.

## ACKNOWLEDGMENTS

The authors thank Dr. Chris Johnson and Dr. Stephen McLaughlin for their technical support on ITC and BLI experiments. This work is supported by the Medical Research Council MC_UP_A024_1009 (JDS), and the Simons Foundation 290362 (JDS).

## REFERENCE

1. Su, M.; Schmitt, C.; Liu, Z.; Roberts, S. J.; Liu, K. C.; Röder, K.; Jäschke, A.; Wales, D. J.; Sutherland J. D. Triplet-encoded prebiotic RNA aminoacylation. J. Am. Chem. Soc., 2023, 145 (29), 15971–15980. 10.1021/jacs.3c03931

2. Wu, L.-F.; Su, M.; Liu, Z.; Bjork, S. J.; & Sutherland, J. D. Interstrand aminoacyl transfer in a tRNA acceptor stem-overhang mimic. J. Am. Chem. Soc. 2021, 143 (30), 11836–11842. 10.1021/jacs.1c05746

3. Roberts, S. J.; Liu, Z.; Sutherland, J. D. Potentially Prebiotic Synthesis of Aminoacyl-RNA via a Bridging Phosphoramidate-Ester Intermediate. J. Am. Chem. Soc. 2022, 144 (42), 4254–4259, DOI: 10.1021/jacs.2c00772

4. Schimmel, P.; Giegé, R.; Moras, D.; Yokoyama, S. An operational RNA code for amino acids and possible relationship to genetic code. Proc. Natl. Acad. Sci. U. S. A. 1993, 90 (19), 8763–8768. 10.1073/pnas.90.19.8763

5. Lehmann, J. Physico-chemical constraints connected with the coding properties of the genetic system. J. Theor. Biol. 2000, 202 (2), 129–144. 10.1006/jtbi.1999.1045

6. Labuda, D.; Striker, G.; Grosjean, H.; Porschke, D. Mechanism of codon recognition by transfer RNA studied with oligonucleotides larger than triplets. Nucleic Acids Res., 1985, 13 (10), 3667–3683. 10.1093/nar/13.10.3667

7. Weber, A. L.; Orgel, L. E. The formation of peptides from the 2′(3′)-glycyl ester of a nucleotide. J. Mol. Evol., 1978, 11 (3), 189–198. 10.1007/BF01734480.

8. Weber, A. L.; Orgel, L. E. The formation of dipeptides from amino acids and the 2′(3′)-glycyl ester of an adenylate. J. Mol. Evol., 1979, 13 (3), 185–191. 10.1007/BF01739478

9. Steinberg, S. M.; Bada, J. L. Peptide decomposition in the neutral pH region via the formation of diketopiperazines. J. Org. Chem., 1983, 48 (13), 2295–2298. 10.1021/jo00161a036

10. Bose, T.; Fridkin, G.; Davidovich, C.; Krupkin, M.; Dinger, N.; Falkovich, A. H.; Peleg, Y.; Agmon, I.; Bashan, A.; Yonath, A. Origin of life: protoribosome forms peptide bonds and links RNA and protein dominated worlds. Nucleic Acids Res., 2022, 50 (4), 1815–1828. 10.1093/nar/gkac052

11. Kawabata, M.; Kawashima, K.; Mutsuro-Aoki, H.; Ando, T., Umehara, T.; Tamura, K. Peptide Bond Formation between Aminoacyl-Minihelices by a Scaffold Derived from the Peptidyl Transferase Center. Life 2022, 12 (4), 573. 10.3390/life12040573

12. Maizels, N.; Weiner, A. M. Phylogeny from function: evidence from the molecular fossil record that tRNA originated in replication, not translation. Proc. Natl. Acad. Sci. U. S. A. 1994, 91 (15), 6729–6734. 10.1073/pnas.91.15.6729

13. Grosjean, H.; Westhof, E. An integrated, structure- and energy-based view of the genetic code. Nucleic Acids Res., 2016, 44 (17), 8020–8040. 10.1093/nar/gkw608

14. Xia, T.; SantaLucia, J.; Burkard, M. E.; Kierzek, R.; Schroeder, S. J.; Jiao, X.; Cox, C.; Turner, D. H. Thermodynamic parameters for an expanded nearest-neighbour model for the formation of RNA duplexes with Watson-Crick base pairs. Biochemistry, 1998, 37 (42), 14719–14735. 10.1021/bi9809425

15. Mathews, D. H.; Sabina, J.; Zuker, M.; Turner, D. H. Expanded sequence dependence of thermodynamic parameters improves prediction of RNA secondary structure. J. Mol. Biol., 1999, 288 (5), 911–940. 10.1006/jmbi.1999.2700

16. Auffinger, P.; Westhof, E. Singly and bifurcated hydrogen-bonded base-pairs in tRNA anticodon hairpins and ribozymes. J. Mol. Biol., 1999, 292 (3), 467–483. 10.1006/jmbi.1999.3080

17. Auffinger P.; Westhof E. An extended structural signature for the tRNA anticodon loop. RNA, 2001, 7 (3), 334–341. 10.1017/s1355838201002382

18. Lagerkvist U. “Two out of three”: an alternative method for codon reading. Proc. Natl. Acad. Sci. U. S. A., 1978, 75 (4), 1759–1762. 10.1073/pnas.75.4.1759

19. Crick, F. H. Codon--anticodon pairing: the wobble hypothesis. J. Mol. Biol., 1966, 19 (2), 548–555. 10.1016/S0022-2836(66)80022-0

20. Patel, B. H.; Percivalle, C.; Ritson, D. J.; Duffy, C. D.; Sutherland, J. D. Common origins of RNA, protein and lipid precursors in a cyanosulfidic protometabolism. Nat. Chem., 2015, 7 (4), 301–307. 10.1038/nchem.2202

21. Ritson, D. J.; Poplawski, M. W.; Bond, A. D.; Sutherland, J. D. Azoles as Auxiliaries and Intermediates in Prebiotic Nucleoside Synthesis. J. Am. Chem. Soc., 2022, 144 (42), 19447–19455, DOI: 10.1021/jacs.2c07774

22. Grosjean, H.; Söll, D. G.; Crothers, D. M. Studies of the complex between transfer RNAs with complementary anticodons: I. Origins of enhanced affinity between complementary triplets. J. Mol. Biol., 1976, 103 (3), 499–519.

23. Selmer, M.; Dunham, C. M.; Murphy, F. V. IV.; Weixlbaumer, A.; Petry. S.; Kelley, C.; Weir, J. R.; Ramakrishnan, V. Structure of the 70S ribosome complexed with mRNA and tRNA. Science, 2006, 313 (5795), 1935–1942. 10.1126/science.1131127

24. Schluenzen, F.; Tocilj, A.; Zarivach, R.; Harms, J.; Gluehmann, M.; Janell, D.; Bashan, A.; Bartels, H.; Agmon, I.; Franceschi, F.; Yonath, A. Structure of functionally activated small ribosomal subunit at 3.3 angstroms resolution. Cell, 2000, 102 (5), 615–623. 10.1016/S0092-8674(00)00084-2

25. Crick, F. H. The origin of the genetic code. J. Mol. Biol., 1968, 38 (3), 367–379, DOI: 10.1016/0022-2836(68)90392-6

26. Weber, A. L.; Orgel L. E. Amino acid activation with adenosine 5’-Phosphorimidazolide. J. Mol. Evol., 1978, 11 (1), 9–16. 10.1007/BF01768020

