## Supplementary Information for "Initial amino acid:codon assignments and strength of codon:anticodon binding"

**This PDF file includes:**

Figures S1 to S32

Tables S1 to S11

### Materials and Methods

#### Materials and general methods

Reagents and solvents were obtained from *Acros Organics*, *Santa Cruz Biotechnology*, *Sigma-Aldrich*, and were used without further purification. pH was measured using a *Mettler Toledo* SevenEasy pH Meter S20 combined with a *ThermoFisher Scientific* Orion 8103BN Ross semi-micro pH electrode or an Orion™ 9810BN Micro pH Electrode. $^{1}$H-, and $^{13}$C-nuclear magnetic resonance (NMR) spectra were acquired using a *Bruker* Ultrashield 400 Plus or *Bruker* Ascend 400 operating at 400.13, and 100.61 MHz, respectively. Data for $^{1}$H- experiments are reported as chemical shifts (relative integral, splitting, coupling constant, assignment) whilst those for $^{13}$C- are reported as chemical shifts (assignment). Chemical shifts ($\delta$) are shown in ppm. Coupling constants ($J$) are given in Hertz and the notations s, d, and t represent the multiplicities singlet, doublet and triplet, respectively. Yields from NMR were calculated by integrating the most distinct peak of each species and dividing by the total normalised integral of each species. Species identities were confirmed by spiking with solutions of authentic samples of known quantities at the same pH as the bulk sample. The increase in integral was then cross-referenced against the known amount added to provide a further method of yield quantification. Analytical High-Pressure Liquid Chromatography (HPLC) was run on Dionex Ultimate 3000 (*Thermo Scientific*) using an Atlantis T3, 3 µm, 4.6  250 mm column. Preparative High-Pressure Liquid Chromatography (HPLC) was run on a *Varian Prep Star* preparative HPLC system with a *Varian Pro Star* UV/vis detector and a *Water Atlantis* T3 Prep OBD 5 µM 19 × 250 mm column.

#### Peptide bond formation from glycyl-adenosine-5'-*O*-methylphosphate (415 mM)

Glycyl-adenosine-5'-*O*-methylphosphate (40 µL, 800 mM in water) was diluted with water (20 µL) and the pH was raised to 8.2 with NaOH (5 M, 17 µL), making a solution with a final adenylate concentration of 415 mM. The solution was kept at room temperature whilst the pH was monitored and maintained at 8.2 over 2.5 hours (as the Orgel results suggested yields plateaued within 2 hours), after which the pH was raised to 12.5 with NaOH (5 M, 17 µL) for 5 minutes at room temperature. The pH was reduced to 10 with HCl (2 M). The final concentration of adenosine-5'-*O*-methylphosphate was measured by nanodrop UV/Vis spectrometry to be 273 mM (assuming an extinction coefficient of 15000 M$^{-1}$ cm$^{-1}$). An aliquot (20 µL) was removed for functionalisation by dansyl chloride, and the remaining solution was diluted with $D_2O$ (400 µL) and analysed by NMR and dansyl labelling (Figure S1, Figure S3B).

#### Peptide bonds formation from glycyl-adenosine-5'-*O*-methylphosphate (35 mM)

Glycyl-adenosine-5'-*O*-methylphosphate (4 µL, 800 mM in water) was diluted with water (56 µL) and the pH was raised to 8.2 with NaOH (1 M, 32 µL), making a solution of adenylate with a final concentration of 35 mM. The solution was kept at room temperature whilst the pH was monitored and maintained at 8.2 over 2.5 hours, after which the pH was raised to 12.5 with NaOH (5 M, 17 µL) for 5 minutes at room temperature. The pH was reduced to 10 with HCl (2 M). The final concentration of adenosine-5'-*O*-methylphosphate was measured by nanodrop UV/vis spectrometry to be 29 mM (assuming an extinction coefficient of 15000 M$^{-1}$ cm$^{-1}$). A 20 µL aliquot was removed for functionalisation by dansyl chloride, and the remaining solution was diluted with $D_2O$ (400 µL) and analysed by NMR and dansyl labelling (Figure S2, Figure S3C).

**General method for the dansylation of crude mixtures**

An aliquot (1 µL) of crude products of peptide formation was diluted to 0.1 M. This solution (1 µL) was combined with a solution of benzylamine (1 µL, 0.1 M), then added to $NaHCO_3$ solution (40 mM, pH 9.4, 398 µL). Dansyl chloride in acetonitrile (5 mM, 200 µL) was added and the solution was kept on ice and maintained at pH 9.2 with NaOH solution (1 M). Pyridine (4%, 20 µL) was added and the solution was analysed by analytical HPLC (Buffer A: 10 mM formic acid, Buffer B: acetonitrile; 15% B at 0 min, 15% B at 1 min, 50% B at 23 min, 95% B at 33.5 min).

Yields were calculated by the integration of peaks that had the same retention time as those of the authentic synthetic standards, followed by normalisation against the integration of the *N*-dansyl-benzyl amine. This method was the product of extensive optimisation by comparison of the yields of dansylation of a mixture of glycine, diglycine and triglycine at a total concentration of 0.1 M with the integrals seen in the $^1$H NMR spectra of the same mixture of glycines (Fig. S3, Table S1).

**Peptide bonds from glycyl-adenosine-5'-*O*-methylphosphate and free glycine (both 207 mM)**

Glycyl-adenosine-5'-*O*-methylphosphate (20 µL, 800 mM in water) was diluted with water (33 µL) and combined with a solution of glycine (20 µL, 800 mM in water). The pH was raised to 8.2 with NaOH (5 M, 4 µL), making a solution with a final concentration of glycine species of 415 mM. The solution was kept at room temperature whilst the pH was monitored and maintained at 8.2 over 2.5 hours, after which the pH was raised to 12.5 with NaOH (5 M, 17 µL) for 5 minutes at room temperature. The pH was reduced to 10 with HCl (2 M). The final concentration of adenosine-5'-*O*-methylphosphate was measured by nanodrop UV/vis spectrometry to be 168 mM (assuming an extinction coefficient of 15000 $M^{-1}$ $cm^{-1}$). A 20 µL aliquot was removed for functionalisation by dansyl chloride, and the remaining solution was diluted with $D_2O$ (400 µL) and analysed by NMR and dansyl labelling (Fig. S4, Fig. S3D).

**Di-glycyl-adenosine-5'-*O*-methylphosphate hydrolysis**

Di-glycyl-adenosine-5'-*O*-methylphosphate (8 µL, 553 mM in water) was diluted with $D_2O$ (432 µL) the pD was raised to 8.2 with NaOD (1 M), making a solution of final concentration 10 mM. The solution was kept at room temperature whilst pH was monitored and maintained at 8.2 for 16 hours. Over this period the solution was periodically analysed by NMR (Fig. S5 + S6). The calculated half-life from manual fitting ($Yield = Yield_0\, e^{-0.012t}$) was 58 minutes. DKP and diglycyl adenosine were identified, as products in a 3:1 ratio.

**Peptide bond formation from formyl-glycyl- and glycyl-adenosine-5'-*O*-methylphosphate (207 mM both)**

Glycyl-adenosine-5'-*O*-methylphosphate (20 µL, 800 mM in water) was diluted with water (16.7 µL) and combined with a solution of *N*-formyl-glycyl-adenosine-5'-*O*-methylphosphate (32.3 µL, 496 mM in water). The pH was raised to 8.2 with NaOH (5 M, 8 µL), making a solution of a final concentration of 415 mM. The solution was kept at room temperature whilst the pH was monitored and maintained at 8.2 over 2.5 hours, after which the pH was raised to 12.5 with NaOH (5 M, 17 µL) for 5 minutes at room temperature. The pH was reduced to 10 with HCl (2 M). The final concentration of adenosine-5'-*O*-methylphosphate was measured by nanodrop UV/vis spectrometry to be 227 mM (assuming an extinction coefficient of 15000 $M^{-1}$ $cm^{-1}$). An aliquot (20 µL) was removed for functionalisation by dansyl chloride, and the remaining solution was diluted with $D_2O$ (400 µL) and analysed by NMR and dansyl labelling (Fig. S7, Fig. S3E).

### Synthesis

#### Adenosine-5'-*O*-methylphosphate

Adenosine-5'-*O*-monophosphate (acid form, 3.0 g, 8.64 mmol), *N,N*-di-isopropyl-ethyl amine (3.0 mL, 17.28 mmol) and DCC (9.0 g, 43.3 mmol) were dissolved in methanol (90 mL) and stirred at room temperature for 64 hours. NaOH solution (1 M, 18.9 mL) was added and then the combined solution was evaporated to dryness *in vacuo*. The solids were resuspended in water (90 mL), filtered and the solution was lyophilised. The powder was resuspended in water (90 mL), filtered and lyophilised affording adenosine-5'-(*O*-methylphosphate) 3.30 g as a white powder (quantitative yield).

**$^1$H NMR** (400 MHz, $D_2O$) $\delta$ = 8.37 (1H, s, H8), 8.18 (1H, s, H2), 6.04 (1H, d, *J* = 5.6 Hz, H1'), 4.41 (1H, dd, J = 5.6, 3.3 Hz, H3'), 4.29 (1H, p, J = 3.3 Hz, H4'), 4.03 (2H, m, H5'), 3.44 (3H d, J = 10.8 Hz, Me-H).

**$^{13}$C NMR** (100 MHz, $D_2O$) $\delta$ = 155.6 (C6), 152.8 (C2), 149.1 (C4), 118.5 (C5), 139.7 (C8), 87.8 (C1'), 83.9 (C4'), 74.9 (C2'), 70.9 (C3'), 65.1 (C5'), 52.8 (Me).

**$^{31}$P NMR** (162 MHz, $D_2O$) $\delta$ = 1.58.

**Mass**: 360.07 [M-1] calculated, 359.9 observed.

#### Glycyl-adenosine-5'-*O*-methylphosphate

*N*-Boc-glycine (175 mg, 1.00 mmol) and carbonyl diimidazole (161 mg, 1.00 mmol) were dissolved in acetonitrile (9.3 mL) and mixed for 10 minutes at room temperature. Adenosine-5'-(*O*-methylphosphate) (350 mg, 0.913 mmol) was dissolved in water (5.0 mL) and added to the acetonitrile mix. The reaction was stirred at room temperature for 10 minutes and then adjusted to pH 4 with HCl (2 M). The solution was reduced to dryness *in vacuo* and then purified by preparative HPLC using (Buffer A: 20 mM formic acid, Buffer B: acetonitrile, 0-20% over 45 minutes). Fractions containing the product were collected and lyophilized. The resultant powder was redissolved in trifluoroacetic acid (2.5 mL) and stirred for 20 minutes at room temperature before concentrating *in vacuo*. The products were purified by preparative HPLC using (Buffer A: 20 mM formic acid, Buffer B: acetonitrile, 0-20% over 45 minutes). Fractions containing the product were collected and lyophilized. Diethyl ether (10 mL × 3) was repeatedly added to the lyophilizate. The resultant suspension was sonicated and then the ether evaporated *in vacuo*. The resulting clear oil was dissolved in 150 µL water and the concentration was measured by nanodrop UV/vis spectroscopy. Assuming an extinction coefficient of 15000 $M^{-1}$ $cm^{-1}$ the concentration was measured to be 1.08 M (17% yield). a = 2' acylated, b = 3' acylated

**$^{1}$H NMR** (400 MHz, *$d_6$*-DMSO) *δ* = 8.57(1H, s, H8b), 8.53 (0.3H, s, H8a), 8.44 (2.6H, br. s, $NH_2$), 8.35 (1.3H, s, H2b), 6.95 (3.9H, brs, $NH_3^+$), 6.27 (0.3H, d, *J* = 4.8 Hz, H1'a), 6.04 (1H, d, *J* = 6.7 Hz, H1'b), 5.81 (0.3H, t, *J* = 4.8 Hz, H2'a), 5.48 (1H, dd, *J* = 5.2, 2.4 Hz, H3'b), 4.96 (1H, dd, *J* = 6.7, 5.2 Hz, H2'b), 4.60 (0.3H, t, *J* = 4.8 Hz, H3'a), 4.37 (1H, m, H4'b), 4.25-4.00 (2.9H, m, H4'a + H5'a + H5'b), 3.97 (2H, d, *J* = 4.6, αb), 3.91 (0.6H m, αa), 3.52 (3H d, *J* = 10.5 Hz, Me-H), 3.50 (0.9H d, *J* = 10.5 Hz, Me-H).

**$^{13}$C NMR** (100 MHz, *$d_6$*-DMSO) *δ* = 166.9 (ester), 153.9 (C6), 149.9 + 149.6 + 149.3 + 148.9 (C2 + C4), 140.6 + 140.3 (C8), 118.9 (C5), 86.7 (C1'b), 85.0 (C1'a), 83.3 (C4'a), 81.1 (C4'b), 76.3 (C2'a), 74.5 (C3'b), 72.2 (C2'b), 68.9 (C3'a), 65.3 (C5'), 53.1 (Me), 40.1 (α).

**$^{31}$P NMR** (162 MHz, *$d_6$*-DMSO) *δ* = -0.37.

**Mass**: 417.09 [M-1] calculated, 416.9 observed.

**Di-glycyl-adenosine-5'-*O*-methylphosphate**

*N*-Boc-diglycine (107 mg, 0.46 mmol) and carbonyl diimidazole (74 mg, 0.46 mmol) were dissolved in acetonitrile (4.6 mL) and mixed for 10 minutes at room temperature. Adenosine-5'-*O*-methylphosphate (80.5 mg, 0.42 mmol) was dissolved in water (2.5 mL) and added to the acetonitrile mix. The reaction was stirred at room temperature for 10 minutes and then adjusted to pH 4 with HCl (2 M). The solution was reduced to dryness *in vacuo* and then purified by preparative HPLC. Fractions containing the product were collected and lyophilized. The resultant powder was redissolved in trifluoroacetic acid (2.50 mL) and stirred for 20 minutes at room temperature before concentrating *in vacuo*. The products were purified by preparative HPLC. Fractions containing the product were collected and lyophilized. Diethyl ether (3 × 10 mL) was repeatedly added to the lyophilizate. The resultant suspension was sonicated and then the ether evaporated *in vacuo*. The resulting clear oil was dissolved in 56 µL water and the concentration was measured by nanodrop UV/vis spectroscopy. Assuming an extinction coefficient of 15000 $M^{-1}$ $cm^{-1}$ the concentration was measured to be 553 mM (7% yield). a = 2' acylated, b = 3' acylated

**$^{1}$H NMR** (400 MHz, *$d_6$*-DMSO) *δ* = 8.91 (1H, t, *J* = 5.4, amide-Hb), 8.80 (0.3H, t, *J* = 6.3, amide-Ha), 8.71 (0.3H, t, *J* = 5.5, DKP-amide), 8.46 (1H, s, H8b), 8.42 (0.3H, s, H8a), 8.25 (1H, s, H2b), 8.24 (0.3H, s, H2a), 8.14 (3.9H, m, $NH_3^+$), 7.81 (2H, br. s, $NH_2$b), 7.76 (0.6H, br. s, $NH_2$a), 6.21 (0.3H, d, *J* = 4.8 Hz, H1'a), 5.98 (1H, d, *J* = 6.7 Hz, H1'b), 5.74 (0.3H, t, *J* = 4.8 Hz, H2'a), 5.37 (1H, dd, *J* = 5.3, 2.9 Hz, H3'b), 4.93 (1H, dd, *J* = 6.7, 5.3 Hz, H2'b), 4.56 (0.3H, t, *J* = 4.8 Hz, H3'a), 4.32 (1H, m, H4'b), 4.18-4.00 (5.2H, m, H4'a, H5'a, H5'b, internal-αb, DKP-α), 3.87 (0.6H, d, *J* = 5.5, internal-αa), 3.67 (2H brs, terminal- αb), 3.62 (0.6H brs, terminal-αa), 3.52 (3H d, *J* = 11.0 Hz, Me-H), 3.51 (0.9H d, *J* = 11.0Hz, Me-H).

**$^{13}$C NMR** (100 MHz, $d_6$-DMSO) $\delta$ = 171.2 (amide), 169.1, 167.0 (ester), 155.5 (C6), 151.8 (C2), 149.7 (C4), 140.2 (C8), 119.4 (C5), 87.3 (C1'b), 85.7 (C1'a), 83.4 (C4'a), 81.2 (C4'b), 75.9 (C2'a), 73.9 (C3'b), 72.2 (C2'b), 69.1 (C3'a), 65.5 (C5'), 53.3 (Me), 40.7, 40.1 ($\alpha$).

**$^{31}$P NMR** (162 MHz, $d_6$-DMSO) $\delta$ = -0.44.

**Mass**: 474.09 [M-1] calculated, 473.9 observed.

#### ***N*-Formyl-glycyl-adenosine-5'-*O*-methylphosphate**

*N*-Formyl-glycine (69.0 mg, 0.66 mmol) and carbonyl diimidazole (106 mg, 0.66 mmol) were dissolved in acetonitrile (6 mL) and mixed for 10 minutes at room temperature. Adenosine-5'-*O*-methylphosphate (210 mg, 0.548 mmol) was dissolved in water (3 mL) and added to the acetonitrile mix. The reaction was stirred at room temperature for 10 minutes and then adjusted to pH 4 with HCl (2 M). The solution was reduced to dryness *in vacuo* and then purified by preparative HPLC. Fractions containing the product were collected and lyophilized. Diethyl ether (3 × 10 mL) was repeatedly added to the lyophilizate. The resulting suspension was sonicated and then the ether evaporated *in vacuo*. The resulting clear oil was dissolved in 170 μL water and the concentration was measured by nanodrop UV/vis spectroscopy. Assuming an extinction coefficient of 15000 $M^{-1}$ $cm^{-1}$ the concentration was measured to be 813 mM (24% yield). a = 3' acylated, b = 2' acylated

**$^{1}$H NMR** (400 MHz, $d_6$-DMSO) $\delta$ = 8.62 + 8.54 (1H, t, $J$ = 6.0, amide), 8.51 (1H, s, H8), 8.44 (1H, s, H8), 8.17 + 8.15 (2H + 2H, s, H2 + Formyl-H), 7.34 (2H, br. s, $NH_2$), 7.30 (2H, br. s, $NH_2$), 5.96 (1H, d, $J$ = 7.1 Hz, H1'a), 5.93 (1H, d, $J$ = 5.5 Hz, H1'b), 5.38 (1H, dd, $J$ = 5.3, 1.5 Hz, H3'a), 4.95 (1H, dd, $J$ = 7.1, 5.3 Hz, H2'a), 4.60 (1H, t, $J$ = 5.5 Hz, H2'b), 4.23 (1H, m, H4'a), 4.19 (1H, t, $J$ = 4.1 Hz, H3'b), 4.05 (4H, s, H$\alpha$ + H4'b), 3.78-3.98 (6H, m, H$\alpha$ + H5'a + H5'b), 3.37 (3H d, $J$ = 10.3 Hz, Me-H), 3.35 (3H d, $J$ = 10.6 Hz, Me-H).

**$^{13}$C NMR** (100 MHz, $d_6$-DMSO) $\delta$ = 169.3 (ester), 161.9 (amide), 156.4 (C6), 153.1 (C2), 150.2 (C4), 139.3 (C8), 119.3 (C5), 87.0 (C1'), 86.3 (C1'), 84.5 (C4'b), 81.6 (C4'a), 74.3 (C3'a), 73.9 (C2'b), 72.2 (C'2a), 70.9 (C3'b), 64.8 (C5'), 64.5 (C5'), 51.8 (Me), 39.5 ($\alpha$).

**$^{31}$P NMR** (162 MHz, $d_6$-DMSO) $\delta$ = -0.3.

**Mass**: 445.1 [M-1] calculated, 444.9 observed.

#### ***N*-formyl-diglycine**

Diglycine (500 mg, 3.78 mmol) was dissolved in neat formic acid (8 mL) and heated to 55 °C. Acetic anhydride (3.0 mL) was added dropwise over 5 minutes. The reaction was allowed to cool to room temperature whilst stirring for 1 hour. Ice was added and the sample was kept at 4 °C overnight. The sample was then lyophilised affording 495 mg of product (82% yield).

**$^{1}$H NMR** (400 MHz, $D_2O$) $\delta$ = 8.15 (1H, s, formyl-H), 3.99 (2H, s, $\alpha$), 3.93 (2H, s, $\alpha$);

**$^{13}$C NMR** (100 MHz, $D_2O$) $\delta$ = 176.6, 170.9, 165.0, 43.4, 41.2.

**Mass**: 159.0 [M-1] calculated, 159.0 observed.

***N*-formyl-triglycine**

Triglycine (140 mg, 0.74 mmol) was dissolved in neat formic acid (2.0 mL) and heated to 55 °C. Acetic anhydride (0.75 mL) was added dropwise over 5 minutes. The reaction was allowed to cool to room temperature whilst stirring for 1 hour. Ice was added and the sample was kept at 4 °C overnight. The sample was then lyophilised affording 159.5 mg of product (quantitative yield).

**$^1$H NMR** (400 MHz, $D_2O$, pD 10.1) $\delta$ = 8.14 (1H, s, formyl-H), 3.99 (2H, s, $\alpha$), 3.93 (2H, s, $\alpha$), 3.71 (1H, s, $\alpha$);

**$^{13}$C NMR** (100 MHz, $D_2O$) $\delta$ = 176.5, 171.8, 171.2, 165.1, 43.3, 42.5, 41.3.

**Mass**: 216.1 [M-1] calculated, 216.0 observed.

***N*-Dansyl-glycine**

Glycine (30.1 mg, 0.408 mmol) in NaOH aq. (1 M, 0.5 mL) was added to dansyl chloride (100 mg, 0.371 mmol) in THF (3.5 mL) and stirred for 1 hour at room temperature. During that time the pH was monitored and kept above 9 with additional additions of 1 M NaOH. The solution was washed with ethyl acetate (3 × 10 mL). The aqueous layer was acidified to pH 4 with 1 M HCl and then washed with ethyl acetate (3 × 10 mL). The combined acidic organic layers were dried over anhydrous magnesium sulphate and then concentrated *in vacuo* affording 2 mg of *N*-dansyl-glycine (1.4% unoptimised yield – sufficient to provide an authentic product standard).

**$^1$H NMR** (400 MHz, $d_6$-DMSO) $\delta$ = 8.44 (1H, d, $J$ = 8.8 Hz,), 8.29 (1H, d, $J$ = 8.8 Hz), 8.10 (1H, d, $J$ = 7.1), 7.58 (2H, m), 7.24 (1H, d, $J$ = 7.5), 3.59 (2H, s, $\alpha$), 2.82 (6H, s, Me).

**$^{13}$C NMR** (100 MHz, $d_6$-DMSO) $\delta$ = 170.3 (Carboxylic acid), 151.3 (N-Ar), 136.4, 129.3, 129.1, 127.8, 127.8, 123.5, 119.4, 115.1, 45.1 (Me), 43.8 ($\alpha$).

**Mass**: 307.1 [M-1] calculated, 307.0 observed.

***N*-Dansyl-diglycine**

Diglycine (53.9 mg, 0.408 mmol) in NaOH aq. (1 M, 0.5 mL) was added to dansyl chloride (100 mg, 0.371 mmol) in THF (3.5 mL) and stirred for 1 hour at room temperature. During that time the pH was monitored and kept above 9 with additional additions of 1 M NaOH. The solution was washed with ethyl acetate (3 × 10 mL). The aqueous layer was acidified to pH 4 with 1 M HCl and then washed with ethyl acetate (3 × 10 mL). The combined acidic organic layers were dried over anhydrous magnesium sulphate and then concentrated *in vacuo* affording 18.8 mg of *N*-dansyl-diglycine (13% yield).

**$^1$H NMR** (400 MHz, $d_6$-DMSO) $\delta$ = 8.46 (1H, d, $J$ = 8.8 Hz), 8.29 (2H, d, $J$ = 8.8 Hz), 8.09 (3H, m), 7.58 (2H, q, $J$ = 8.0), 7.26 (1H, d, $J$ = 7.8), 3.68 (2H, d, $J$ = 5.6, 3.50$\alpha$), (2H, s, $\alpha$), 2.83 (6H, s, Me).

**$^{13}$C NMR** (100 MHz, $d_6$-DMSO) $\delta$ = 170.9 (carboxylic acid), 168.0 (amide), 151.2 (N-Ar), 135.9, 129.4, 129.0, 128.0, 127.9, 123.5, 119.2, 115.1, 45.1 (Me), 44.9 ($\alpha$), 40.6 (terminal-$\alpha$).

**Mass**: 364.1 [M-1] calculated, 364.0 observed.

***N*-Dansyl-triglycine**

Triglycine (77.2 mg, 0.408 mmol) in NaOH aq. (1 M, 0.5 mL) was added to dansyl chloride (100 mg, 0.371 mmol) in THF (3.5 mL) and stirred for 1 hour at room temperature. During that time the pH was monitored and kept above 9 with additional additions of 1 M NaOH. The solution was washed with ethyl acetate (3 × 10 mL). The aqueous layer was acidified to pH 4 with 1 M HCl and then washed with ethyl acetate (3 × 10 mL). The combined acidic organic layers were dried over anhydrous magnesium sulphate and then concentrated *in vacuo* affording 18.4 mg of *N*-dansyl-triglycine (11% yield).

**$^1$H NMR** (400 MHz, $d_6$-DMSO) $\delta$ = 8.46 (1H, d, $J$ = 8.4 Hz,), 8.29 (1H, d, $J$ = 8.8 Hz), 8.11 (4H, m), 7.60 (2H, m), 7.26 (1H, d, $J$ = 7.5), 3.74 (2H, d, $J$ = 5.9, $\alpha$), 3.66 (2H, d, $J$ = 5.7, $\alpha$), 3.50 (2H, s, $\alpha$), 2.83 (6H, s, Me).

**$^{13}$C NMR** (100 MHz, $d_6$-DMSO) $\delta$ = 171.0, 168.8, 167.9 (amide, carboxylic acid), 151.5 (N-Ar), 136.4, 129.5, 129.1, 128.2, 127.9, 123.5, 119.2, 115.2, 45.1 (Me), 44.7 ($\alpha$), 41.7 ($\alpha$), 40.2 ($\alpha$).

**Mass**: 421.1 [M-1] calculated, 421.0 observed.

**Solid phase synthesis of RNA oligomers**

All biotin-labelled RNA oligomers were purchased from *Sigma-Aldrich* and used without further purity checks. All other RNA oligomers were synthesized using a *GE Healthcare* ÄKTA Oligopilot plus 10 on a 20 to 50 µmol scale. After automated synthesis, RNAs were first cleaved from the solid support by treating with 3 mL of a 1:1 mixture of $NH_3$ aqueous solution (28% wt) and $CH_3NH_2$ ethanol solution (33% wt) at 55°C for 80 minutes in a tube with a sealed cap. The solid was removed by filtration and washed with 50% EtOH/$H_2O$. The solutions were combined and evaporated to dryness under reduced pressure. Silyl protecting groups were removed by treating the residues with 2 mL of a 1:1 (v/v) mixture of triethylamine trihydrofluoride and DMSO at 65 °C for 150 minutes in a tube with a sealed cap. After brief cooling at -32 °C, 40 mL of cold 50 mM $NaClO_4$ in acetone was added to the solution to precipitate the oligoribonucleotides. The resulting mixture was centrifuged, purified using preparative HPLC, checked using analytical HPLC, and lyophilized. The purified RNA was redissolved in 2 mL of water and passed through a *Waters* Sep-Pak C18 Cartridge with 10 g sorbent. Eluates were checked using analytical HPLC, combined, and lyophilized. The resulting white powder was stored at -32 °C for future use.

Certain anticodon loops and ssRNA are not accessible due to self-dimerization and G-quadruplex formation. Refer to Fig. S15 for a detailed discussion. The anticodon loop and ssRNA may potentially form a four-base pairing complex. See Fig. S16 for details.

**Isothermal titration calorimetry experiment for anticodon loop(s) and ssRNA binding**

Isothermal titration calorimetry (ITC) experiments were performed on a *Malvern* MicroCal iTC 200 at 10 °C. Anticodon loops and tri-/hexamer ssRNA were dissolved in binding buffer (50 mM HEPES pH 7.2, 100 mM NaCl, 100 mM $MgCl_2$). In a typical experiment, an anticodon loop solution (800 µM, 42 µL) was injected 28 times at 200-second intervals from a syringe into the sample cell containing ssRNA (40 μM, 350 μL) at a stirring speed of 750 RPM. The first injection was 0.5 μL of anticodon loop solution, and the subsequent 27 injections were 1.5 μL each time. The control experiment was conducted by titrating the anticodon loop with the blank buffer. The data were analyzed by MicroCal PEAQ-ITC and plotted using GraphPad Prism.

**Biolayer interferometry experiment for anticodon loop and ssRNA binding**

Biolayer interferometry (BLI) experiments were performed on a *FortéBio* Octet RED384 system with streptavidin biosensors at 20 °C. The biotin-labelled ssRNA (0.9 μM) was diluted in the same buffer as used in the ITC experiments, i.e. 50 mM HEPES pH 7.2, 100 mM NaCl, 100 mM $MgCl_2$, and bound to the streptavidin biosensor for 180 s, followed by a washing step (60 s) with ITC buffer, an association step (180 s or 300 s) of the anticodon loops (0 to ca. 800 μM) to the sensor-bound ssRNA, and a dissociation step (180 s or 300 s) in the buffer. The assay was performed in a solid black 96-/384-well plate, using an agitation set at 1000 rpm. The control experiment was conducted by dipping the biosensor with ssRNA into the blank buffer. The data were analyzed by Data Analysis HT V11.0 and plotted using GraphPad Prism.

**Discussion on exceptions involving family-box codons with less optimal binding affinity**

We observed that 12 out of 32 family-box codons exhibited $K_D$ values larger than 130 µM. These specific codons are highlighted in red in the table below.

| BLI | U | C | A | G | |
|---|---|---|---|---|---|
| U | / | 260 | / | / | U |
| | 450 | 75 | 1500 | / | C |
| | / | 280 | / | / | A |
| | 240 | 310 | 210 | 370 | G |
| C | 150 | 230 | 510 | 75 | U |
| | 65 | 14 | 230 | 90 | C |
| | / | 75 | / | 36 | A |
| | 730 | 75 | 270 | 85 | G[a] |
| A[d] | / | 530 | / | 170 | U[d] |
| | 1000 | 27 | 860 | 180 | C |
| | 420 | 150 | / | 1000 | A |
| | 330 | 130 | 1400 | 85 | G |
| G | 470 | 120 | 190 | 115 | U |
| | 35 | 17 | 230 | 60 | C[a] |
| | 400 | 95 | 780 | 130 | A |
| | 160 | 38 | 140 | 35 | G |

We attribute the suboptimal binding affinities to two factors:

1) The solubility of the anticodon loops is compromised, particularly in cases where loops contain multiple purines. These loops, which pair with UCU, CUU, CCU, ACU, and GUU, are less soluble in the aqueous buffer.

2) Certain loops have a propensity to form self-dimers in our model, thereby reducing the availability of free anticodon loops in solution (Fig. S15). Consequently, this leads to an apparent decrease in binding affinity, as observed with loops pairing to UCA, CUG, and ACU.

The $K_D$ of the codon AGG binding to its corresponding loops exhibits a low value of 85 µM (highlighted in blue), equivalent to the $K_D$ of CGG. We maintain that even though AGG may form stable sequential bindings on a ssRNA, it does not alter our hypothesis that "family-box codons bind tighter." Arginine is designated to AGG and CGN boxes, and all CGN codons display strong binding to their anticodon loops. Consequently, arginine can undergo transpeptidation with both CGN and AGG codon-anticodon loop interactions.

**Fig. S1** Stacked [1]H-NMR spectra showing the peptidyl products of the high concentration reaction of glycyl-adenosine-5'-*O*-methylphosphate (415 mM). Identities were confirmed by serial spiking of triglycine, diglycine and DKP, whilst glycine was identified by its characteristic peak shift. Yields of 6% diglycine, 12% diketopiperazine (DKP), and < 0.5% triglycine were observed.

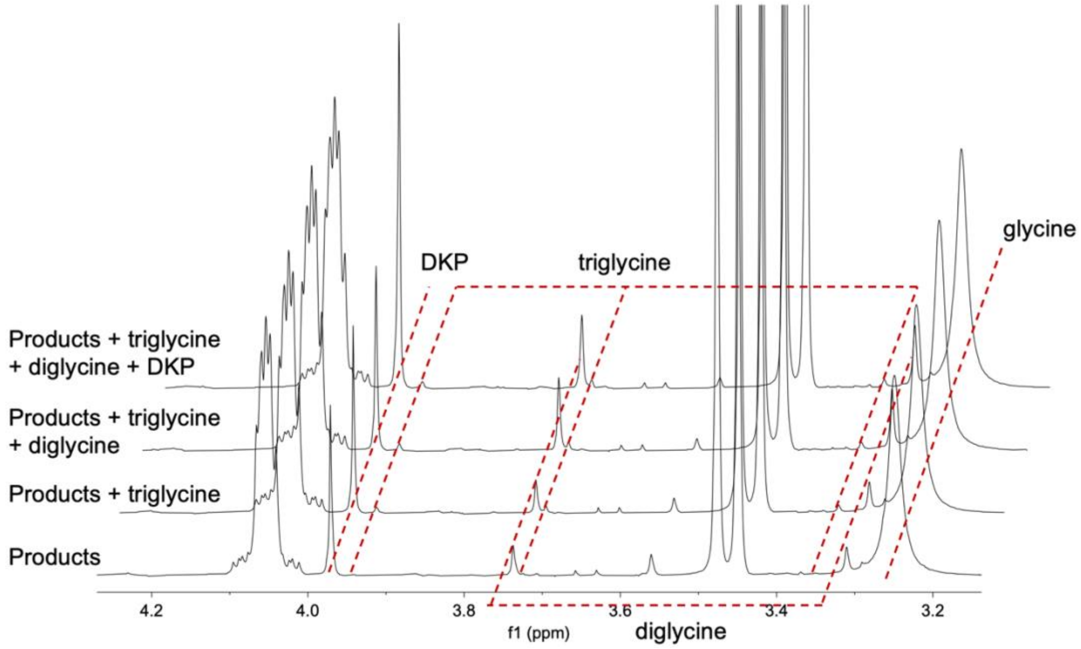

**Fig. S2** Stacked [1]H-NMR spectra showing the peptidyl products of the high concentration reaction of free glycine and glycyl-adenosine-5'-*O*-methylphosphate (both 207 mM). Identities were confirmed by serial spiking of triglycine, diglycine and DKP, whilst glycine was identified by its characteristic peak shift. Yields of 3.4% diglycine, 3% diketopiperazine (DKP), and < 0.1% triglycine were observed.

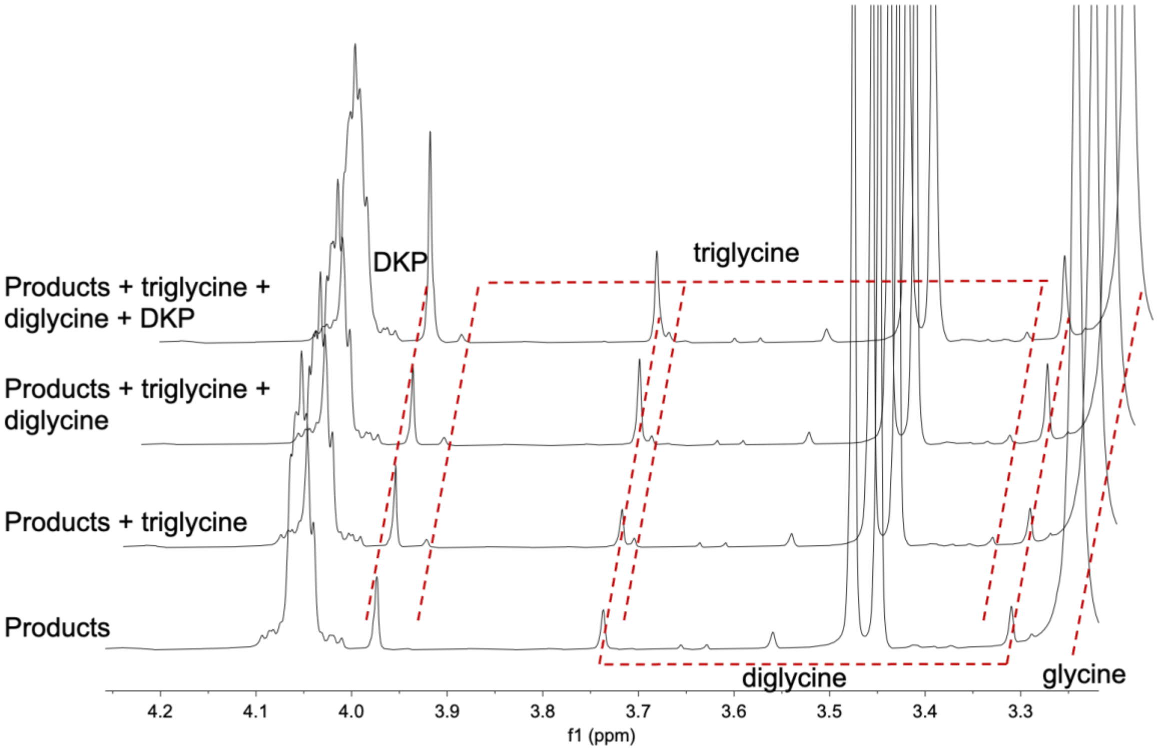

**Fig. S3 HPLC chromatograms of the products of peptide bond formation after dansylation.** Top: 11.3 – 39.6 min. Bottom: 14.96 – 20.72 min at increased intentisty. A) Standard mix of glycine, diglycine and triglycine; B) products of glycyl-adenosine-5'-*O*-methylphosphate at 415 mM (refer to Fig. S1); C) products of glycyl-adenosine-5'-*O*-methylphosphate and free glycine at 207 mM (refer to Fig. S2); D) products of glycyl-adenosine-5'-*O*-methylphosphate at 35 mM (refer to Fig. S4); E) products of glycyl-adenosine-5'-*O*-methylphosphate and formyl-glycyl-adenosine-5'-*O*-methylphosphate at 207 mM (refer to Fig. S7). Detection wavelength: 300 nm.

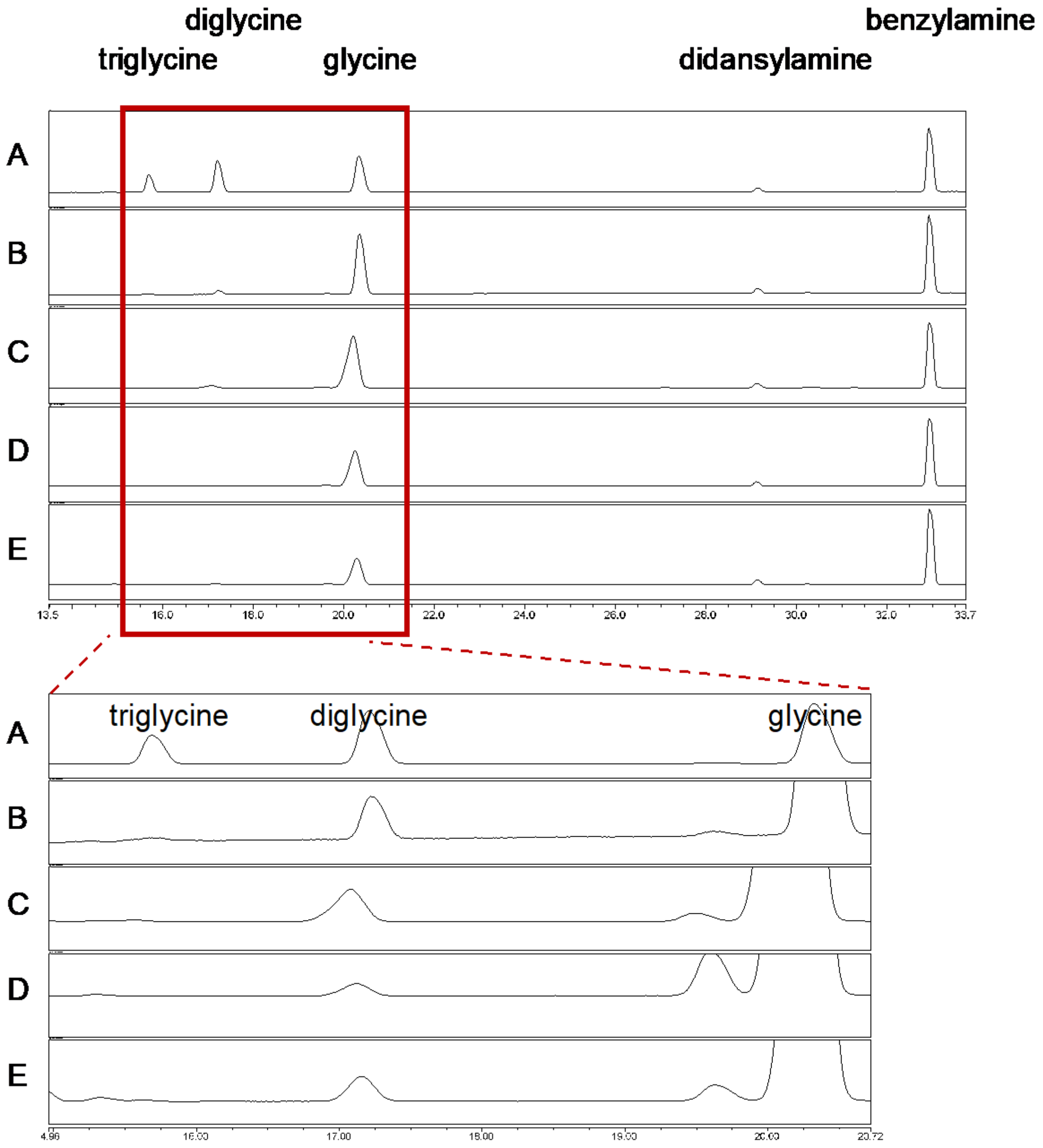

**Fig. S4** Stacked $^{1}$H-NMR spectra showing the peptidyl products of the low concentration reaction of glycyl-adenosine-5'-*O*-methylphosphate (35 mM). Identities were confirmed by serial spiking of diglycine and DKP, whilst glycine was identified by its characteristic peak shift. Traces of DKP, but no diglycine were observed.

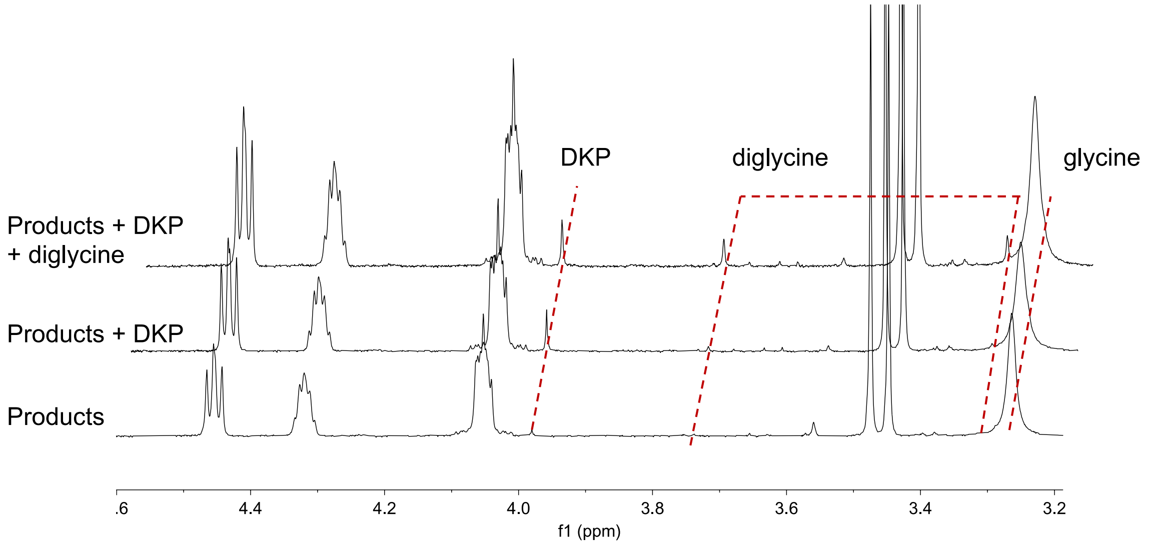

**Fig. S5** Yields and kinetic date of the hydrolysis of diglycyl-adenosine-5'-*O*-methylphosphate. Conditions: adenylate (10 mM), pH = 8.2, 25 °C.

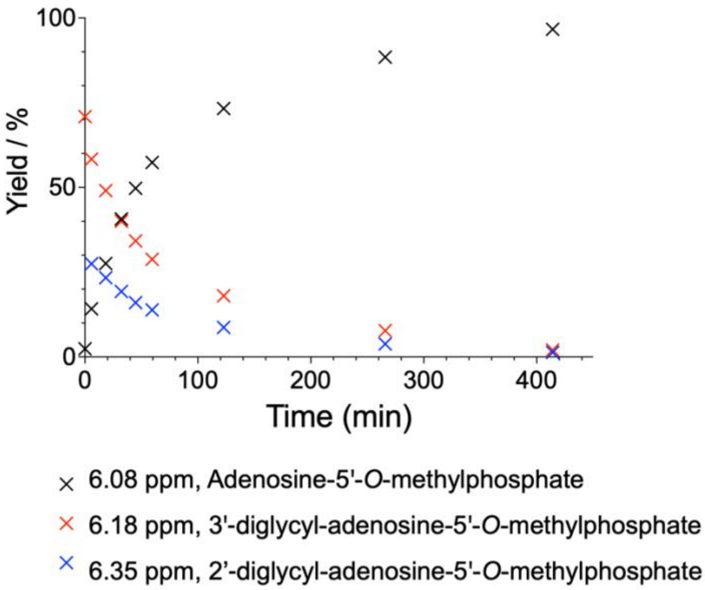

**Fig. S6** Stacked $^{1}$H-NMR spectra between 5.15 – 6.60 ppm (top) and 3.25 – 4.55 ppm (bottom) for the half life of hydrolysis of diglycyl-adenosine-5'-*O*-methylphosphate. After 16 hours the ratio of DKP and diglycine was calculated from the normalised integration of 3.98 ppm (DKP) and 3.75 ppm (diglycine) to be 3:1.

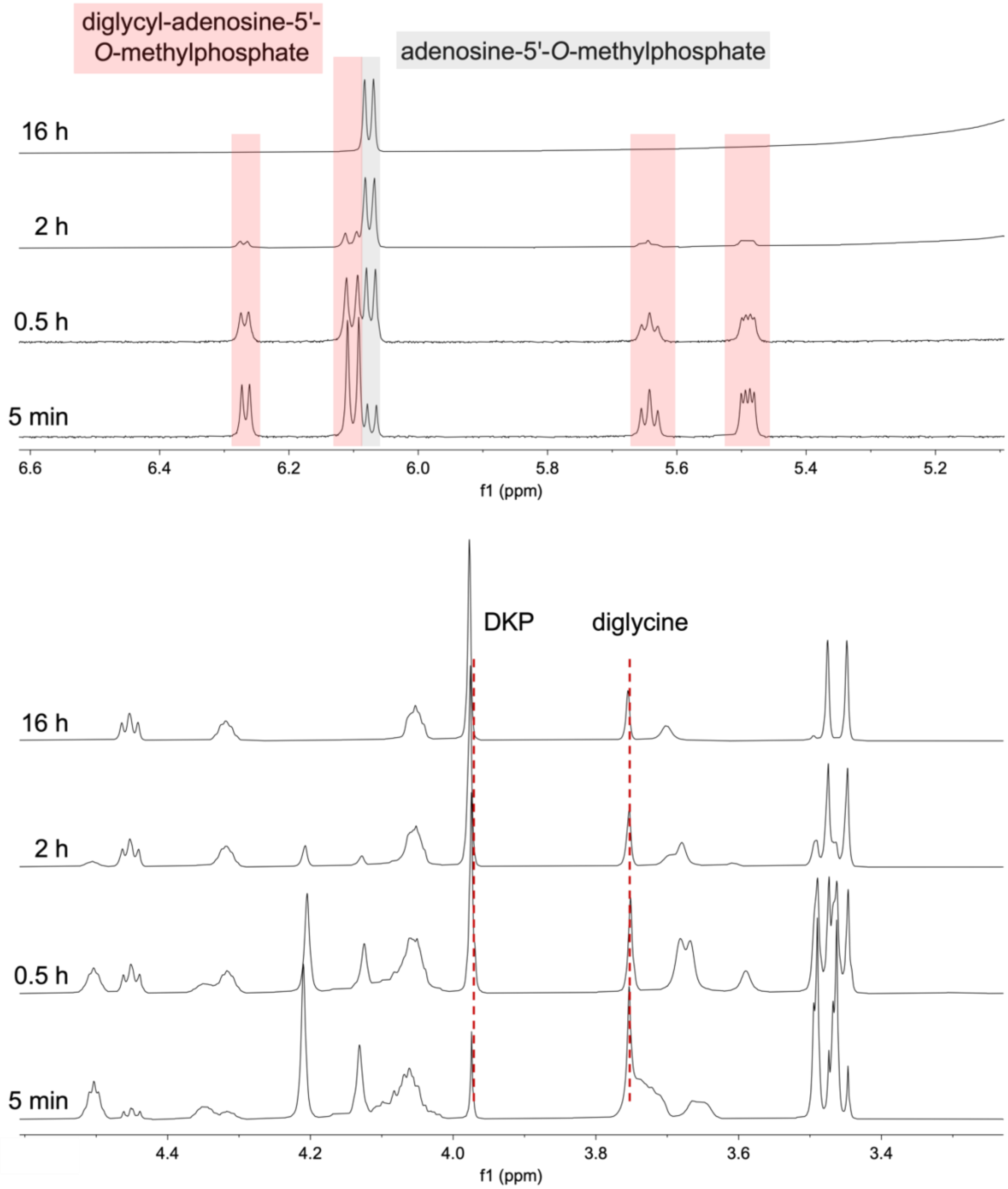

**Fig. S7** Stacked $^1$H-NMR spectra showing the peptidyl products of the high concentration reaction of formyl-glycyl-adenosine-5'-*O*-methylphosphate (207 mM) and free glycyl-adenosine-5'-*O*-methylphosphate (207 mM). Identities were confirmed by serial spiking of triglycine, diglycine and DKP, whilst glycine was identified by its characteristic peak shift. Yield of 1.5% diglycine, 5.4% *N*-formyl-digly, 5.2% DKP, ~0.1% tripeptides are observed. F-, formyl group.

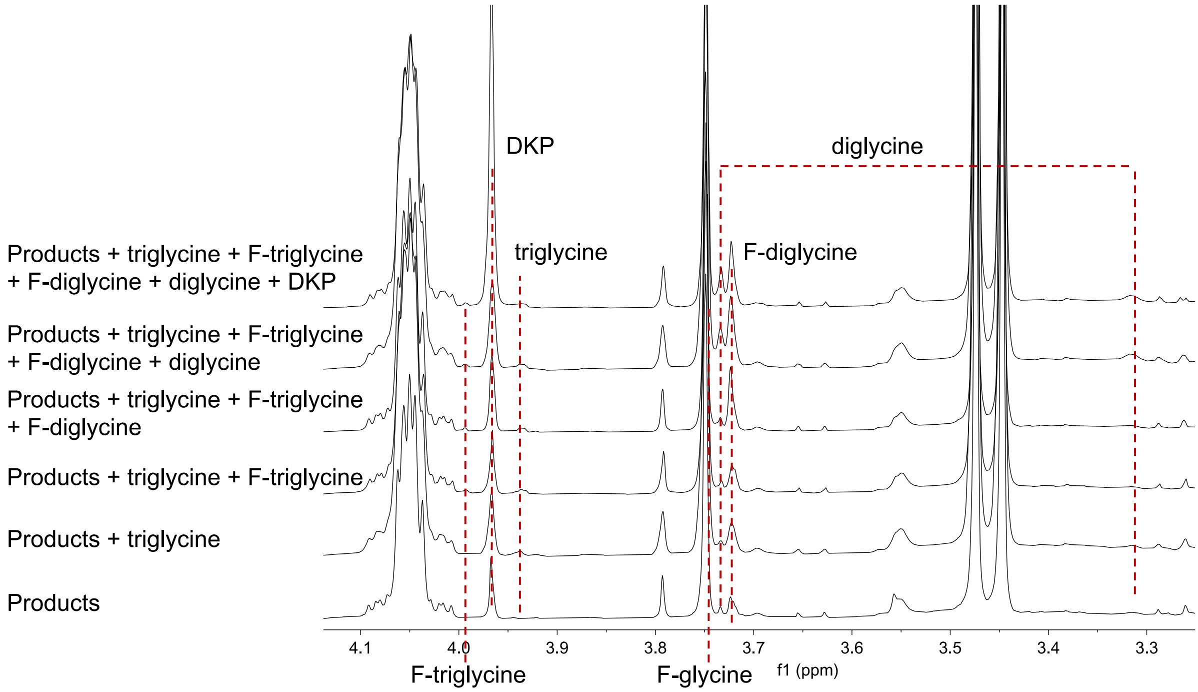

**Fig. S8 Isothermal titration calorimetry curves of anticodon loops with varied residue at positions 33, 35, and 37 (consensus tRNA numbering).** It is documented that the 2’-hydroxyl group of U33 forms a hydrogen bond with N7 of A35 (A). In the case of G35, a bifurcated hydrogen bond bridges the (U33)O2'-H and guanine N7 and O6. The hydrogen bond is less obvious when residue 35 is a pyrimidine (C or U). Titration of the loop 5'-CUGGUAA to codon 5'-ACC revealed a $K_D$ of 5.7 µM (B). Replacing the second G of the anticodon to C abrogated the binding (C). A parallel example showed that loop 5'-CUCGCAA bound to 5'-GCG with a $K_D$ of 1.1 µM. Altering the U33 to A decreased the affinity to 6.0 µM (D,E). These results confirmed that U33:R35 hydrogen bond enhances the binding of the anticodon loop to ssRNA. In addition, A37 contributes towards the tight binding through base stacking as shown by the reduction in binding when it is changed to C (F,G). Conditions: 100 mM NaCl, 100 mM $MgCl_2$, 50 mM HEPES, pH 7.2, 10 ºC.

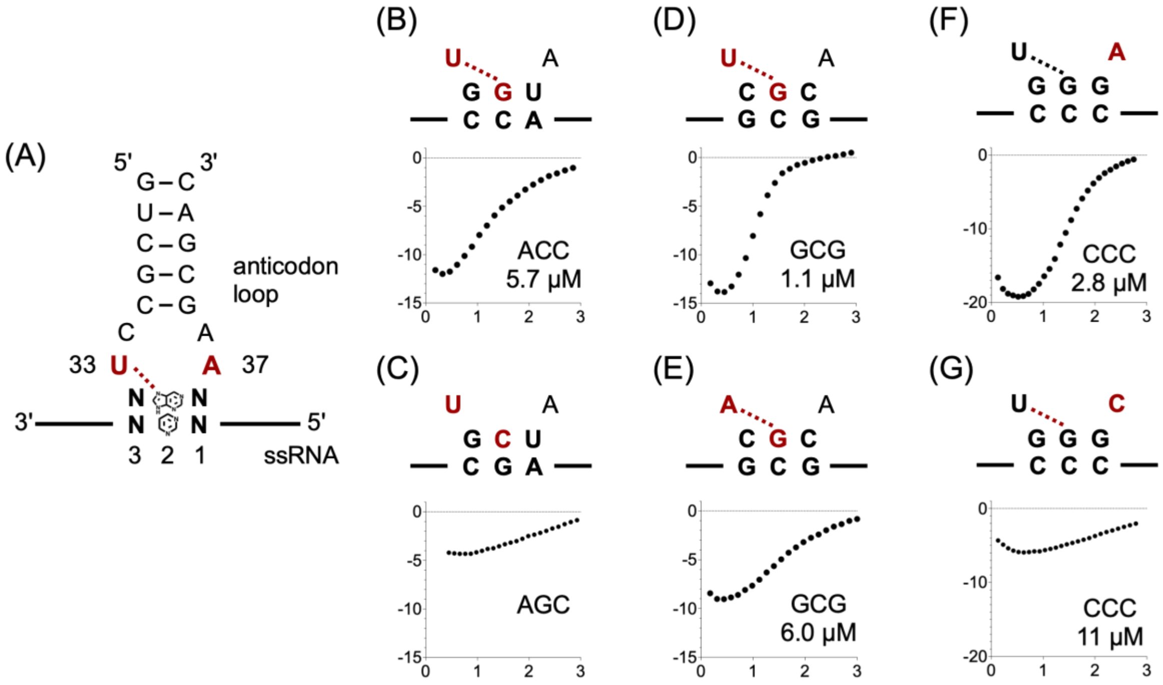

**Fig. S9 Measurement of anticodon loop:codon binding affinity using BLI.** (A) Schematic representation of an anticodon loop binds to a biotin-labelled ssRNA containing the respective codon; (B) typical BLI binding curves of four codons, two family-box codons in black (top, CACCC and AGUCA), and two split-box codons in red (below, CCAUU and AUUCC). Conditions: 0–800 μM each anticodon loop, 100 mM NaCl, 100 mM $MgCl_2$, 50 mM HEPES, pH 7.2, 20 °C.

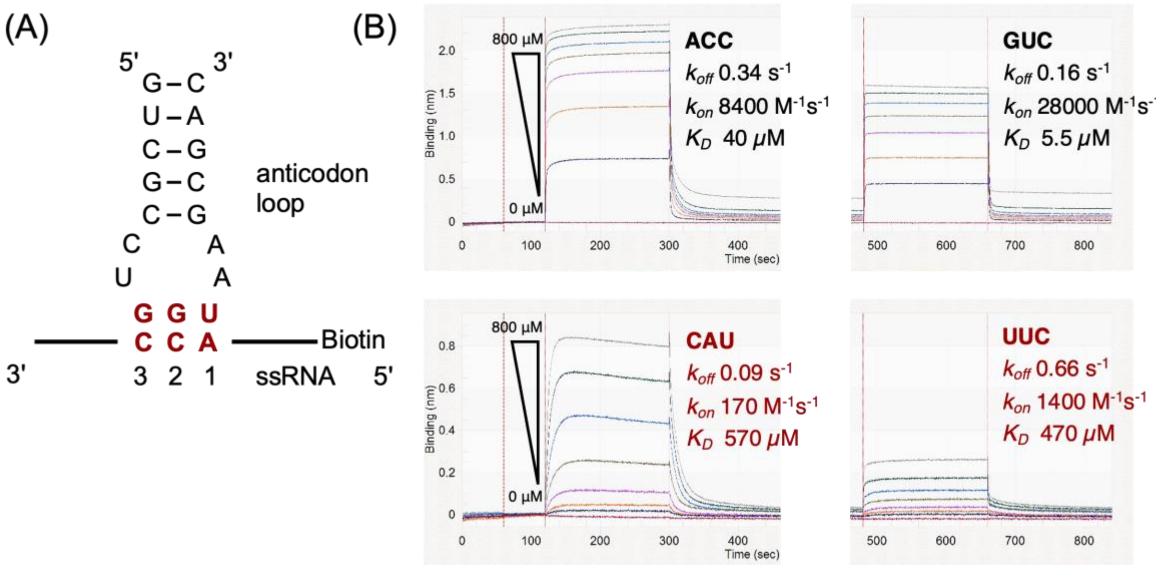

**Fig. S10** Box and Whisker plots of (A) equilibrium dissociation constants, (B) association constants, and (C) disocation constants of measured anticodon loop:codon ending with A/C/G/U in two codon classes (family and split boxes). The plot offset on the right shows comparasion for all family and split box codons. P-value of permutation test for Pearson correlation coefficient is displayed. For family-box codons: NNA, n = 6; NNC, n = 8; NNG, n = 6; NNU, n =8. For split-box codons: NNA, n = 3; NNC, n = 7; NNG, n = 8; NNU, n = 8. Sequences with an n value less than 8 result from certain sequences exhibiting a binding affinity weaker than detectable by BLI ($K_D$ > 3000).

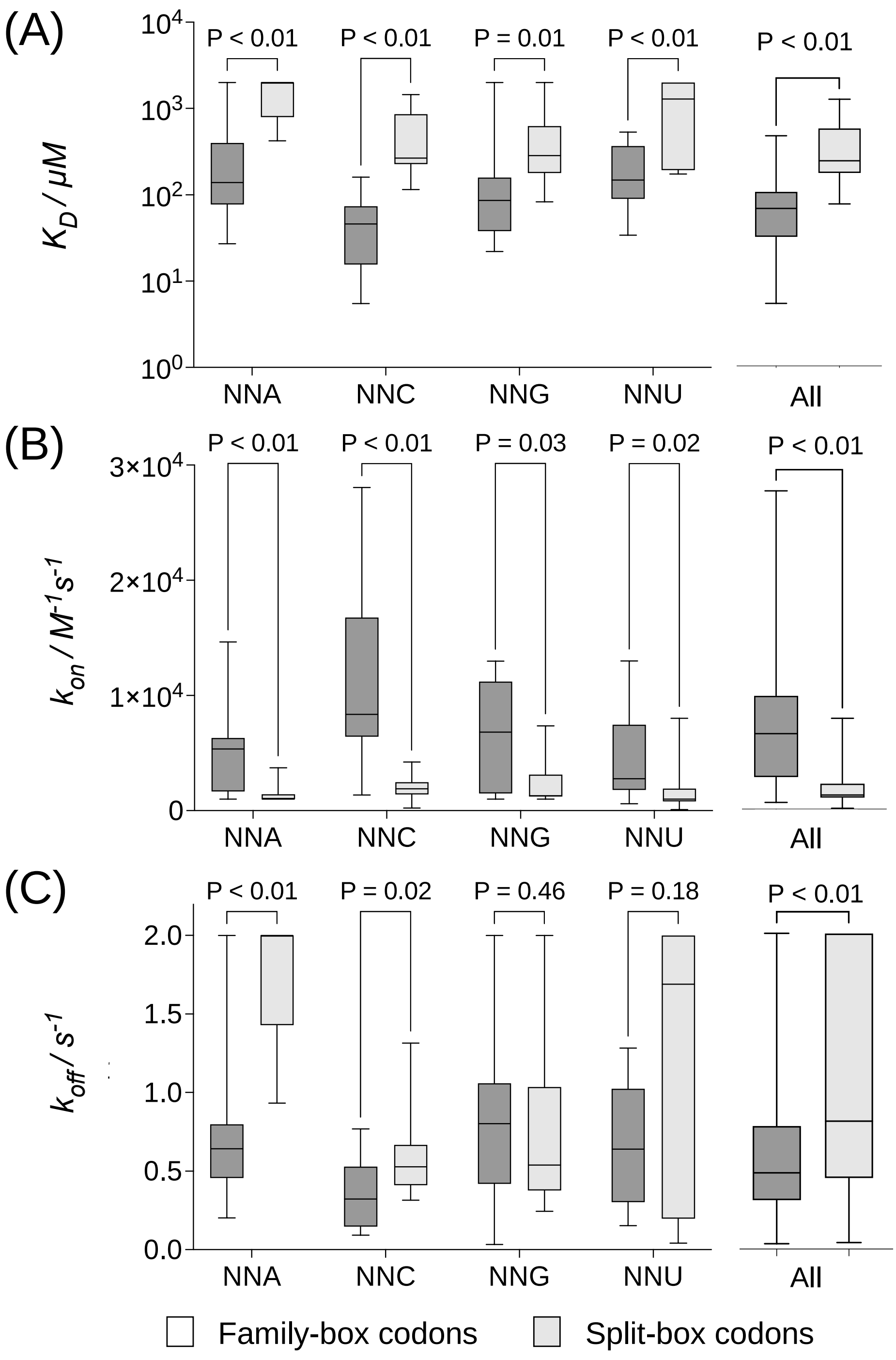

**Fig. S11** Isothermal titration calorimetry curves of anticodon loops with A/C/G/U-ending codons binding to a ssRNA containing either a single triplet (light grey or red) or two continguous identical triplets (black or red). A) Codons ending in A; B) Codons ending C; C) Codons ending in G; D) Codons ending in U. Black curves represent family-box codons; red curves represent split-box codons. Empty circles in GCC represent an exothermic process due to ssRNA self-dimer and are excluded from the calculation. Titration of hexamer guanine is absent due to the self-aggregation of oligo guanine in the excluded buffer. Conditions: 800 µM anticodon loop, 40 µM ssRNA, 100 mM NaCl, 100 mM $MgCl_2$, 50 mM HEPES, pH 7.2, 10 ºC. Numerical values for $K_{D,}$ molar ratio (R), and enthalpy change (kcal/mol) can be found in tables S8 and S9.

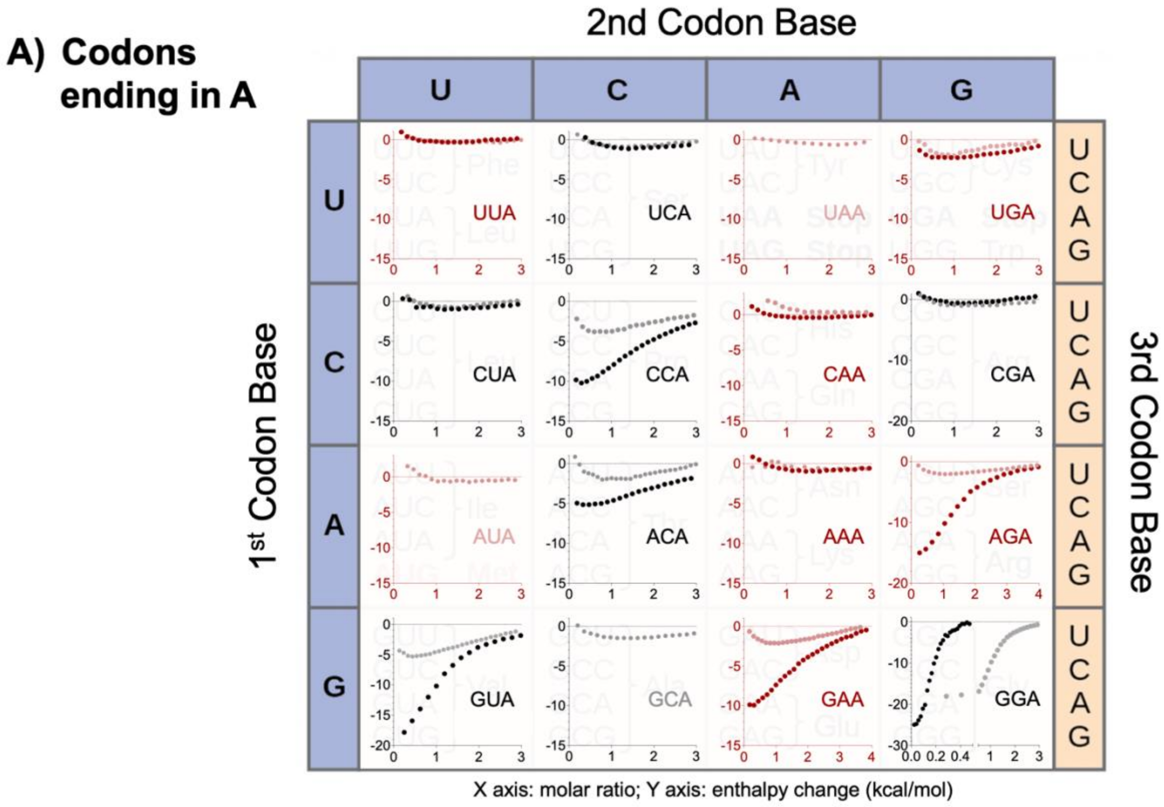

**B) Codons ending in C**

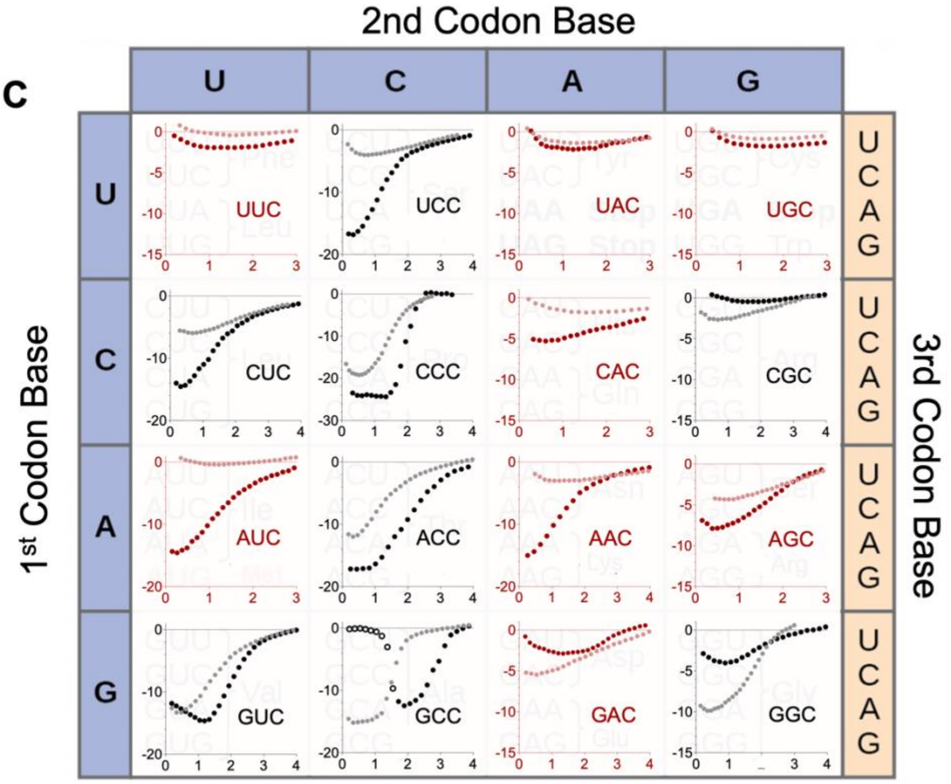


X axis: molar ratio; Y axis: enthalpy change (kcal/mol)

**C) Codons ending in G**

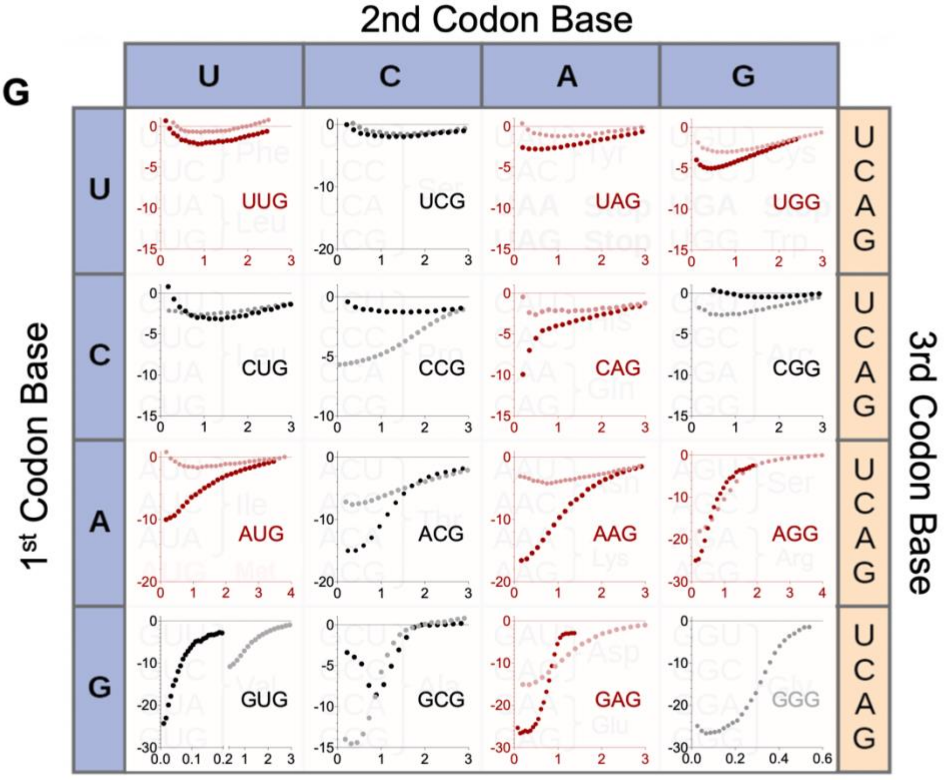


X axis: molar ratio; Y axis: enthalpy change (kcal/mol)

**D) Codons ending in U**

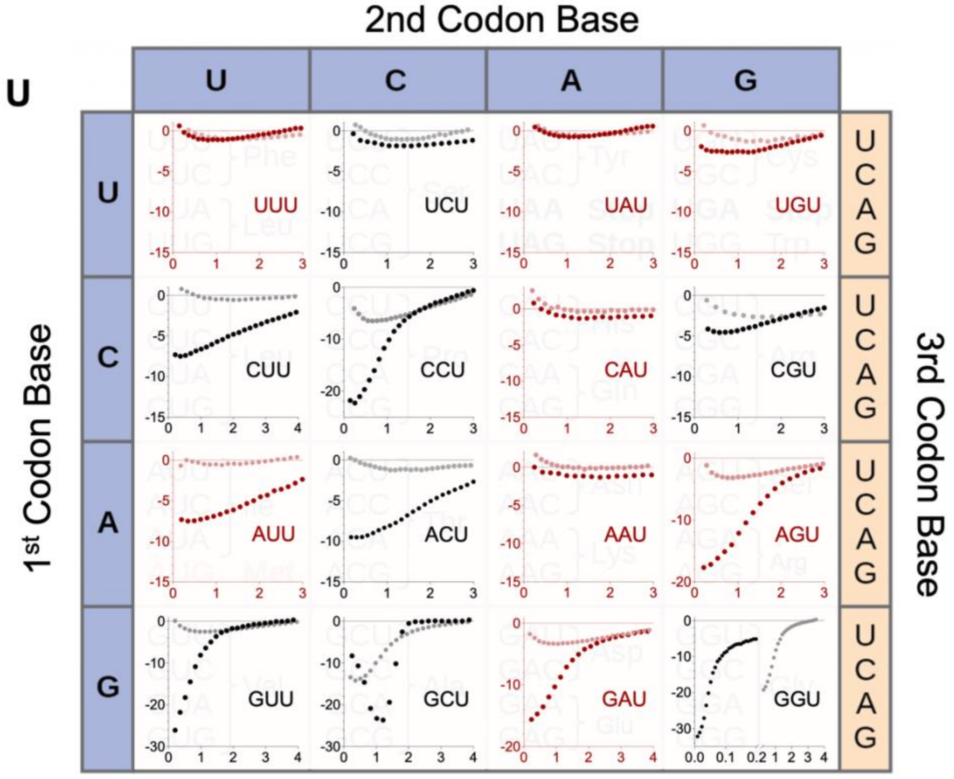


X axis: molar ratio; Y axis: enthalpy change (kcal/mol)

**Fig. S12** Isothermal titration calorimetry curves of family-box anticodon loop interacting with another family-box loop. Conditions: 800 µM anticodon loops, 100 mM NaCl, 100 mM $MgCl_2$, 50 mM HEPES, pH 7.2, 10 ºC.

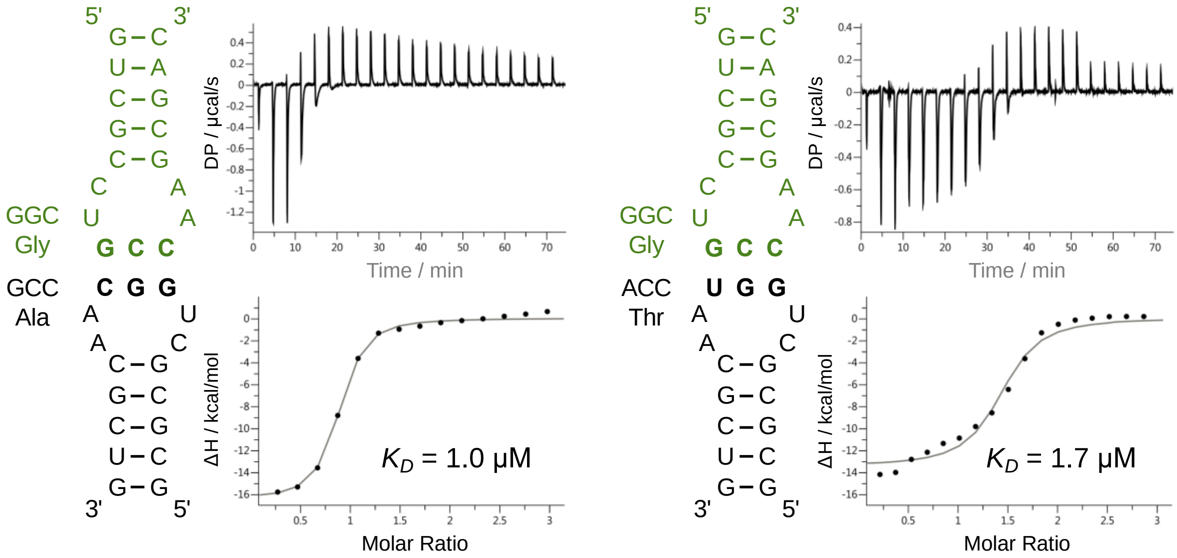

**Fig. S13** BLI curves depicting the binding of ssRNAs containing either two family-box codons or one family-box (black font) and one split-box codon (blue font). Initially, the ssRNAs were saturated with one codon at a fixed concentration (200 µM). Subsequently, a second loop was added with a gradient concentration (ranging from 0 to 400 µM), whilst maintaining the first loop at the same fixed concentration. Then, the biotinylated RNA complex was moved back to the first anticodon loop solution to wash out the second loop. The binding measurement was repeated four more times for a total of five cycles. Conditions: 100 mM NaCl, 100 mM $MgCl_2$, 50 mM HEPES, pH 7.2, 20 ºC.

5'-A CCC UCC G   (Pro-Ser)

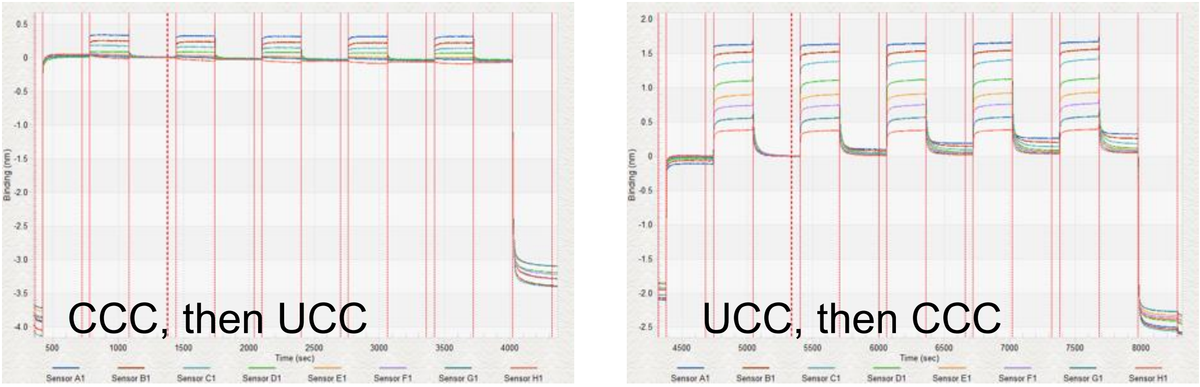


5'-A ACG GUC C  (Thr-Val)

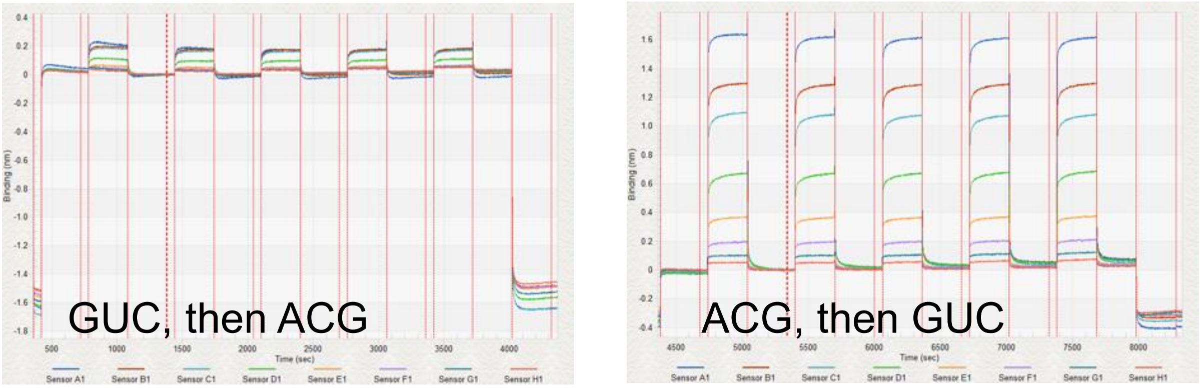


5'-A CGA GGU C  (Arg-Gly)

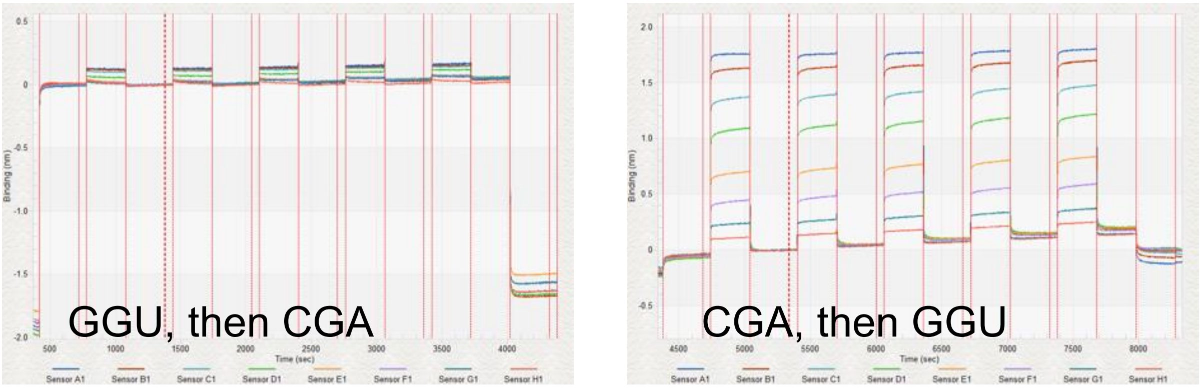

5'-A GAC GCC C (Asp-Ala)

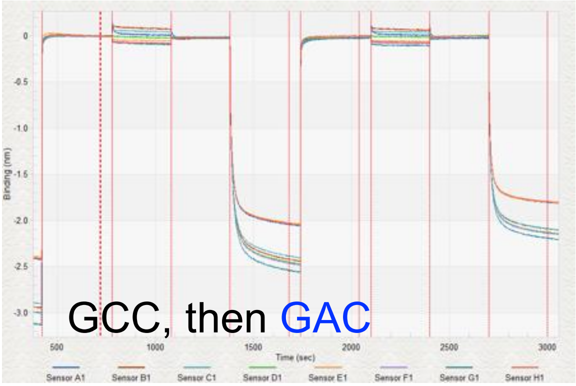


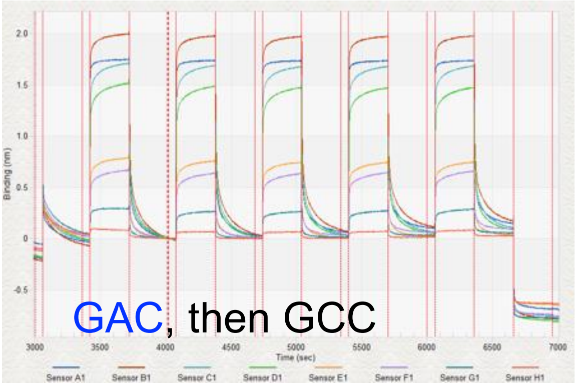


5‘-A GUC UAC C (Val-Tyr)

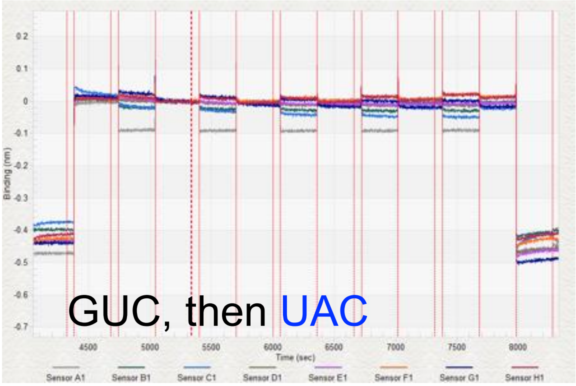


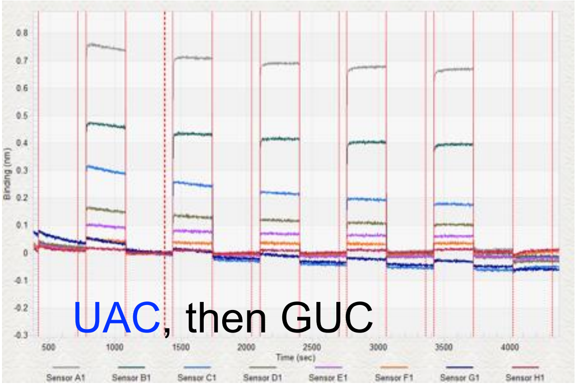

**Fig. S14** Biolayer interferometery binding curves using ssRNA with two different split-box codons. (A) single anticodon loop 5'-CUGAAAA binding to UUC; (B) single anticodon loop 5'-CUAUGAA binding to CAU; (C) equally mixed two anticodon loops aim at binding to UUC and CAU. Neither hairpins binding to UUC or CAU could be included at saturating conditions due to their low binding affinity. Simultaneous binding afforded an apparent $K_D$ of 190 µM. As this is very similar to that we recorded for CAU on its own (180 µM) we conclude only the CAU is binding. Conditions: 0–800 µM each anticodon loop, 100 mM NaCl, 100 mM $MgCl_2$, 50 mM HEPES, pH 7.2, 20 ºC.

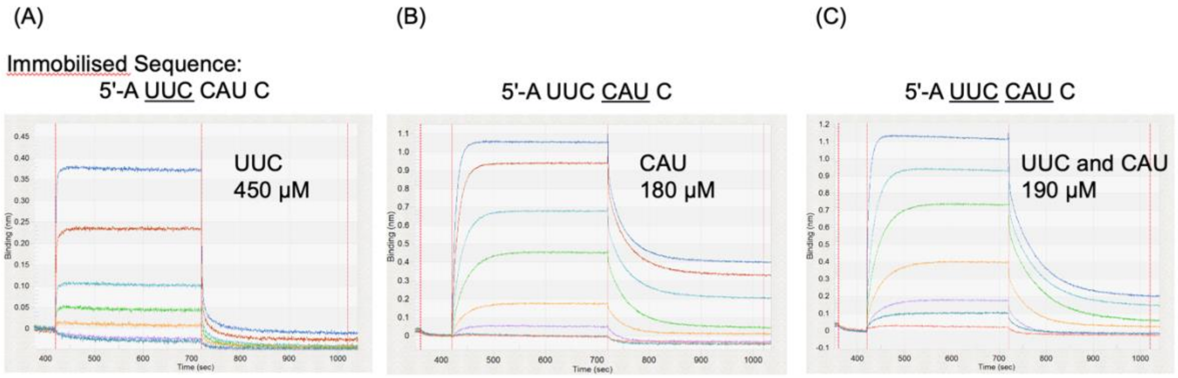

**Fig. S15** Anticodon loops that are able to form self-dimers. General position of loop sequences are underlined. Bold letters indicate anticodon triplets. Anticodon loops for codons ANU, UNA, GNC, and CNG are likely to dimerize, reducing the authentic concentration for the loop in both BLI and ITC and leading to a false stronger equilibrium dissociation constants. The issue was alleviated by denaturing and quickly reannealing the anticodon loops.

```
5' GUCGC  CUANUAA  GCGAC
   |||||   || ||   |||||
3' CAGCG  AAUNAUC  CGCUG

5' GUCGC  CUUNAAA  GCGAC
   |||||   || ||   |||||
3' CAGCG  AAANUUC  CGCUG

5' GUCGC  CUGNCAA  GCGAC
   |||||   || ||   |||||
3' CAGCG  AACNGUC  CGCUG

5' GUCGC  CUCNGAA  GCGAC
   |||||   || ||   |||||
3' CAGCG  AAGNCUC  CGCUG
```

**Fig. S16** Schematic representation of tRNA anticodon loop and homo-triplet binding where four bases pairing is theoretically possible. The numbering on the anticodon loop is consistent with the existing tRNA anticodon loop domain, aligning with the standard structure of a typical tRNA, which comprises 76 nucleotides.

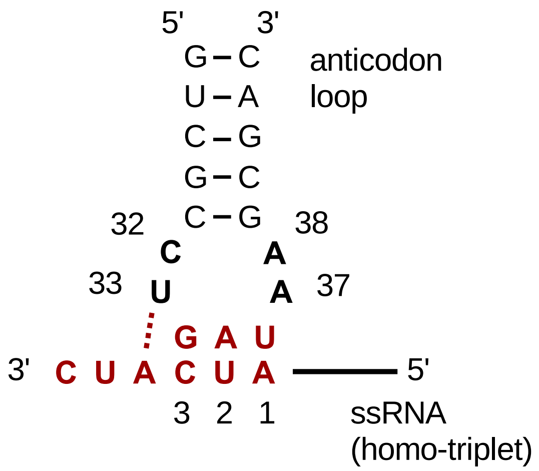

**Fig. S17** Binding affinity of single anticodon-loop (5'- CUNGGAA) and trimer RNA (5'-CCN) with one mismatched base pair.

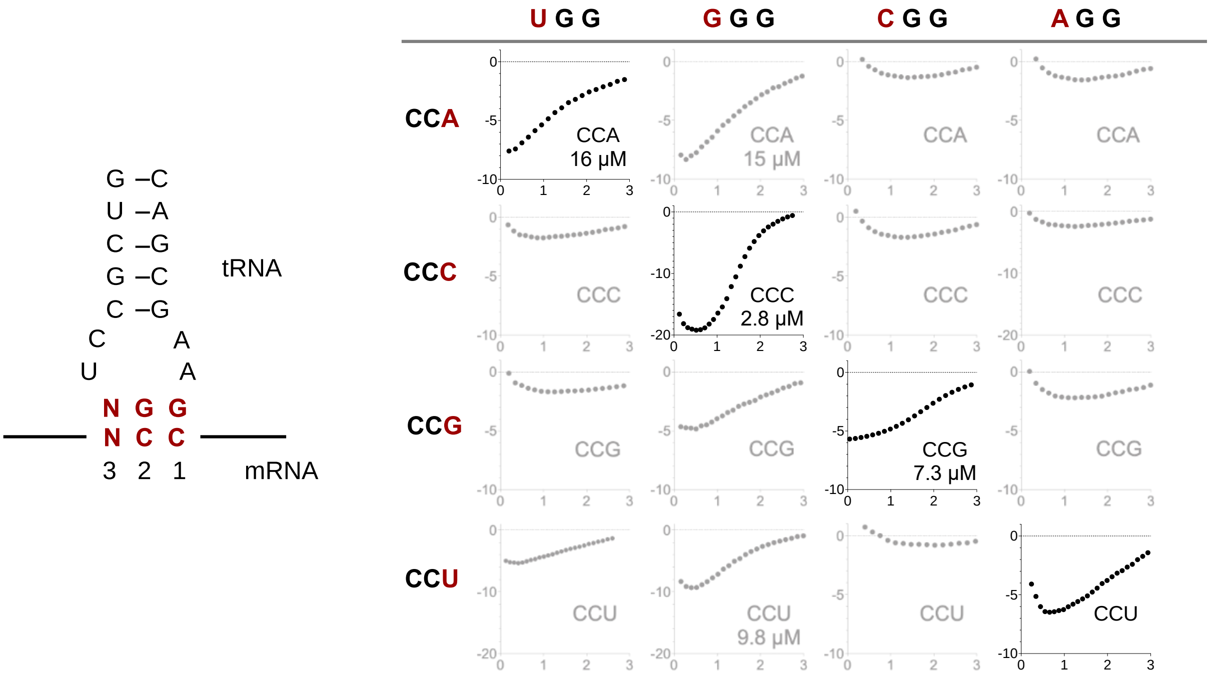

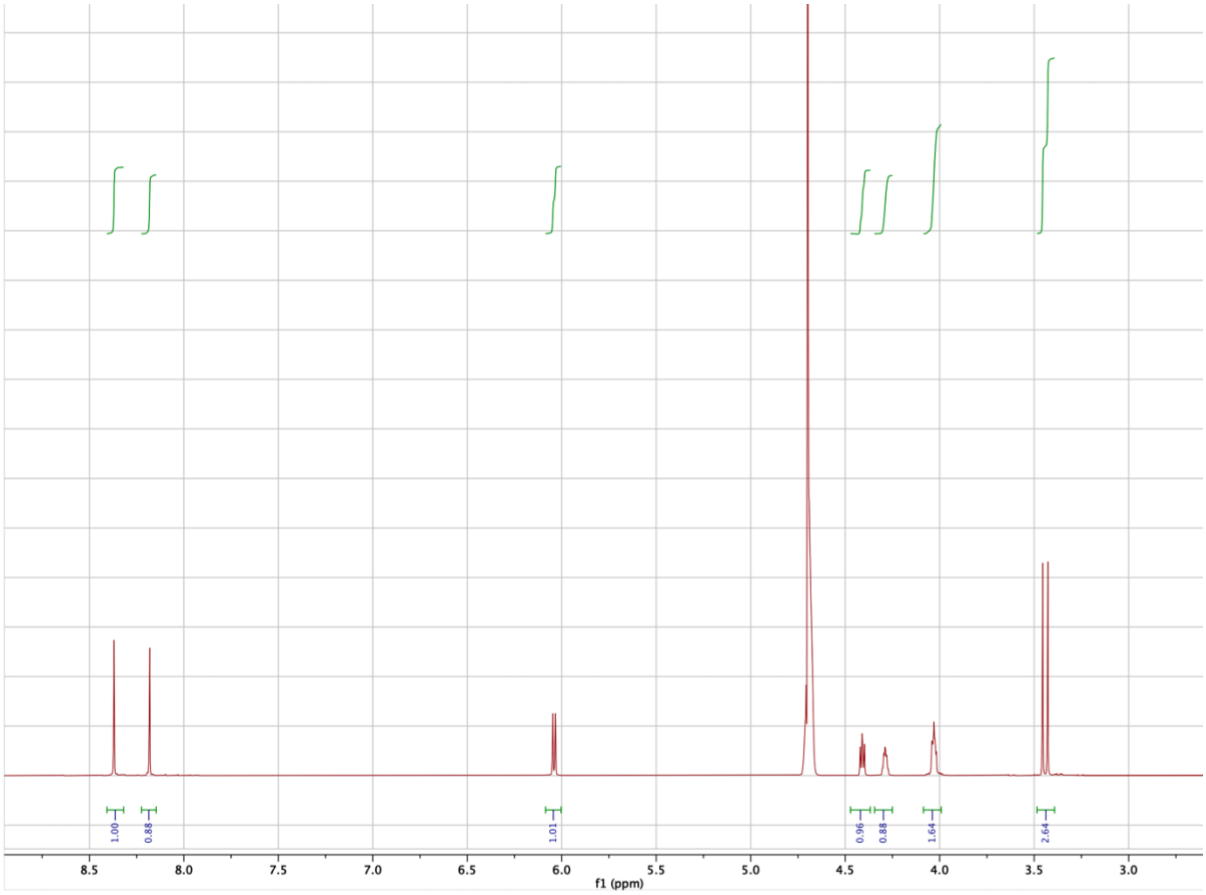


**Fig. S18** [1]H-NMR spectrum of adenosine-5'-*O*-methylphosphate

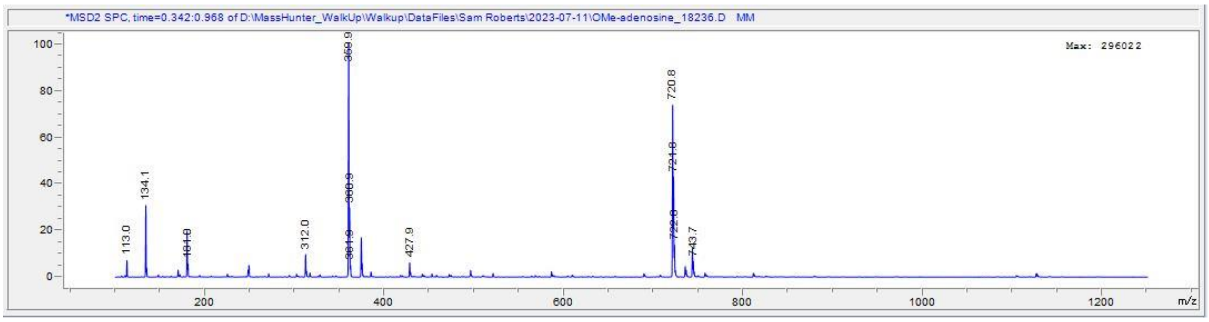


**Fig. S19** Negative ion low resolution mass spectrum of adenosine-5'-*O*-methylphosphate

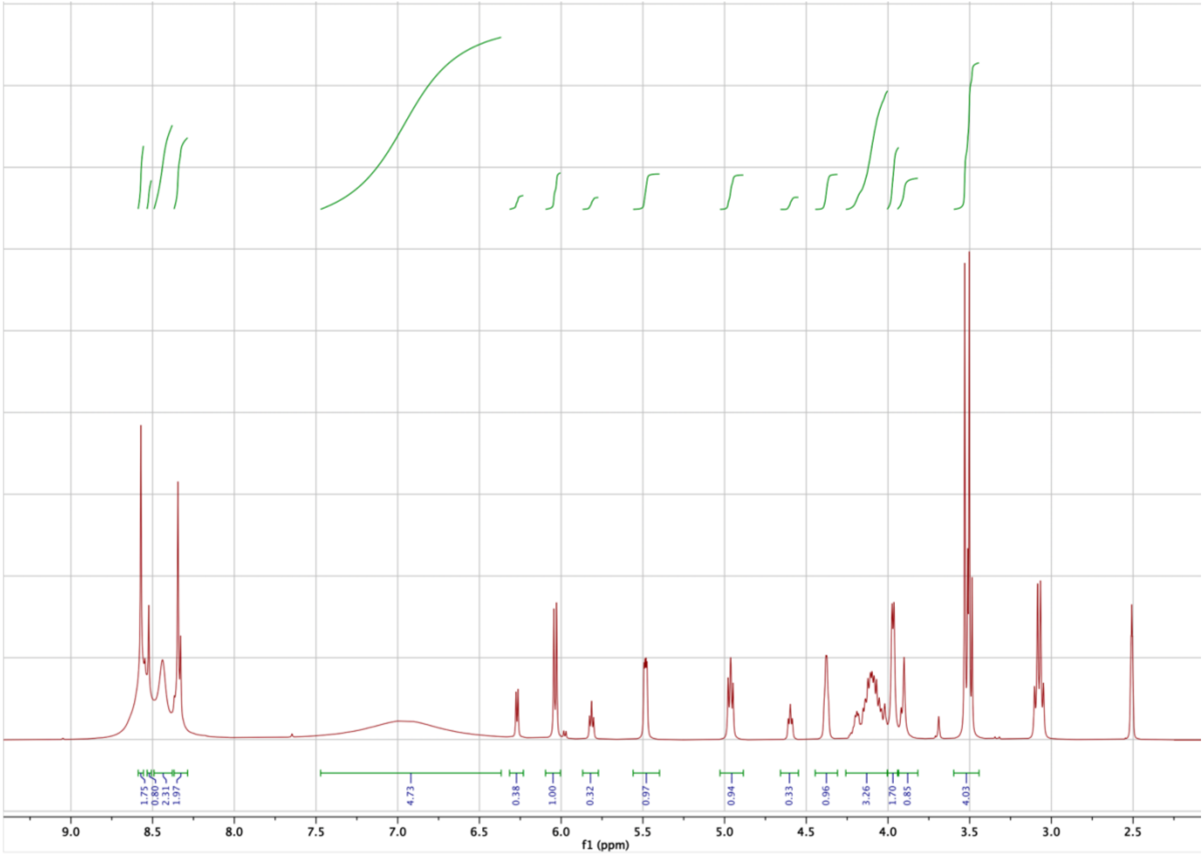

**Fig. S20** $^{1}$H-NMR spectrum of 2'/3'-glycyl-adenosine-5'-*O*-methylphosphate

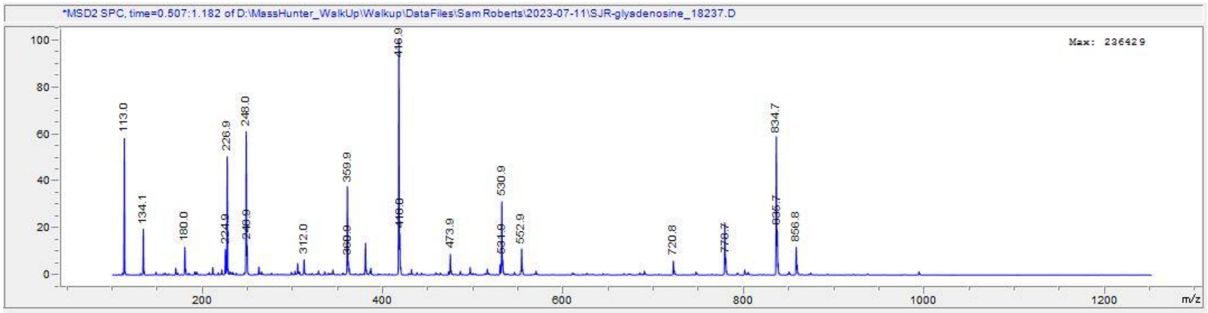

**Fig. S21 –** Negative ion low resolution mass spectrum of 2'/3'-glycyl-adenosine-5'-*O*-methylphosphate

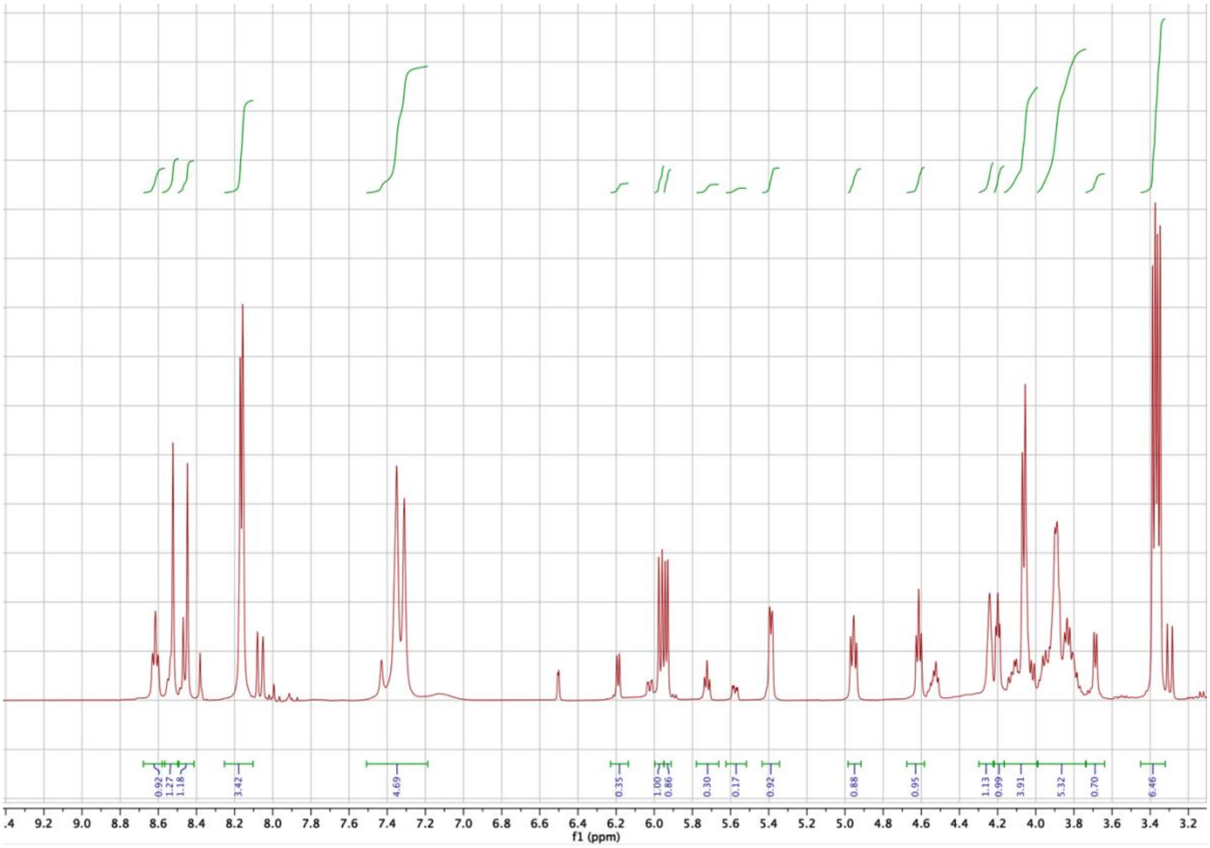

**Fig. S22** $^{1}$H-NMR spectrum of 2'/3'-*N*-formyl-glycyl-adenosine-5'-*O*-methylphosphate

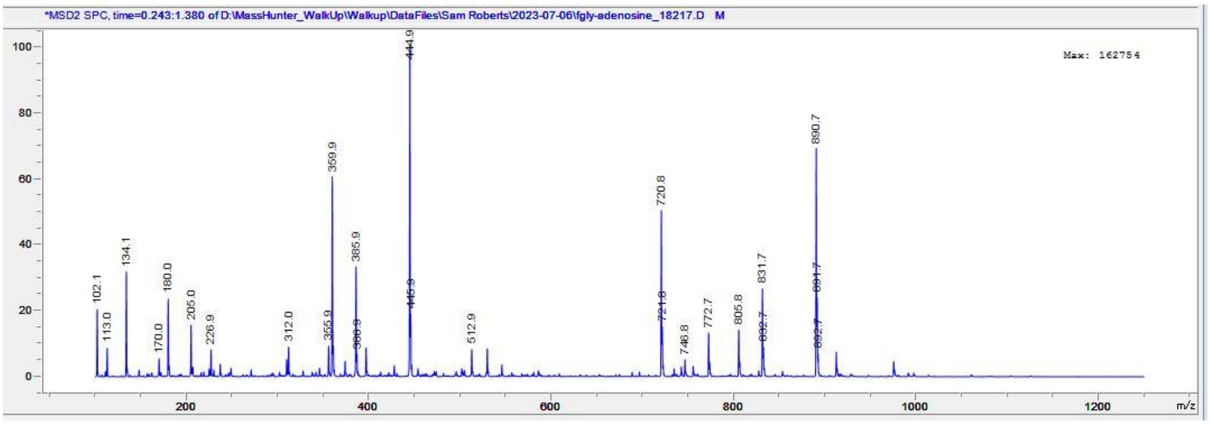


**Fig. S23** Negative ion low resolution mass spectrum of 2'/3'-*N*-formyl-*g*lycyl-adenosine-5'-*O*-methylphosphate

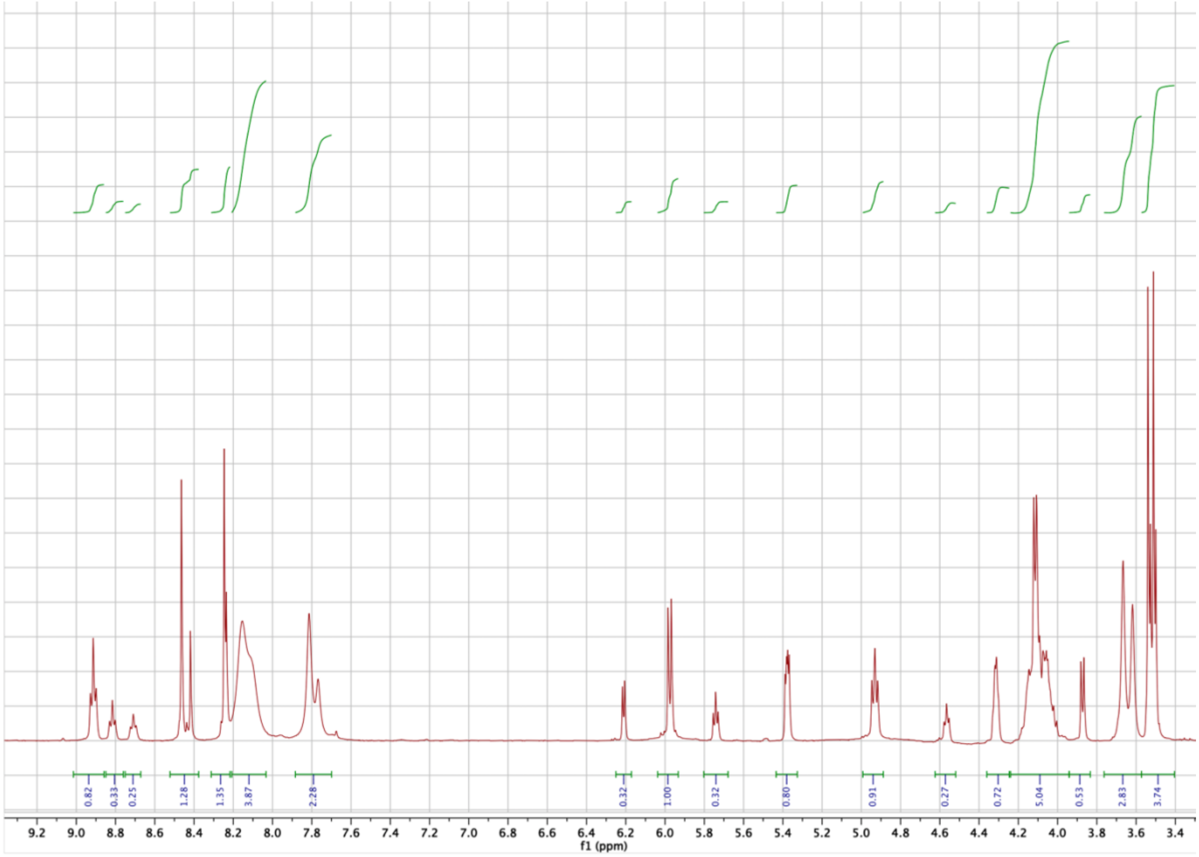


**Fig. S24** [1]H-NMR spectrum of 2'/3'-diglycyl-adenosine-5'-*O*-methylphosphate

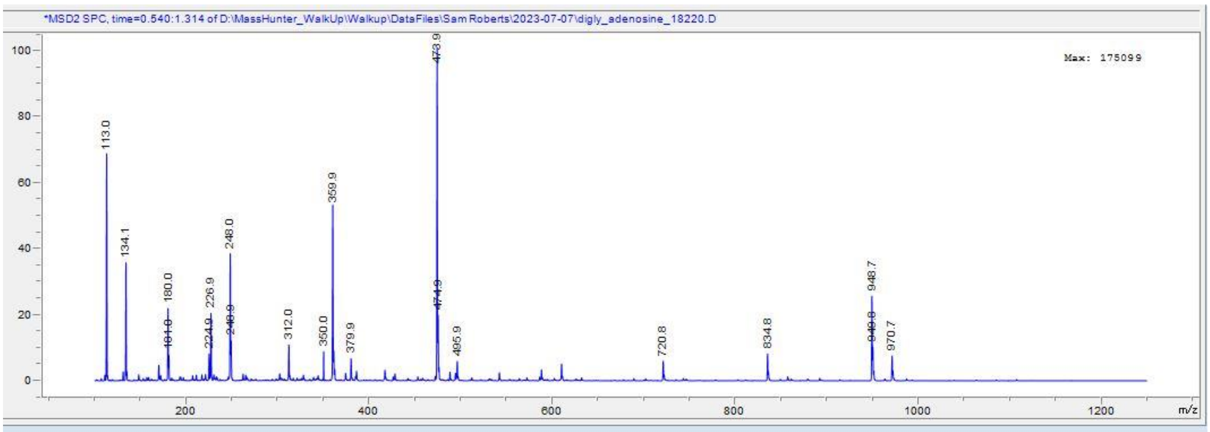


**Fig. S25** Negative ion low resolution mass spectrum 2'/3'-diglycyl-adenosine-5'-*O*-methylphosphate

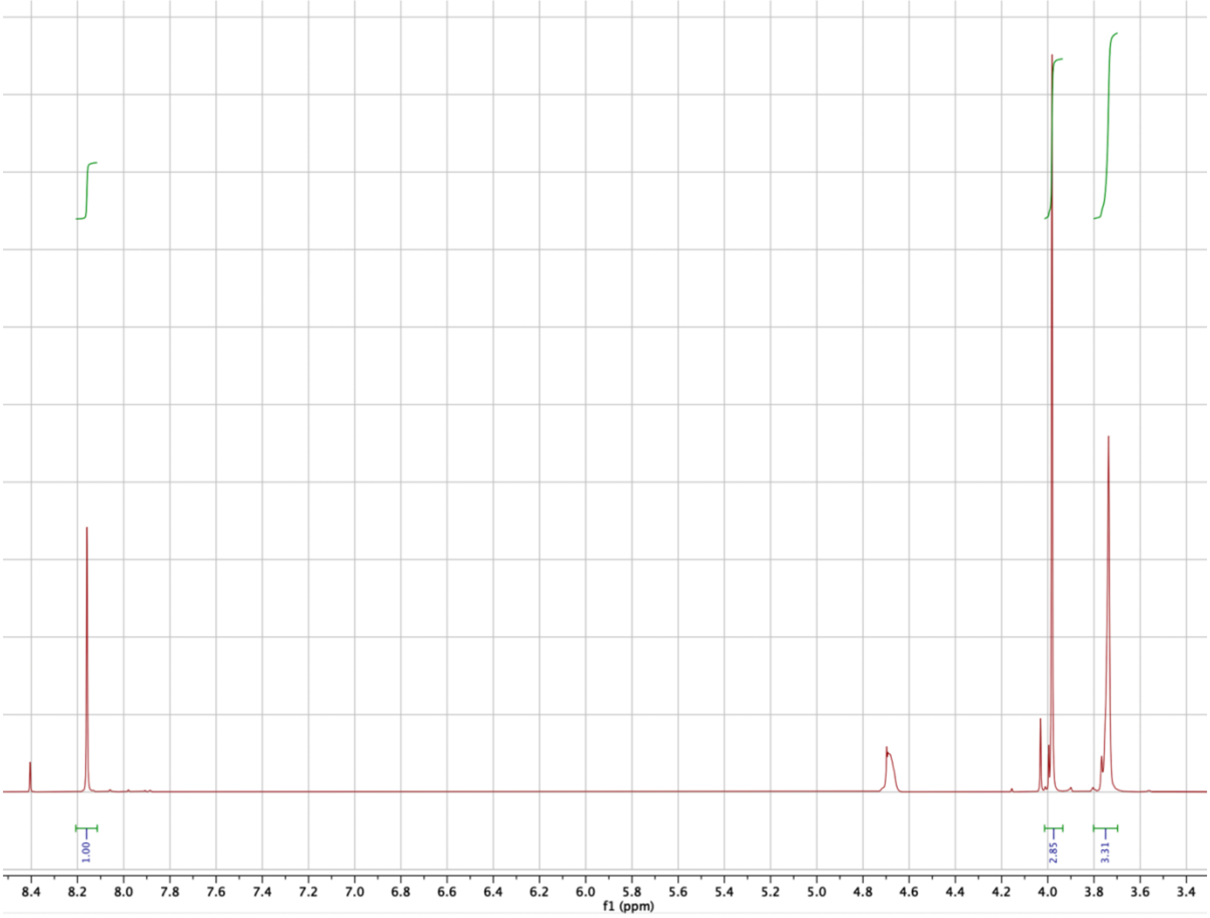


**Fig. S26** [1]H-NMR spectrum of *N*-formyl-diglycine

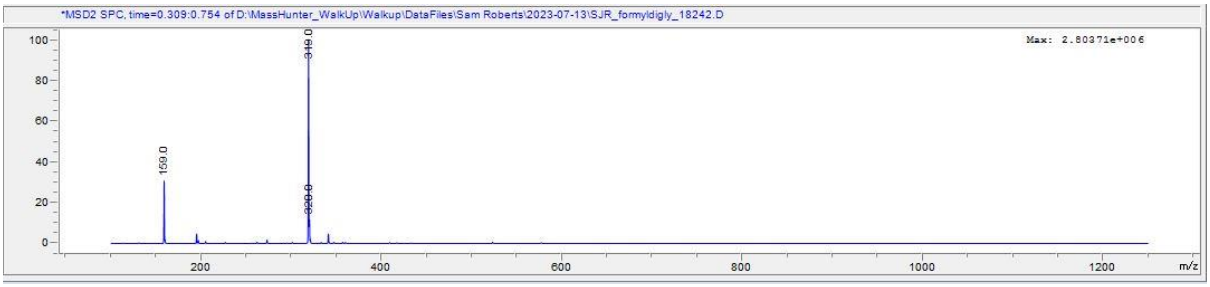


**Fig. S27** Negative ion low-resolution mass spectrum *N*-formyl-diglycine

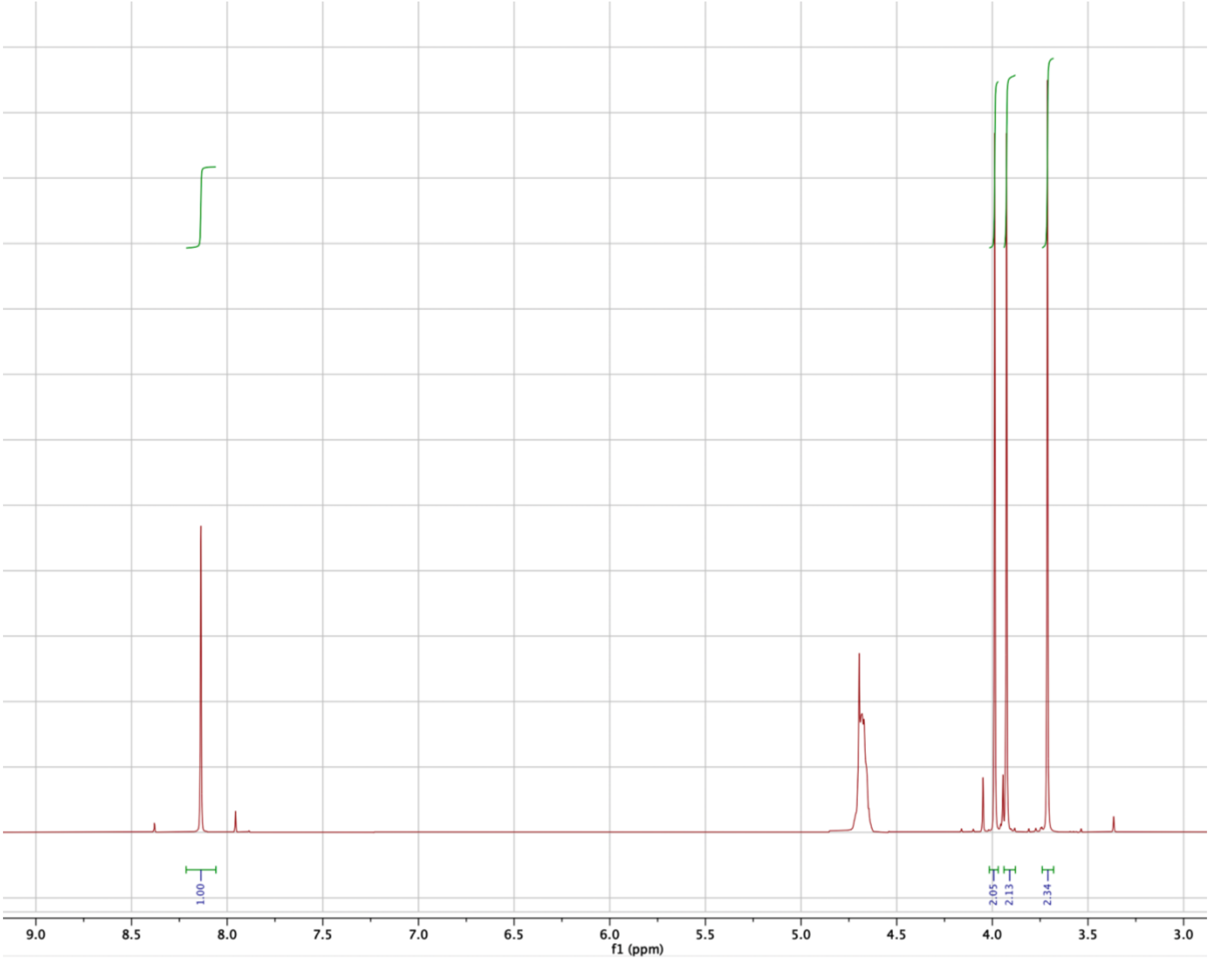


**Fig. S28** $^{1}$H-NMR spectrum of *N*-formyl-triglycine

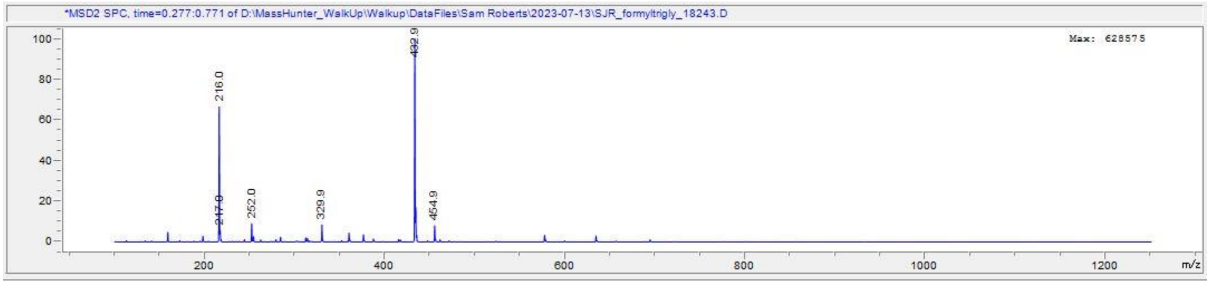


**Fig. S29 –** Negative ion low resolution mass spectrum *N*-formyl-triglycine

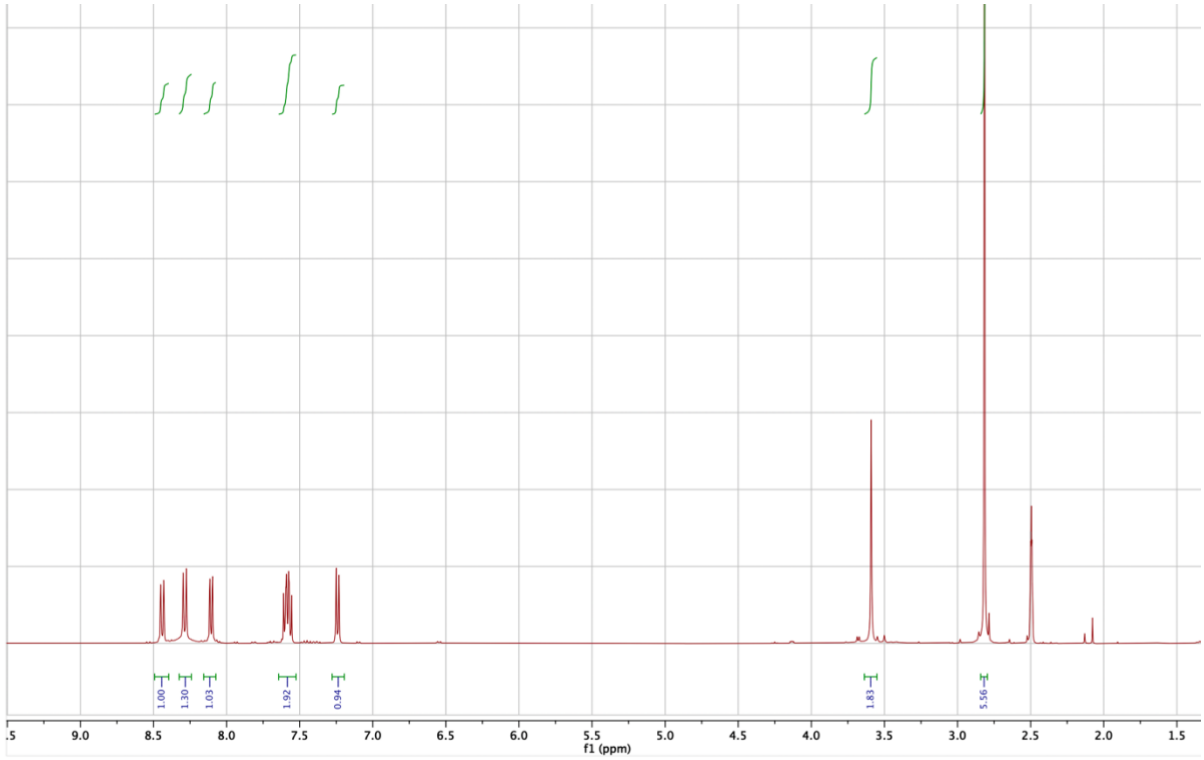


**Fig. S30** [1]H-NMR spectrum of *N*-dansyl-glycine

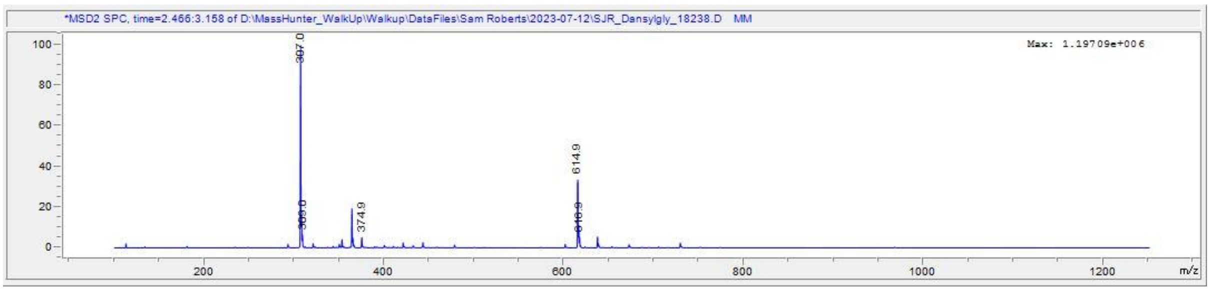


**Fig. S31** Negative ion low resolution mass spectrum *N*-dansyl-glycine

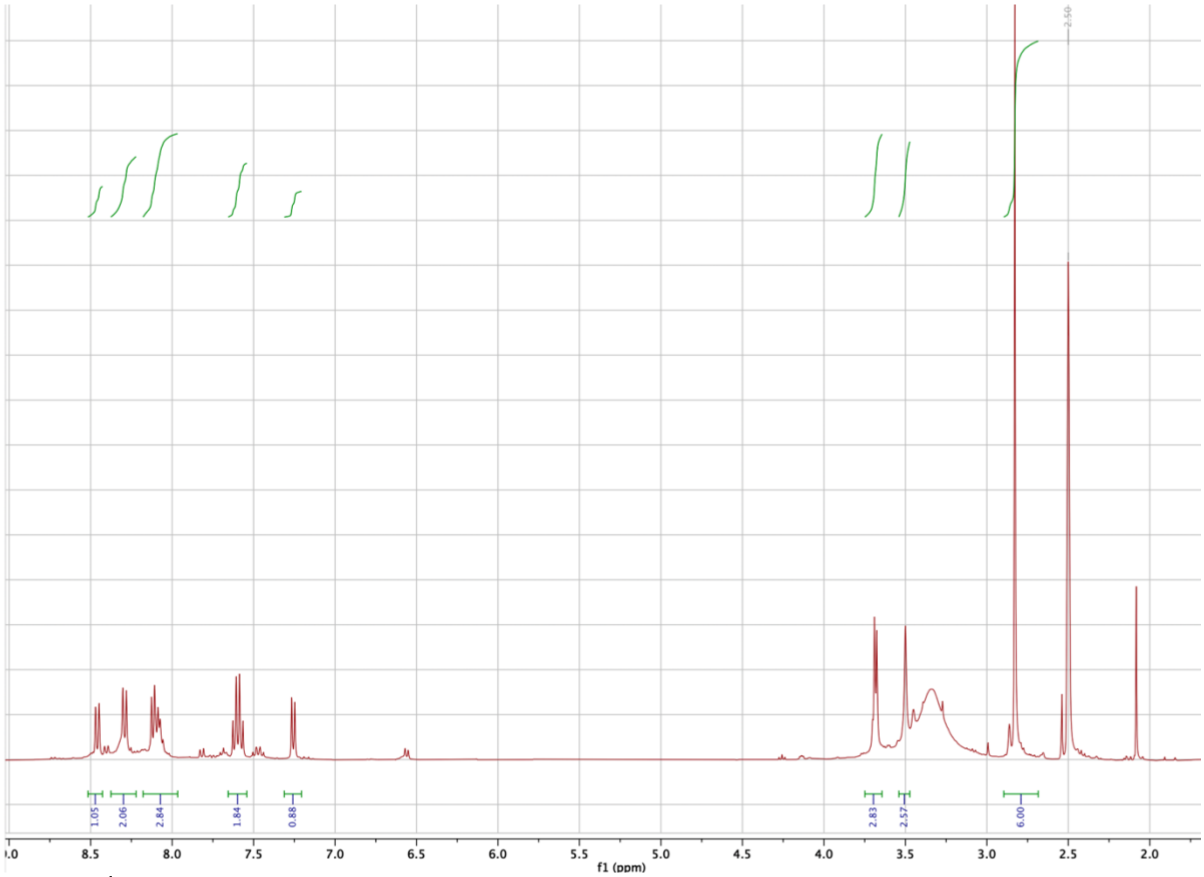


**Fig. S32** $^{1}$H-NMR spectrum of *N*-dansyl-diglycine

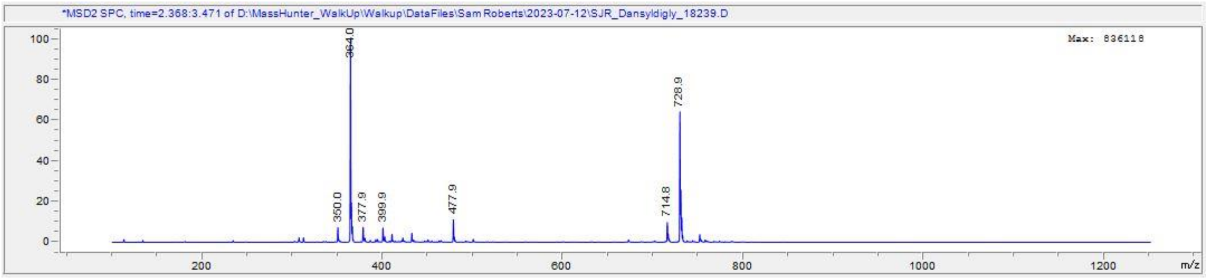


**Fig. S33** Negative ion low-resolution mass spectrum *N*-dansyl-diglycine

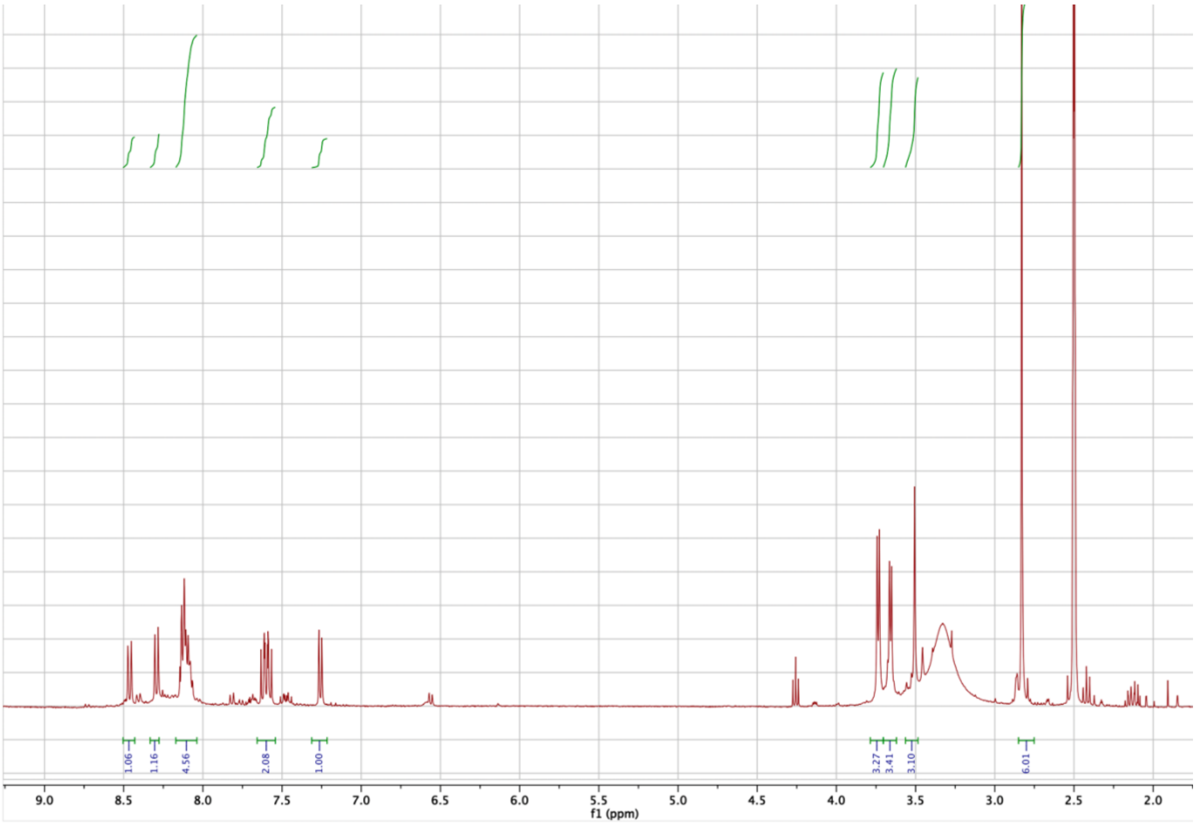


**Fig. S34** [1]H-NMR spectrum of *N*-dansyl-triglycine

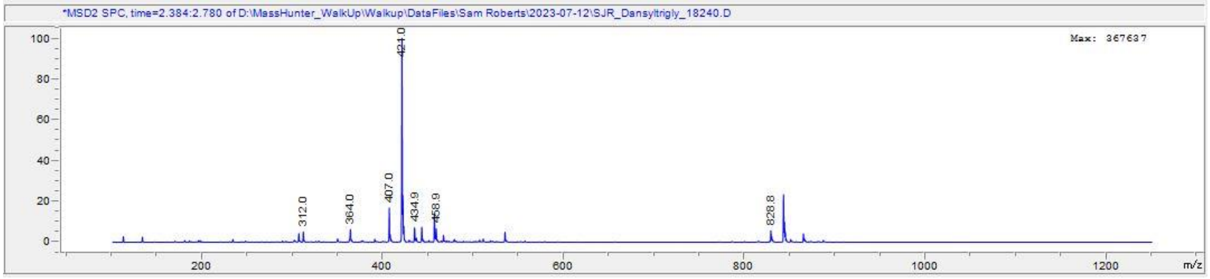


**Fig. S35** Negative ion low-resolution mass spectrum *N*-dansyl-triglycine

**Table S1** Comparison of the yields (%) of dansylation of a mixture of glycine, diglycine and triglycine with the observed ratio of the mixture by $^{1}H$ NMR.

| | | | |
|---|---|---|---|
| Yields (%) By Dansylation | 23.97 | 50.91 | 101.97 |
| Expected Yields (%) By NMR | 20.50 | 43.56 | 100.00 |

**Table S2** Yields (%) relative to the total amount of glycine species from the reactions of adenylates presented in this paper. Quantified by $^1$H NMR direct integration (blue), $^1$H NMR integration after the addition of known amounts of standard (std., black) or by HPLC analysis post-dansyl functionalisation (red). * Lack of a nucleophilic centre (free amino group) prevents dansyl functionalisation. ** Peak could not be observed by $^1$H NMR due to apparent resonance broadening.

| Reagents | Method | Tri-Gly | Di-Gly | Gly | DKP* | Formyl -trigly* | Formyl-digly* | Formyl-gly* |
|---|---|---|---|---|---|---|---|---|
| Gly-adenosine (415 mM) | Dansyl | 0.12 | 3.06 | 71.0 | - | - | - | - |
| | NMR | 0.36 | 5.64 | 75.7 | 11.8 | - | - | - |
| | NMR std. | 0.50 | 6.09 | - | 12.8 | - | - | - |
| Gly-adenosine + Glycine (207 mM both) | Dansyl | 0.05 | 2 | 63.2 | - | - | - | - |
| | NMR | 0.09 | 3.39 | 83.8 | 2.97 | - | - | - |
| | NMR std. | 0.09 | 3.32 | - | 3 | - | - | - |
| Gly-adenosine (35 mM) | NMR | 0.00 | 0.42 | 94.7 | 1.13 | - | - | - |
| | NMR std. | 0.00 | 0.42 | - | 1.26 | - | - | - |
| Gly-adenosine + Formyl-gly-adenosine (207 mM both) | Dansyl | 0.04 | 1.36 | 44.8 | - | - | - | - |
| | NMR | 0.12 | 1.74 | -** | 5.18 | 0.15 | 5.44 | 31.01 |
| | NMR std. | 0.64 | 1.80 | - | 5.61 | 0.13 | 5.33 | - |
| Digly-adenosine (10mM) | NMR | - | 22.7 | - | 77.3 | - | - | - |

**Table S3** Synthesized anticodon loops RNAs for the BLI and ITC experiments. Found, observed (raw) mass number from LCMS in negative ion mode; Expt., the experimental mass number calculated from the observed values, i.e. Expt. = (Found + 1) × charges, 4 or 5 charges; Calc., mass number in theory. Underlined residues highlight the loop sequence.

| No. | Sequence (5' -3') | Calc. | Expt. | Found |
|---|---|---|---|---|
| 1 | GUCGC CU UUU AA GCGAC | 5360.7 | 5360.0 | 1071.3 |
| 2 | GUCGC CU GUU AA GCGAC | 5399.8 | 5400.4 | 1349.1 |
| 3 | GUCGC CU CUU AA GCGAC | 5359.8 | 5360.0 | 1071.1 |
| 4 | GUCGC CU AUU AA GCGAC | 5383.8 | 5384.5 | 1075.9 |
| 5 | GUCGC CU UGU AA GCGAC | 5399.8 | 5401.0 | 1079.2 |
| 6 | GUCGC CU GGU AA GCGAC | 5438.8 | 5439.2 | 1358.8 |
| 7 | GUCGC CU CGU AA GCGAC | 5398.8 | 5400.0 | 1079.0 |
| 8 | GUCGC CU AGU AA GCGAC | 5422.8 | 5424.0 | 1083.8 |
| 9 | GUCGC CU UCU AA GCGAC | 5359.8 | 5360.0 | 1071.0 |
| 10 | GUCGC CU GCU AA GCGAC | 5398.8 | 5398.5 | 1078.7 |
| 11 | GUCGC CU CCU AA GCGAC | 5358.8 | 5360.0 | 1071.0 |
| 12 | GUCGC CU ACU AA GCGAC | 5382.8 | 5383.0 | 1075.6 |
| 13 | GUCGC CU UAU AA GCGAC | 5383.8 | 5384.5 | 1075.9 |
| 14 | GUCGC CU GAU AA GCGAC | 5422.8 | 5423.6 | 1354.9 |
| 15 | GUCGC CU CAU AA GCGAC | 5382.8 | 5383.5 | 1075.7 |
| 16 | GUCGC CU AAU AA GCGAC | 5406.8 | 5407.5 | 1080.5 |
| 17 | GUCGC CU UUG AA GCGAC | 5399.8 | 5400.0 | 1079.0 |
| 18 | GUCGC CU GUG AA GCGAC | 5438.8 | 5439.6 | 1358.9 |
| 19 | GUCGC CU CUG AA GCGAC | 5398.8 | 5399.5 | 1078.9 |
| 20 | GUCGC CU AUG AA GCGAC | 5422.8 | 5423.5 | 1083.7 |
| 21 | GUCGC CU UGG AA GCGAC | 5438.8 | 5439.5 | 1086.9 |
| 22 | GUCGC CU GGG AA GCGAC | 5477.8 | 5478.5 | 1094.7 |
| 23 | GUCGC CU CGG AA GCGAC | 5437.8 | 5438.8 | 1358.7 |
| 24 | GUCGG CU AGG AA CCGAC | 5461.8 | 5463.0 | 1091.6 |
| 25 | GUCGC CU UCG AA GCGAC | 5398.8 | 5399.5 | 1078.9 |
| 26 | GUCGC CU GCG AA GCGAC | 5437.8 | 5439.5 | 1086.9 |
| 27 | GUCGC CU CCG AA GCGAC | 5397.8 | 5399.0 | 1078.8 |
| 28 | GUCGC CU ACG AA GCGAC | 5421.8 | 5421.5 | 1083.3 |
| 29 | GUCGC CU UAG AA GCGAC | 5422.8 | 5423.0 | 1083.6 |
| 30 | GUCGC CU GAG AA GCGAC | 5461.8 | 5462.5 | 1091.5 |
| 31 | GUCGC CU CAG AA GCGAC | 5421.8 | 5421.5 | 1083.3 |
| 32 | GUCGC CU AAG AA GCGAC | 5445.8 | 5446.5 | 1088.3 |
| 33 | GUCGC CU UUC AA GCGAC | 5359.8 | 5360.5 | 1071.1 |
| 34 | GUCGC CU GUC AA GCGAC | 5398.8 | 5399.6 | 1348.9 |
| 35 | GUCGC CU CUC AA GCGAC | 5358.8 | 5359.6 | 1338.9 |
| 36 | GUCGC CU AUC AA GCGAC | 5382.8 | 5383.0 | 1075.6 |
| 37 | GUCGG CU UGC AA CCGAC | 5398.8 | 5399.5 | 1078.9 |

| | | | | |
|---|---|---|---|---|
| 38 | GUCGC CU GGC AA GCGAC | 5437.8 | 5438.4 | 1358.6 |
| 39 | GUCGC CU CGC AA GCGAC | 5397.8 | 5398.0 | 1348.5 |
| 40 | GUCGC CU AGC AA GCGAC | 5421.8 | 5422.5 | 1083.5 |
| 41 | GUCGC CU UCC AA GCGAC | 5358.8 | 5358.5 | 1070.7 |
| 42 | GUCGC CU GCC AA GCGAC | 5397.8 | 5399.0 | 1078.8 |
| 43 | GUCGC CU CCC AA GCGAC | 5357.8 | 5359.0 | 1070.8 |
| 44 | GUCGC CU ACC AA GCGAC | 5381.8 | 5382.0 | 1075.4 |
| 45 | GUCGC CU UAC AA GCGAC | 5382.8 | 5383.2 | 1344.8 |
| 46 | GUCGC CU GAC AA GCGAC | 5421.8 | 5422.0 | 1083.4 |
| 47 | GUCGC CU CAC AA GCGAC | 5381.8 | 5382.4 | 1344.6 |
| 48 | GUCGC CU AAC AA GCGAC | 5405.8 | 5407.0 | 1080.4 |
| 49 | GUCGC CU UUA AA GCGAC | 5383.8 | 5384.5 | 1075.9 |
| 50 | GUCGC CU GUA AA GCGAC | 5422.8 | 5422.5 | 1083.5 |
| 51 | GUCGC CU CUA AA GCGAC | 5382.8 | 5383.5 | 1075.7 |
| 52 | GUCGC CU AUA AA GCGAC | 5406.8 | 5407.5 | 1080.5 |
| 53 | GUCGC CU UGA AA GCGAC | 5422.8 | 5424.0 | 1083.8 |
| 54 | GUCGC CU GGA AA GCGAC | 5461.8 | 5462.0 | 1364.5 |
| 55 | GUCGC CU CGA AA GCGAC | 5421.8 | 5423.0 | 1083.6 |
| 56 | GUCGC CU AGA AA GCGAC | 5445.8 | 5446.5 | 1088.3 |
| 57 | GUCGC CU UCA AA GCGAC | 5382.8 | 5383.5 | 1075.7 |
| 58 | GUCGC CU GCA AA GCGAC | 5421.8 | 5422.8 | 1354.7 |
| 59 | GUCGC CU CCA AA GCGAC | 5381.8 | 5381.0 | 1075.2 |
| 60 | GUCGC CU ACA AA GCGAC | 5405.8 | 5407.0 | 1080.4 |
| 61 | GUCGC CU UAA AA GCGAC | 5406.8 | 5407.6 | 1350.9 |
| 62 | GUCGC CU GAA AA GCGAC | 5445.8 | 5446.8 | 1360.7 |
| 63 | GUCGC CU CAA AA GCGAC | 5405.8 | 5406.5 | 1080.3 |
| 64 | GUCGC CU AAA AA GCGAC | 5429.8 | 5429.5 | 1084.9 |

**Table S4** Purchased biotin-labelled oligoribonucleotides for BLI experiments. In most cases, the first and final nucleosides of the octamer were fixed to adenosine and cytosine and were not used as a part of the codon.

| No. | Sequence (5' -3') |
|---|---|
| 400 | Biotin-C UUC CAU U |
| 401 | Biotin-A GUC ACC C |
| 402 | Biotin-A UUC CAU C |
| 403 | Biotin-A GCC CUC C |
| 404 | Biotin-A GUU UCU C |
| 405 | Biotin-A GUA UCG C |
| 406 | Biotin-A GUG CUU C |
| 407 | Biotin-A GCA ACU C |
| 408 | Biotin-A GCG ACA C |
| 409 | Biotin-A GAU GGU C |
| 410 | Biotin-A GAG UUA C |
| 411 | Biotin-A GGC CGU C |
| 412 | Biotin-A GGA CGC C |
| 413 | Biotin-A GGG CUA C |
| 414 | Biotin-A CUG UUG C |
| 415 | Biotin-A AUA AGC C |
| 416 | Biotin-A AAU UAC C |
| 417 | Biotin-A AGA CAG C |
| 418 | Biotin-A UAG AAA C |
| 419 | Biotin-A UGA AGG C |
| 420 | Biotin-A CGG UCA C |
| 421 | Biotin-A GCA CGG C |
| 422 | Biotin-A CGG GCU C |
| 423 | Biotin-A GGA UCG C |
| 424 | Biotin-A UGA UAG C |
| 425 | Biotin-A CCG UCC C |
| 426 | Biotin-A GAC GAA U |
| 427 | Biotin-C GAC GAA C |
| 428 | Biotin-A GGU CUU C |
| 429 | Biotin-A CCA GCG U |
| 430 | Biotin-A CGA GGU C |
| 431 | Biotin-A ACG GUC C |
| 432 | Biotin-A CCC UCC G |
| 433 | Biotin-A GAC GCC C |
| 434 | Biotin-A GUC UAC C |

**Table S5** Synthesized trinucleotides (mono-triplets) and purchased hexanucleotides (homo-triplets) for the ITC experiments. Found, observed (raw) mass number from LCMS in negative ion mode; Expt., the experimental mass number calculated from the observed values, i.e. Expt. = Found + 1; Calc., mass number in theory.

| No. | Triplet | Calc. | Expt. | Found | Homo-triplet (5' -3') |
|---|---|---|---|---|---|
| 1 | AAA | 925.2 | 924.9 | 923.9 | AAA AAA C |
| 2 | AAC | 901.2 | 901.0 | 900.0 | AAC AAC C |
| 3 | AAG | 941.2 | 941.0 | 940.0 | AAG AAG U |
| 4 | AAU | 902.2 | 902.0 | 901.0 | AAU AAU U |
| 5 | ACA | 901.2 | 901.0 | 900.0 | ACA ACA U |
| 6 | ACC | 877.2 | 876.9 | 875.9 | C ACC ACC |
| 7 | ACG | 917.2 | 917.0 | 916.0 | ACG ACG |
| 8 | ACU | 878.2 | 878.0 | 877.0 | ACU ACU U |
| 9 | AGA | 941.2 | 940.9 | 939.9 | AGA AGA U |
| 10 | AGC | 917.2 | 917.0 | 916.0 | C AGC AGC |
| 11 | AGG | 957.2 | 956.9 | 955.9 | AGG AGG U |
| 12 | AGU | 918.2 | 917.9 | 916.9 | AGU AGU U |
| 13 | AUA | 902.2 | 902.0 | 901.0 | AUA AUA U |
| 14 | AUC | 878.2 | 878.0 | 877.0 | C AUC AUC |
| 15 | AUG | 918.2 | 917.9 | 916.9 | G AUG AUG |
| 16 | AUU | 879.2 | 878.9 | 877.9 | AUU AUU U |
| 17 | CAA | 901.2 | 900.9 | 899.9 | CAA CAA UU |
| 18 | CAC | 877.2 | 877.0 | 876.0 | CAC CAC C |
| 19 | CAG | 917.2 | 916.9 | 915.9 | CAG CAG C |
| 20 | CAU | 878.2 | 877.9 | 876.9 | CAU CAU C |
| 21 | CCA | 877.2 | 877.0 | 876.0 | CCA CCA UU |
| 22 | CCC | 853.2 | 853.0 | 852.0 | CCC CCC |
| 23 | CCG | 893.2 | 892.1 | 891.1 | G CCG CCG |
| 24 | CCU | 854.2 | 853.9 | 852.9 | CCU CCU UU |
| 25 | CGA | 917.2 | 917.0 | 916.0 | CGA CGA C |
| 26 | CGC | 893.2 | 892.9 | 891.9 | CGC CGC U |
| 27 | CGG | 933.2 | 933.0 | 932.0 | CGG CGG |
| 28 | CGU | 894.2 | 893.9 | 892.9 | CGU CGU UU |
| 29 | CUA | 878.2 | 877.9 | 876.9 | CUA CUA C |
| 30 | CUC | 854.2 | 853.9 | 852.9 | CUC CUC CC |
| 31 | CUG | 894.2 | 893.9 | 892.9 | G CUG CUG |
| 32 | CUU | 855.1 | 854.9 | 853.9 | CUU CUU C |
| 33 | GAA | 941.2 | 940.9 | 939.9 | GAA GAA UU |
| 34 | GAC | 917.2 | 916.9 | 915.9 | C GAC GAC |
| 35 | GAG | 957.2 | 956.9 | 955.9 | GAG GAG U |
| 36 | GAU | 918.2 | 917.9 | 916.9 | GAU GAU G |
| 37 | GCA | 917.2 | 916.9 | 915.9 | GCA GCA UU |
| 38 | GCC | 893.2 | 892.9 | 891.9 | GCC GCC G |
| 39 | GCG | 933.2 | 933.2 | 932.2 | G GCG GCG |
| 40 | GCU | 894.2 | 893.9 | 892.9 | GCU GCU G |
| 41 | GGA | 957.2 | 956.9 | 955.9 | GGA GGA U |
| 42 | GGC | 933.2 | 933.0 | 932.0 | GGC GGC G |

| | | | | | |
|---|---|---|---|---|---|
| 43 | GGG | 973.2 | 973.0 | 972.0 | GGG GGG |
| 44 | GGU | 934.2 | 933.9 | 932.9 | GGU GGU G |
| 45 | GUA | 918.2 | 918.0 | 917.0 | GUA GUA G |
| 46 | GUC | 894.2 | 893.9 | 892.9 | GUC GUC G |
| 47 | GUG | 934.2 | 933.9 | 932.9 | G GUG GUG |
| 48 | GUU | 895.2 | 894.9 | 893.9 | GUU GUU G |
| 49 | UAA | 902.2 | 901.9 | 900.9 | UAA UAA U |
| 50 | UAC | 878.2 | 877.9 | 876.9 | UAC UAC CC |
| 51 | UAG | 918.2 | 917.9 | 916.9 | G UAG UAG |
| 52 | UAU | 879.2 | 878.9 | 877.9 | UAU UAU U |
| 53 | UCA | 878.2 | 877.9 | 876.9 | UCA UCA U |
| 54 | UCC | 854.2 | 853.9 | 852.9 | UCC UCC UU |
| 55 | UCG | 894.2 | 893.9 | 892.9 | G UCG UCG |
| 56 | UCU | 855.1 | 854.9 | 853.9 | UCU UCU U |
| 57 | UGA | 918.2 | 917.9 | 916.9 | UGA UGA C |
| 58 | UGC | 894.2 | 893.9 | 892.9 | UGC UGC C |
| 59 | UGG | 934.2 | 933.9 | 932.9 | UGG UGG U |
| 60 | UGU | 895.2 | 894.9 | 893.9 | UGU UGU C |
| 61 | UUA | 879.2 | 878.9 | 877.9 | UUA UUA U |
| 62 | UUC | 855.1 | 854.9 | 853.9 | C UUC UUC |
| 63 | UUG | 895.2 | 894.9 | 893.9 | G UUG UUG |
| 64 | UUU | 856.1 | 855.9 | 854.9 | CC UUU UUU |

**Table S6** Biolayer interferometry data showing anticodon loops binding to biotin-labelled ssRNA. -1 and +1 indicate the residue before and after the codon of interest. $K_D$, equilibrium dissociation constant in µM (highlighted with a blue font); $k_{off}$, dissociation constant in $s^{-1}$; $k_{on}$, association constant in $M^{-1}s^{-1}$. M, mean; SD, standard deviation; CV, coefficient of variation; N, number of replicates; /, no binding detected. Bases in red highlight the residues which may form four base pairings. Values of $K_D$ <10 µM were rounded to one decimal place; values of $K_D$ in 10–50 µM were rounded to the nearest 1; values of $K_D$ in 50–150 µM were rounded to the nearest 5; values of $K_D$ in 150–1000 µM were rounded to the nearest 10; values of $K_D$ >1000 µM were rounded to the nearest 100. Conditions: 0–800 µM anticodon loop, 100 mM NaCl, 100 mM $MgCl_2$, 50 mM HEPES, pH 7.2, 20 ºC.

Family Box Codons:

| No. | Position | | | | | Codon | $K_D$ | | | $k_{off}$ | | | $k_{on}$ | | | N |
|---|---|---|---|---|---|---|---|---|---|---|---|---|---|---|---|---|
| | -1 | 1 | 2 | 3 | +1 | | M | SD | CV | M | SD | CV | M | SD | CV | |
| 1 | G | **A** | **C** | **A** | G | **ACA** | **150** | 39 | 26% | 0.798 | 0.150 | 19% | 5424 | 1046 | 19% | 4 |
| 1 | G | **A** | **C** | **A** | C | **ACA** | **160** | 10 | 6% | 0.456 | 0.017 | 4% | 2943 | 282 | 10% | 4 |
| 2 | C | **A** | **C** | **C** | C | **ACC** | **13** | 2 | 16% | 0.273 | 0.037 | 14% | 21478 | 4573 | 21% | 5 |
| 2 | A | **A** | **C** | **C** | U | **ACC** | **40** | 0.5 | 1% | 0.335 | 0.007 | 2% | 8366 | 168 | 2% | 4 |
| 3 | G | **A** | **C** | **G** | C | **ACG** | **110** | 4.8 | 4% | 0.802 | 0.064 | 8% | 7327 | 651 | 9% | 4 |
| 3 | A | **A** | **C** | **G** | G | **ACG** | **145** | 1.9 | 1% | 0.504 | 0.009 | 2% | 3491 | 66 | 2% | 4 |
| 4 | A | **A** | **C** | **U** | C | **ACU** | **530** | 47 | 9% | 1.283 | 0.167 | 13% | 2402 | 268 | 11% | 4 |
| 5 | U | **C** | **C** | **A** | U | **CCA** | **32** | 2.0 | 6% | 0.203 | 0.020 | 10% | 6326 | 262 | 4% | 4 |
| 5 | A | **C** | **C** | **A** | G | **CCA** | **75** | 12 | 16% | 0.642 | 0.051 | 8% | 8397 | 1127 | 13% | 4 |
| 6 | G | **C** | **C** | **C** | U | **CCC** | **12** | 2.2 | 18% | 0.202 | 0.021 | 10% | 17103 | 1742 | 10% | 3 |
| 6 | A | **C** | **C** | **C** | U | **CCC** | **15** | 5.3 | 35% | 0.298 | 0.028 | 9% | 20226 | 5481 | 27% | 5 |
| 6 | A | **C** | **C** | **C** | G | **CCC** | **16** | 1.4 | 9% | 0.120 | 0.014 | 12% | 7692 | 1362 | 18% | 4 |
| 7 | A | **C** | **C** | **G** | U | **CCG** | **65** | 11 | 17% | 0.421 | 0.067 | 16% | 6820 | 1241 | 18% | 4 |
| 7 | G | **C** | **C** | **G** | U | **CCG** | **85** | 16 | 19% | 0.410 | 0.095 | 23% | 4884 | 1488 | 30% | 4 |
| 8 | C | **C** | **C** | **U** | C | **CCU** | **230** | 26 | 11% | 0.640 | 0.056 | 9% | 2778 | 218 | 8% | 4 |
| 9 | G | **C** | **G** | **A** | C | **CGA** | **27** | 3.1 | 11% | 0.397 | 0.010 | 3% | 14643 | 2049 | 14% | 3 |
| 9 | A | **C** | **G** | **A** | G | **CGA** | **45** | 11.2 | 25% | 0.401 | 0.053 | 13% | 9968 | 3022 | 30% | 5 |
| 10 | C | **C** | **G** | **C** | C | **CGC** | **34** | 2.7 | 8% | 0.553 | 0.058 | 10% | 16425 | 2998 | 18% | 4 |
| 10 | C | **C** | **G** | **C** | C | **CGC** | **84** | 13 | 15% | 0.668 | 0.032 | 5% | 8046 | 1085 | 13% | 4 |
| 10 | C | **C** | **G** | **C** | C | **CGC** | **160** | 19 | 12% | 0.504 | 0.071 | 14% | 3150 | 223 | 7% | 5 |
| 11 | A | **C** | **G** | **G** | C | **CGG** | **80** | 11 | 14% | 1.059 | 0.110 | 10% | 12987 | 841 | 6% | 3 |
| 11 | A | **C** | **G** | **G** | G | **CGG** | **90** | 19 | 21% | 0.973 | 0.147 | 15% | 10774 | 909 | 8% | 4 |
| 12 | C | **C** | **G** | **U** | C | **CGU** | **75** | 18 | 23% | 0.466 | 0.131 | 28% | 6364 | 538 | 8% | 4 |
| 13 | G | **C** | **U** | **A** | C | **CUA** | **/** | / | / | **/** | / | / | **/** | / | / | **/** |
| 14 | C | **C** | **U** | **C** | C | **CUC** | **65** | 5.8 | 9% | 0.322 | 0.028 | 9% | 4861 | 179 | 4% | 4 |
| 15 | A | **C** | **U** | **G** | U | **CUG** | **730** | 216 | 30% | 0.644 | 0.088 | 14% | 937 | 214 | 23% | 5 |
| 16 | G | **C** | **U** | **U** | C | **CUU** | **150** | 34 | 23% | 0.169 | 0.010 | 6% | 1177 | 228 | 19% | 4 |
| 17 | A | **G** | **C** | **A** | C | **GCA** | **95** | 19 | 20% | 0.513 | 0.108 | 21% | 5353 | 89 | 2% | 3 |
| 18 | A | **G** | **C** | **C** | C | **GCC** | **15** | 2.4 | 16% | 0.312 | 0.037 | 12% | 21595 | 2269 | 11% | 4 |
| 18 | C | **G** | **C** | **C** | C | **GCC** | **15** | 1.1 | 7% | 0.134 | 0.016 | 12% | 9418 | 1461 | 16% | 4 |
| 18 | G | **G** | **C** | **C** | G | **GCC** | **22** | 1.7 | 8% | 0.131 | 0.026 | 20% | 5914 | 779 | 13% | 3 |
| 19 | A | **G** | **C** | **G** | A | **GCG** | **22** | 4.0 | 19% | 0.033 | 0.013 | 38% | 1500 | 343 | 23% | 3 |
| 19 | A | **G** | **C** | **G** | U | **GCG** | **38** | 0.8 | 2% | 0.425 | 0.012 | 3% | 11208 | 219 | 2% | 3 |
| 20 | G | **G** | **C** | **U** | C | **GCU** | **120** | 11 | 9% | 0.747 | 0.004 | 1% | 6507 | 633 | 10% | 3 |
| 21 | A | **G** | **G** | **A** | C | **GGA** | **120** | 4.1 | 3% | 0.556 | 0.034 | 6% | 4755 | 226 | 5% | 3 |
| 21 | A | **G** | **G** | **A** | U | **GGA** | **140** | 15 | 10% | 0.785 | 0.100 | 13% | 5731 | 1267 | 22% | 3 |
| 22 | A | **G** | **G** | **C** | C | **GGC** | **50** | 5.7 | 11% | 0.427 | 0.018 | 4% | 8551 | 1106 | 13% | 3 |
| 22 | G | **G** | **G** | **C** | U | **GGC** | **50** | 12 | 23% | 0.136 | 0.013 | 10% | 2713 | 327 | 12% | 3 |
| 22 | G | **G** | **G** | **C** | U | **GGC** | **65** | 8.7 | 13% | 0.092 | 0.028 | 30% | 1367 | 219 | 16% | 3 |
| 22 | C | **G** | **G** | **C** | C | **GGC** | **75** | 3.1 | 4% | 0.714 | 0.025 | 3% | 9663 | 394 | 4% | 4 |
| 23 | A | **G** | **G** | **G** | C | **GGG** | **35** | 2.5 | 7% | 0.418 | 0.040 | 10% | 11890 | 1929 | 16% | 3 |
| 24 | U | **G** | **G** | **U** | C | **GGU** | **34** | 0.5 | 2% | 0.436 | 0.018 | 4% | 12993 | 335 | 3% | 3 |
| 24 | A | **G** | **G** | **U** | C | **GGU** | **105** | 47 | 45% | 0.804 | 0.092 | 11% | 8424 | 2432 | 29% | 4 |
| 24 | A | **G** | **G** | **U** | C | **GGU** | **120** | 13.1 | 11% | 0.624 | 0.018 | 3% | 5290 | 732 | 14% | 5 |

| | | | | | | | | | | | | | | | | |
|---|---|---|---|---|---|---|---|---|---|---|---|---|---|---|---|---|
| 25 | A | **G** | **U** | **A** | U | **GUA** | **400** | 78 | 19% | 0.666 | 0.192 | 29% | 1654 | 234 | 14% | 4 |
| 26 | C | **G** | **U** | **C** | C | **GUC** | **5.5** | 0.5 | 9% | 0.155 | 0.018 | 12% | 28057 | 931 | 3% | 3 |
| 26 | A | **G** | **U** | **C** | A | **GUC** | **17** | 0.6 | 4% | 0.308 | 0.016 | 5% | 17685 | 1364 | 8% | 4 |
| 26 | A | **G** | **U** | **C** | C | **GUC** | **46** | 7.2 | 16% | 0.352 | 0.030 | 9% | 7756 | 847 | 11% | 5 |
| 26 | G | **G** | **U** | **C** | C | **GUC** | **65** | 2.9 | 5% | 0.491 | 0.011 | 2% | 7317 | 430 | 6% | 5 |
| 27 | A | **G** | **U** | **G** | C | **GUG** | **160** | 28 | 18% | 0.900 | 0.121 | 13% | 5813 | 1340 | 23% | 4 |
| 28 | A | **G** | **U** | **U** | U | **GUU** | **470** | 132 | 28% | 1.246 | 0.196 | 16% | 2717 | 504 | 19% | 3 |
| 29 | G | **U** | **C** | **A** | C | **UCA** | **280** | 60.9 | 22% | 0.117 | 0.017 | 15% | 435 | 91 | 21% | 5 |
| 30 | A | **U** | **C** | **C** | C | **UCC** | **73** | 8.9 | 12% | 0.501 | 0.008 | 2% | 6897 | 855 | 12% | 4 |
| 30 | C | **U** | **C** | **C** | G | **UCC** | **75** | 7.6 | 10% | 0.470 | 0.025 | 5% | 6439 | 391 | 6% | 4 |
| 30 | G | **U** | **C** | **C** | C | **UCC** | **77** | 24 | 32% | 0.768 | 0.213 | 28% | 10124 | 723 | 7% | 4 |
| 30 | C | **U** | **C** | **C** | C | **UCC** | **80** | 1.8 | 2% | 0.567 | 0.009 | 2% | 7025 | 196 | 3% | 4 |
| 31 | A | **U** | **C** | **G** | C | **UCG** | **310** | 31.8 | 10% | 0.272 | 0.015 | 6% | 903 | 147 | 16% | 4 |
| 32 | G | **U** | **C** | **U** | U | **UCU** | **260** | 71 | 27% | 0.152 | 0.036 | 24% | 606 | 213 | 35% | 4 |

**Table S6** continued. Split-box codons:

| No. | Position | | | | | Codon | $K_D$ | | | $k_{off}$ | | | $k_{on}$ | | | N |
|---|---|---|---|---|---|---|---|---|---|---|---|---|---|---|---|---|
| | -1 | 1 | 2 | 3 | +1 | | M | SD | CV | M | SD | CV | M | SD | CV | |
| 1 | G | **A** | **A** | **A** | C | AAA | / | / | / | / | / | / | / | / | / | / |
| 2 | C | **A** | **A** | **C** | U | AAC | **860** | 212 | 25% | 1.315 | 0.062 | 5% | 1596 | 409 | 26% | 3 |
| 3 | G | **A** | **A** | **G** | G | AAG | **1400** | 311 | 22% | 1.713 | 0.457 | 27% | 1248 | 334 | 27% | 3 |
| 4 | A | **A** | **A** | **U** | U | AAU | / | / | / | / | / | / | / | / | / | / |
| 5 | A | **A** | **G** | **A** | C | AGA | **1000** | 203 | 20% | 1.635 | 0.196 | 12% | 1591 | 181 | 11% | 3 |
| 6 | C | **A** | **G** | **C** | G | AGC | **115** | 9.8 | 9% | 0.483 | 0.039 | 8% | 4214 | 288 | 7% | 4 |
| 6 | A | **A** | **G** | **C** | C | AGC | **250** | 32 | 13% | 0.528 | 0.077 | 15% | 2109 | 143 | 7% | 3 |
| 7 | A | **A** | **G** | **G** | C | AGG | **85** | 19 | 22% | 0.603 | 0.081 | 13% | 7361 | 1046 | 14% | 3 |
| 8 | G | **A** | **G** | **U** | U | AGU | **170** | 33 | 20% | 1.379 | 0.158 | 11% | 8004 | 803 | 10% | 3 |
| 9 | G | **A** | **U** | **A** | G | AUA | **420** | 44 | 10% | 1.559 | 0.173 | 11% | 3706 | 99 | 3% | 3 |
| 10 | G | **A** | **U** | **C** | G | AUC | **1000** | 240 | 24% | 1.224 | 0.139 | 11% | 1226 | 261 | 21% | 4 |
| 11 | G | **A** | **U** | **G** | G | AUG | **330** | 32 | 10% | 0.810 | 0.078 | 10% | 2461 | 24 | 1% | 3 |
| 12 | A | **A** | **U** | **U** | A | AUU | / | / | / | / | / | / | / | / | / | / |
| 13 | G | **C** | **A** | **A** | C | CAA | / | / | / | / | / | / | / | / | / | / |
| 14 | G | **C** | **A** | **C** | G | CAC | **230** | 50 | 22% | 0.315 | 0.040 | 13% | 1453 | 411 | 28% | 3 |
| 15 | C | **C** | **A** | **G** | C | CAG | **190** | 14 | 7% | 0.244 | 0.049 | 20% | 1303 | 352 | 27% | 4 |
| 15 | C | **C** | **A** | **G** | C | CAG | **351** | 95 | 27% | 0.403 | 0.091 | 23% | 1161 | 151 | 13% | 3 |
| 16 | C | **C** | **A** | **U** | C | CAU | **450** | 73 | 16% | 0.041 | 0.006 | 14% | 92 | 17 | 19% | 5 |
| 16 | C | **C** | **A** | **U** | U | CAU | **570** | 33 | 6% | 0.095 | 0.004 | 5% | 168 | 7.8 | 5% | 3 |
| 17 | C | **G** | **A** | **A** | C | GAA | **750** | 82 | 11% | 1.038 | 0.251 | 24% | 1387 | 304 | 22% | 3 |
| 17 | C | **G** | **A** | **A** | U | GAA | **810** | 160 | 20% | 0.933 | 0.072 | 8% | 1169 | 151 | 13% | 3 |
| 18 | A | **G** | **A** | **C** | G | GAC | **170** | 10 | 6% | 0.410 | 0.040 | 10% | 2443 | 99 | 4% | 3 |
| 18 | A | **G** | **A** | **C** | G | GAC | **240** | 27 | 11% | 0.668 | 0.008 | 1% | 2793 | 331 | 12% | 4 |
| 18 | G | **G** | **A** | **C** | G | GAC | **250** | 35 | 14% | 0.658 | 0.077 | 12% | 2474 | 400 | 16% | 5 |
| 18 | C | **G** | **A** | **C** | G | GAC | **270** | 37 | 14% | 0.505 | 0.079 | 16% | 1898 | 32 | 2% | 3 |
| 19 | A | **G** | **A** | **G** | U | GAG | **140** | 6.9 | 5% | 0.736 | 0.082 | 11% | 5139 | 322 | 6% | 3 |
| 20 | A | **G** | **A** | **U** | G | GAU | **190** | 32 | 17% | 0.761 | 0.094 | 12% | 3934 | 357 | 9% | 3 |
| 20 | G | **G** | **A** | **U** | C | GAU | **190** | 35 | 18% | 0.231 | 0.034 | 15% | 1244 | 99 | 8% | 3 |
| 21 | A | **U** | **A** | **A** | G | UAA | / | / | / | / | / | / | / | / | / | / |
| 22 | U | **U** | **A** | **C** | C | UAC | **1500** | 188 | 13% | 0.332 | 0.100 | 30% | 230 | 65 | 28% | 4 |
| 23 | A | **U** | **A** | **G** | C | UAG | **210** | 47 | 23% | 0.424 | 0.110 | 26% | 2060 | 60 | 3% | 3 |
| 24 | G | **U** | **A** | **U** | C | UAU | / | / | / | / | / | / | / | / | / | / |
| 25 | A | **U** | **G** | **A** | U | UGA | / | / | / | / | / | / | / | / | / | / |
| 26 | G | **U** | **G** | **C** | U | UGC | / | / | / | / | / | / | / | / | / | / |
| 27 | A | **U** | **G** | **G** | U | UGG | **370** | 6 | 2% | 0.473 | 0.002 | 0% | 1263 | 27 | 2% | 3 |
| 28 | C | **U** | **G** | **U** | U | UGU | / | / | / | / | / | / | / | / | / | / |
| 29 | G | **U** | **U** | **A** | C | UUA | / | / | / | / | / | / | / | / | / | / |
| 30 | C | **U** | **U** | **C** | C | UUC | **470** | 33 | 7% | 0.659 | 0.009 | 1% | 1405 | 124 | 9% | 3 |
| 31 | G | **U** | **U** | **G** | C | UUG | **240** | 88 | 37% | 0.292 | 0.092 | 31% | 1245 | 90 | 7% | 3 |
| 32 | G | **U** | **U** | **U** | C | UUU | / | / | / | / | / | / | / | / | / | / |

**Table S7** Isothermal titration calorimetry raw data showing anticodon loops binding to ssRNA containing a single RNA triplet (light green) or two identical contiguous RNA triplets (dark green). $K_D$, equilibrium dissociation constant in µM; R, the mole ratio of the loop bound to ssRNA; dH, enthalpy change in kcal/mol; dG, free Gibbs energy change in kcal/mol. Bases in red highlight the residues which may form four base pairing. Conditions: 40 µM ssRNA in cell, 800 µM anticodon loop in syringe, 100 mM NaCl, 100 mM $MgCl_2$, 50 mM HEPES, pH 7.2, 10 ºC.

| Codon | Rep | Cell | Syringe | R | $K_D$ µM | dH kcal/mol | dG kcal/mol |
|---|---|---|---|---|---|---|---|
| **AAA** | 1 | AAA | GUCGC CU UUU AA GCGAC | / | / | / | / |
| | 2 | AAA | GUCGC CU UUU AA GCGAC | / | / | / | / |
| | 1 | AAA AAA C | GUCGC CU UUU AA GCGAC | / | / | / | / |
| | 2 | AAA AAA C | GUCGC CU UUU AA GCGAC | / | / | / | / |
| **AAC** | 1 | AAC | GUCGC CU GUU AA GCGAC | / | / | / | / |
| | 2 | AAC | GUCGC CU GUU AA GCGAC | / | / | / | / |
| | 1 | AAC AAC C | GUCGC CU GUU AA GCGAC | 1.37 | 8.47 | -19.60 | -6.57 |
| | 2 | AAC AAC C | GUCGC CU GUU AA GCGAC | 1.31 | 9.38 | -20.10 | -6.52 |
| **AAG** | 1 | AAG | GUCGC CU CUU AA GCGAC | / | / | / | / |
| | 2 | AAG | GUCGC CU CUU AA GCGAC | / | / | / | / |
| | 1 | AAG AAG U | GUCGC CU CUU AA GCGAC | 1.20 | 11.50 | -22.10 | -6.40 |
| | 2 | AAG AAG U | GUCGC CU CUU AA GCGAC | 1.21 | 11.80 | -22.00 | -6.39 |
| **AAU** | 1 | AAU | GUCGC CU AUU AA GCGAC | / | / | / | / |
| | 2 | AAU | GUCGC CU AUU AA GCGAC | / | / | / | / |
| | 1 | AAU AAU U | GUCGC CU AUU AA GCGAC | / | / | / | / |
| | 2 | AAU AAU U | GUCGC CU AUU AA GCGAC | / | / | / | / |
| **UAA** | 1 | UAA | GUCGC CU UUA AA GCGAC | / | / | / | / |
| | 2 | UAA | GUCGC CU UUA AA GCGAC | / | / | / | / |
| | 1 | UAA UAA U | GUCGC CU UUA AA GCGAC | / | / | / | / |
| | 2 | UAA UAA U | GUCGC CU UUA AA GCGAC | / | / | / | / |
| **UAC** | 1 | UAC | GUCGC CU GUA AA GCGAC | / | / | / | / |
| | 2 | UAC | GUCGC CU GUA AA GCGAC | / | / | / | / |
| | 1 | UAC UAC CC | GUCGC CU GUA AA GCGAC | / | / | / | / |
| | 2 | UAC UAC CC | GUCGC CU GUA AA GCGAC | / | / | / | / |
| **UAG** | 1 | UAG | GUCGC CU CUA AA GCGAC | / | / | / | / |
| | 2 | UAG | GUCGC CU CUA AA GCGAC | / | / | / | / |
| | 1 | G UAG UAG | GUCGC CU CUA AA GCGAC | / | / | / | / |
| | 2 | G UAG UAG | GUCGC CU CUA AA GCGAC | / | / | / | / |
| **UAU** | 1 | UAU | GUCGC CU AUA AA GCGAC | / | / | / | / |
| | 2 | UAU | GUCGC CU AUA AA GCGAC | / | / | / | / |
| | 1 | AU UAU UAU | GUCGC CU AUA AA GCGAC | / | / | / | / |
| | 2 | AU UAU UAU | GUCGC CU AUA AA GCGAC | / | / | / | / |
| **AGA** | 1 | AGA | GUCGC CU UCU AA GCGAC | / | / | / | / |
| | 2 | AGA | GUCGC CU UCU AA GCGAC | / | / | / | / |
| | 1 | AGA AGA U | GUCGC CU UCU AA GCGAC | 1.48 | 15.30 | -14.90 | -6.25 |
| | 2 | AGA AGA U | GUCGC CU UCU AA GCGAC | 1.54 | 15.70 | -15.00 | -6.23 |
| **AGC** | 1 | AGC | GUCGC CU GCU AA GCGAC | / | / | / | / |
| | 2 | AGC | GUCGC CU GCU AA GCGAC | / | / | / | / |
| | 1 | C AGC AGC | GUCGC CU GCU AA GCGAC | 1.76 | 3.37 | -7.46 | -7.10 |
| | 2 | C AGC AGC | GUCGC CU GCU AA GCGAC | 1.82 | 3.55 | -8.13 | -7.06 |
| **AGG** | 1 | AGG | GUCGC CU CCU AA GCGAC | 1.05 | 5.89 | -21.30 | -6.78 |
| | 2 | AGG | GUCGC CU CCU AA GCGAC | 1.12 | 4.73 | -19.80 | -6.91 |
| | 1 | AGG AGG U | GUCGC CU CCU AA GCGAC | 0.63 | 2.66 | -37.20 | -7.22 |
| | 2 | AGG AGG U | GUCGC CU CCU AA GCGAC | 0.62 | 1.93 | -32.80 | -7.41 |
| **AGU** | 1 | AGU | GUCGC CU ACU AA GCGAC | / | / | / | / |
| | 2 | AGU | GUCGC CU ACU AA GCGAC | / | / | / | / |
| | 1 | AGU AGU U | GUCGC CU ACU AA GCGAC | 1.17 | 6.78 | -21.10 | -6.70 |
| | 2 | AGU AGU U | GUCGC CU ACU AA GCGAC | 1.33 | 6.92 | -21.30 | -6.69 |
| **AUA** | 1 | AUA | GUCGC CU UAU AA GCGAC | / | / | / | / |

| | | | | | | | |
|---|---|---|---|---|---|---|---|
| | 2 | AUA | GUCGC CU UAU AA GCGAC | / | / | / | / |
| | 1 | AUA AUA U | GUCGC CU UAU AA GCGAC | / | / | / | / |
| | 2 | AUA AUA U | GUCGC CU UAU AA GCGAC | / | / | / | / |
| **AUC** | 1 | AUC | GUCGC CU GAU AA GCGAC | / | / | / | / |
| | 2 | AUC | GUCGC CU GAU AA GCGAC | / | / | / | / |
| | 1 | C AUC AUC | GUCGC CU GAU AA GCGAC | 1.15 | 8.55 | -16.10 | -6.57 |
| | 2 | C AUC AUC | GUCGC CU GAU AA GCGAC | 1.34 | 8.89 | -17.70 | -6.55 |
| **AUG** | 1 | AUG | GUCGC CU CAU AA GCGAC | / | / | / | / |
| | 2 | AUG | GUCGC CU CAU AA GCGAC | / | / | / | / |
| | 1 | G AUG AUG | GUCGC CU CAU AA GCGAC | 1.30 | 18.70 | -14.70 | -6.13 |
| | 2 | G AUG AUG | GUCGC CU CAU AA GCGAC | 1.10 | 16.90 | -14.20 | -6.19 |
| **AUU** | 1 | AUU | GUCGC CU AAU AA GCGAC | / | / | / | / |
| | 2 | AUU | GUCGC CU AAU AA GCGAC | / | / | / | / |
| | 1 | AUU AUU AU | GUCGC CU AAU AA GCGAC | / | / | / | / |
| | 2 | AUU AUU AU | GUCGC CU AAU AA GCGAC | / | / | / | / |
| **CAA** | 1 | CAA | GUCGC CU UUG AA GCGAC | / | / | / | / |
| | 2 | CAA | GUCGC CU UUG AA GCGAC | / | / | / | / |
| | 1 | CAA CAA UU | GUCGC CU UUG AA GCGAC | / | / | / | / |
| | 2 | CAA CAA UU | GUCGC CU UUG AA GCGAC | / | / | / | / |
| **CAC** | 1 | CAC | GUCGC CU GUG AA GCGAC | / | / | / | / |
| | 2 | CAC | GUCGC CU GUG AA GCGAC | / | / | / | / |
| | 1 | CAC CAC C | GUCGC CU GUG AA GCGAC | / | / | / | / |
| | 2 | CAC CAC C | GUCGC CU GUG AA GCGAC | / | / | / | / |
| **CAG** | 1 | CAG | GUCGC CU CUG AA GCGAC | / | / | / | / |
| | 2 | CAG | GUCGC CU CUG AA GCGAC | / | / | / | / |
| | 1 | CAG CAG C | GUCGC CU CUG AA GCGAC | / | / | / | / |
| | 2 | CAG CAG C | GUCGC CU CUG AA GCGAC | / | / | / | / |
| **CAU** | 1 | CAU | GUCGC CU UAG AA GCGAC | / | / | / | / |
| | 2 | CAU | GUCGC CU UAG AA GCGAC | / | / | / | / |
| | 1 | CAU CAU U | GUCGC CU UAG AA GCGAC | / | / | / | / |
| | 2 | CAU CAU U | GUCGC CU UAG AA GCGAC | / | / | / | / |
| **GAA** | 1 | GAA | GUCGC CU UUC AA GCGAC | / | / | / | / |
| | 2 | GAA | GUCGC CU UUC AA GCGAC | / | / | / | / |
| | 1 | GAA GAA CU | GUCGC CU UUC AA GCGAC | 1.69 | 14.00 | -12.40 | -6.30 |
| | 2 | GAA GAA CU | GUCGC CU UUC AA GCGAC | 1.70 | 13.10 | -11.10 | -6.33 |
| **GAC** | 1 | GAC | GUCGC CU GUC AA GCGAC | / | / | / | / |
| | 2 | GAC | GUCGC CU GUC AA GCGAC | / | / | / | / |
| | 1 | C GAC GAC | GUCGC CU GUC AA GCGAC | / | / | / | / |
| | 2 | C GAC GAC | GUCGC CU GUC AA GCGAC | / | / | / | / |
| **GAG** | 1 | GAG | GUCGC CU CUC AA GCGAC | 1.26 | 6.90 | -18.00 | -6.69 |
| | 2 | GAG | GUCGC CU CUC AA GCGAC | 1.36 | 7.41 | -17.60 | -6.65 |
| | 1 | GAG GAG U | GUCGC CU CUC AA GCGAC | 0.73 | 1.15 | -33.90 | -7.70 |
| | 2 | GAG GAG U | GUCGC CU CUC AA GCGAC | 0.79 | 0.55 | -27.60 | -8.12 |
| **GAU** | 1 | GAU | GUCGC CU AUC AA GCGAC | / | / | / | / |
| | 2 | GAU | GUCGC CU AUC AA GCGAC | / | / | / | / |
| | 1 | GAU GAU G | GUCGC CU AUC AA GCGAC | 1.13 | 12.50 | -24.20 | -6.36 |
| | 2 | GAU GAU G | GUCGC CU AUC AA GCGAC | 1.18 | 11.90 | -23.60 | -6.39 |
| **UGA** | 1 | UGA | GUCGC CU UCA AA GCGAC | / | / | / | / |
| | 2 | UGA | GUCGC CU UCA AA GCGAC | / | / | / | / |
| | 1 | UGA UGA C | GUCGC CU UCA AA GCGAC | / | / | / | / |
| | 2 | UGA UGA C | GUCGC CU UCA AA GCGAC | / | / | / | / |
| **UGC** | 1 | UGC | GUCGC CU GCA AA GCGAC | / | / | / | / |
| | 2 | UGC | GUCGC CU GCA AA GCGAC | / | / | / | / |
| | 1 | UGC UGC C | GUCGC CU GCA AA GCGAC | / | / | / | / |
| | 2 | UGG UGG U | GUCGC CU GCA AA GCGAC | / | / | / | / |
| **UGG** | 1 | UGG | GUCGC CU CCA AA GCGAC | / | / | / | / |
| | 2 | UGG | GUCGC CU CCA AA GCGAC | / | / | / | / |
| | 1 | UGG UGG U | GUCGC CU CCA AA GCGAC | / | / | / | / |
| | 2 | UGG UGG U | GUCGC CU CCA AA GCGAC | / | / | / | / |
| **UGU** | 1 | UGU | GUCGC CU ACA AA GCGAC | / | / | / | / |
| | 2 | UGU | GUCGC CU ACA AA GCGAC | / | / | / | / |

| | | | | | | | |
|---|---|---|---|---|---|---|---|
| | 1 | UGU UGU C | GUCGC CU ACA AA GCGAC | / | / | / | / |
| | 2 | UGU UGU C | GUCGC CU ACA AA GCGAC | / | / | / | / |
| **UUA** | 1 | UUA | GUCGC CU UAA AA GCGAC | / | / | / | / |
| | 2 | UUA | GUCGC CU UAA AA GCGAC | / | / | / | / |
| | 1 | A UUA UUA U | GUCGC CU UAA AA GCGAC | / | / | / | / |
| | 2 | A UUA UUA U | GUCGC CU UAA AA GCGAC | / | / | / | / |
| **UUC** | 1 | UUC | GUCGC CU GAA AA GCGAC | / | / | / | / |
| | 2 | UUC | GUCGC CU GAA AA GCGAC | / | / | / | / |
| | 1 | C UUC UUC | GUCGC CU GAA AA GCGAC | / | / | / | / |
| | 2 | C UUC UUC | GUCGC CU GAA AA GCGAC | / | / | / | / |
| **UUG** | 1 | UUG | GUCGC CU CAA AA GCGAC | / | / | / | / |
| | 2 | UUG | GUCGC CU CAA AA GCGAC | / | / | / | / |
| | 1 | G UUG UUG | GUCGC CU CAA AA GCGAC | / | / | / | / |
| | 2 | G UUG UUG | GUCGC CU CAA AA GCGAC | / | / | / | / |
| **UUU** | 1 | UUU | GUCGC CU AAA AA GCGAC | / | / | / | / |
| | 2 | UUU | GUCGC CU AAA AA GCGAC | / | / | / | / |
| | 1 | CC UUU UUU | GUCGC CU AAA AA GCGAC | / | / | / | / |
| | 2 | CC UUU UUU | GUCGC CU AAA AA GCGAC | / | / | / | / |
| **ACA** | 1 | ACA | GUCGC CU UGU AA GCGAC | / | / | / | / |
| | 2 | ACA | GUCGC CU UGU AA GCGAC | / | / | / | / |
| | 1 | ACA ACA U | GUCGC CU UGU AA GCGAC | 2.18 | 18.90 | -6.61 | -6.12 |
| | 2 | ACA ACA U | GUCGC CU UGU AA GCGAC | 2.28 | 18.40 | -6.65 | -6.14 |
| **ACC** | 1 | ACC | GUCGC CU GGU AA GCGAC | 1.40 | 5.72 | -13.70 | -6.80 |
| | 2 | ACC | GUCGC CU GGU AA GCGAC | 1.36 | 5.71 | -13.90 | -6.80 |
| | 1 | C ACC ACC | GUCGC CU GGU AA GCGAC | 2.01 | 3.78 | -19.00 | -7.03 |
| | 2 | C ACC ACC | GUCGC CU GGU AA GCGAC | 1.68 | 4.76 | -18.40 | -6.90 |
| **ACG** | 1 | ACG | GUCGC CU CGU AA GCGAC | 1.61 | 10.60 | -12.10 | -6.45 |
| | 2 | ACG | GUCGC CU CGU AA GCGAC | 1.53 | 12.40 | -10.50 | -6.36 |
| | 1 | ACG ACG | GUCGC CU CGU AA GCGAC | 0.92 | 4.35 | -20.60 | -6.95 |
| | 2 | ACG ACG | GUCGC CU CGU AA GCGAC | 1.27 | 7.74 | -19.10 | -6.63 |
| **ACU** | 1 | ACU | GUCGC CU AGU AA GCGAC | / | / | / | / |
| | 2 | ACU | GUCGC CU AGU AA GCGAC | / | / | / | / |
| | 1 | ACU ACU U | GUCGC CU AGU AA GCGAC | 2.22 | 13.20 | -11.60 | -6.32 |
| | 2 | ACU ACU U | GUCGC CU AGU AA GCGAC | 1.80 | 12.80 | -11.10 | -6.34 |
| **CGA** | 1 | CGA | GUCGC CU UCG AA GCGAC | / | / | / | / |
| | 2 | CGA | GUCGC CU UCG AA GCGAC | / | / | / | / |
| | 1 | CGA CGA C | GUCGC CU UCG AA GCGAC | / | / | / | / |
| | 2 | CGA CGA C | GUCGC CU UCG AA GCGAC | / | / | / | / |
| **CGC** | 1 | CGC | GUCGC CU GCG AA GCGAC | / | / | / | / |
| | 2 | CGC | GUCGC CU GCG AA GCGAC | / | / | / | / |
| | 1 | CGC CGC C | GUCGC CU GCG AA GCGAC | / | / | / | / |
| | 2 | CGC CGC C | GUCGC CU GCG AA GCGAC | / | / | / | / |
| **CGG** | 1 | CGG | GUCGC CU CCG AA GCGAC | 1.98 | 13.30 | -7.62 | -6.32 |
| | 2 | CGG | GUCGC CU CCG AA GCGAC | 1.68 | 14.90 | -8.83 | -6.25 |
| | 1 | CGG CGG | GUCGC CU CCG AA GCGAC | 2.51 | 7.03 | -5.36 | -6.69 |
| | 2 | CGG CGG | GUCGC CU CCG AA GCGAC | 2.45 | 6.22 | -5.28 | -6.75 |
| **CGU** | 1 | CGU | GUCGC CU ACG AA GCGAC | / | / | / | / |
| | 2 | CGU | GUCGC CU ACG AA GCGAC | / | / | / | / |
| | 1 | CCU CGU UU | GUCGC CU ACG AA GCGAC | 1.68 | 11.90 | -12.00 | -6.38 |
| | 2 | CCU CGU UU | GUCGC CU ACG AA GCGAC | 1.72 | 11.50 | -11.80 | -6.40 |
| | 1 | CGU CGU UU | GUCGC CU ACG AA GCGAC | 2.31 | 11.90 | -5.18 | -6.38 |
| | 2 | CGU CGU UU | GUCGC CU ACG AA GCGAC | 2.50 | 10.90 | -5.14 | -6.43 |
| **CUA** | 1 | CUA | GUCGC CU UAG AA GCGAC | / | / | / | / |
| | 2 | CUA | GUCGC CU UAG AA GCGAC | / | / | / | / |
| | 1 | CUA CUA C | GUCGC CU UAG AA GCGAC | / | / | / | / |
| | 2 | CUA CUA C | GUCGC CU UAG AA GCGAC | / | / | / | / |
| **CUC** | 1 | CUC | GUCGC CU GAG AA GCGAC | / | / | / | / |
| | 2 | CUC | GUCGC CU GAG AA GCGAC | / | / | / | / |
| | 1 | CUC CUC CC | GUCGC CU GAG AA GCGAC | 1.60 | 14.8 | -18.70 | -6.26 |
| | 2 | CUC CUC CC | GUCGC CU GAG AA GCGAC | 1.47 | 16.2 | -19.10 | -6.21 |
| **CUG** | 1 | CUG | GUCGC CU CAG AA GCGAC | / | / | / | / |

| | | | | | | | |
|---|---|---|---|---|---|---|---|
| | 2 | CUG | GUCGC CU CAG AA GCGAC | / | / | / | / |
| | 1 | G CUG CUG | GUCGC CU CAG AA GCGAC | / | / | / | / |
| | 2 | G CUG CUG | GUCGC CU CAG AA GCGAC | / | / | / | / |
| **CUU** | 1 | CUU | GUCGC CU AAG AA GCGAC | / | / | / | / |
| | 2 | CUU | GUCGC CU AAG AA GCGAC | / | / | / | / |
| | 1 | C CUU UUUU | GUCGC CU AAG AA GCGAC | / | / | / | / |
| | 2 | C CUU UUUU | GUCGC CU AAG AA GCGAC | / | / | / | / |
| | 1 | CUU CUU U | GUCGC CU AAG AA GCGAC | 2.20 | 23.60 | -10.20 | -6.00 |
| | 2 | CUU CUU U | GUCGC CU AAG AA GCGAC | 2.25 | 20.90 | -9.54 | -6.06 |
| **GGA** | 1 | GGA | GUCGC CU UCC AA GCGAC | 1.12 | 5.19 | -19.50 | -6.85 |
| | 2 | GGA | GUCGC CU UCC AA GCGAC | 1.16 | 5.42 | -19.30 | -6.82 |
| | 1 | GGA U | GUCGC CU UCC AA GCGAC | 1.01 | 3.93 | -21.50 | -7.01 |
| | 2 | GGA U | GUCGC CU UCC AA GCGAC | 1.02 | 3.74 | -21.90 | -7.04 |
| | 1 | GGA GGA U | GUCGC CU UCC AA GCGAC | 0.17 | 0.62 | -31.30 | -8.05 |
| | 2 | GGA GGA U | GUCGC CU UCC AA GCGAC | 0.16 | 0.58 | -28.70 | -8.10 |
| **GGC** | 1 | GGC | GUCGC CU GCC AA GCGAC | 1.71 | 1.88 | -9.87 | -7.42 |
| | 2 | GGC | GUCGC CU GCC AA GCGAC | 1.42 | 1.08 | -7.81 | -7.73 |
| | 1 | GGC GGC G | GUCGC CU GCC AA GCGAC | 1.26 | 2.05 | -3.77 | -7.65 |
| | 2 | GGC GGC G | GUCGC CU GCC AA GCGAC | 1.36 | 2.08 | -5.25 | -7.61 |
| **GGG** | 1 | GGG | GUCGC CU CCC AA GCGAC | 0.32 | 0.34 | -29.40 | -8.38 |
| | 2 | GGG | GUCGC CU CCC AA GCGAC | 0.33 | 0.27 | -27.30 | -8.51 |
| | 1 | GGG GGG | GUCGC CU CCC AA GCGAC | / | / | / | / |
| | 2 | GGG GGG | GUCGC CU CCC AA GCGAC | / | / | / | / |
| **GGU** | 1 | GGU | GUCGC CU ACC AA GCGAC | 0.93 | 4.23 | -22.40 | -6.97 |
| | 2 | GGU | GUCGC CU ACC AA GCGAC | 1.00 | 4.93 | -22.00 | -6.89 |
| | 1 | GGU GGU G | GUCGC CU ACC AA GCGAC | 0.04 | 5.42 | -80.00 | -6.83 |
| | 2 | GGU GGU G | GUCGC CU ACC AA GCGAC | 0.04 | 5.31 | -80.00 | -6.84 |
| **GUA** | 1 | GUA | GUCGC CU UAC AA GCGAC | / | / | / | / |
| | 2 | GUA | GUCGC CU UAC AA GCGAC | / | / | / | / |
| | 1 | GUA GUA G | GUCGC CU UAC AA GCGAC | 0.98 | 14.10 | -28.40 | -6.29 |
| | 2 | GUA GUA G | GUCGC CU UAC AA GCGAC | 0.80 | 16.30 | -23.60 | -6.20 |
| **GUC** | 1 | GUC | GUCGC CU GAC AA GCGAC | 1.54 | 5.55 | -15.60 | -6.81 |
| | 2 | GUC | GUCGC CU GAC AA GCGAC | 1.59 | 5.52 | -15.50 | -6.82 |
| | 1 | GUC GUC G | GUCGC CU GAC AA GCGAC | 2.16 | 1.41 | -14.10 | -7.59 |
| | 2 | GUC GUC G | GUCGC CU GAC AA GCGAC | 1.86 | 1.03 | -15.60 | -7.77 |
| **GUG** | 1 | GUG | GUCGC CU CAC AA GCGAC | 1.21 | 9.46 | -14.00 | -6.51 |
| | 2 | GUG | GUCGC CU CAC AA GCGAC | 1.30 | 9.95 | -13.80 | -6.49 |
| | 1 | G GUG GUG | GUCGC CU CAC AA GCGAC | 0.04 | 3.84 | -68.10 | -7.03 |
| | 2 | G GUG GUG | GUCGC CU CAC AA GCGAC | 0.04 | 3.75 | -63.30 | -7.04 |
| **GUU** | 1 | GUU | GUCGC CU AAC AA GCGAC | / | / | / | / |
| | 2 | GUU | GUCGC CU AAC AA GCGAC | / | / | / | / |
| | 1 | GUU GUU G | GUCGC CU AAC AA GCGAC | 0.64 | 7.44 | -36.30 | -6.65 |
| | 2 | GUU GUU G | GUCGC CU AAC AA GCGAC | 0.62 | 6.89 | -39.40 | -6.69 |
| **UCA** | 1 | UCA | GUCGC CU UGA AA GCGAC | / | / | / | / |
| | 2 | UCA | GUCGC CU UGA AA GCGAC | / | / | / | / |
| | 1 | UCA UCA U | GUCGC CU UGA AA GCGAC | / | / | / | / |
| | 2 | UCA UCA U | GUCGC CU UGA AA GCGAC | / | / | / | / |
| **UCC** | 1 | UCC | GUCGC CU GGA AA GCGAC | 2.60 | 8.55 | -4.54 | -6.57 |
| | 2 | UCC | GUCGC CU GGA AA GCGAC | 2.52 | 9.05 | -4.76 | -6.54 |
| | 1 | UCC UCC UU | GUCGC CU GGA AA GCGAC | 1.24 | 8.50 | -20.40 | -6.59 |
| | 2 | UCC UCC UU | GUCGC CU GGA AA GCGAC | 1.30 | 10.10 | -21.90 | -6.48 |
| **UCG** | 1 | UCG | GUCGC CU CGA AA GCGAC | / | / | / | / |
| | 2 | UCG | GUCGC CU CGA AA GCGAC | / | / | / | / |
| | 1 | CC UCG UUU | GUCGC CU CGA AA GCGAC | / | / | / | / |
| | 2 | CC UCG UUU | GUCGC CU CGA AA GCGAC | / | / | / | / |
| | 1 | G UCG UCG | GUCGC CU CGA AA GCGAC | / | / | / | / |
| | 2 | G UCG UCG | GUCGC CU CGA AA GCGAC | / | / | / | / |
| **UCU** | 1 | UCU | GUCGC CU AGA AA GCGAC | / | / | / | / |
| | 2 | UCU | GUCGC CU AGA AA GCGAC | / | / | / | / |
| | 1 | UCU UCU U | GUCGC CU AGA AA GCGAC | / | / | / | / |
| | 2 | UCU UCU U | GUCGC CU AGA AA GCGAC | / | / | / | / |

| | | | | | | | |
|---|---|---|---|---|---|---|---|
| **CCA** | 1 | CCA | GUCGC CU UGG AA GCGAC | / | / | / | / |
| | 2 | CCA | GUCGC CU UGG AA GCGAC | / | / | / | / |
| | 1 | CCA UUUUU | GUCGC CU UGG AA GCGAC | 1.46 | 18.30 | -10.60 | -6.14 |
| | 2 | CCA UUUUU | GUCGC CU UGG AA GCGAC | 1.46 | 18.10 | -10.50 | -6.15 |
| | 1 | CCA CCA UU | GUCGC CU UGG AA GCGAC | 1.95 | 22.60 | -13.50 | -6.02 |
| | 2 | CCA CCA UU | GUCGC CU UGG AA GCGAC | 1.91 | 22.40 | -13.30 | -6.03 |
| **CCC** | 1 | CCC | GUCGC CU GGG AA GCGAC | 1.28 | 2.77 | -20.60 | -7.21 |
| | 2 | CCC | GUCGC CU GGG AA GCGAC | 1.28 | 2.92 | -20.90 | -7.18 |
| | 1 | CCC CCC | GUCGC CU GGG AA GCGAC | 1.88 | 0.33 | -24.50 | -8.41 |
| | 2 | CCC CCC | GUCGC CU GGG AA GCGAC | 1.91 | 0.55 | -25.40 | -8.12 |
| | 1 | CCCCCCCC | GUCGC CU GGG AA GCGAC | 3.01 | 2.44 | -24.20 | -7.27 |
| | 2 | CCCCCCCC | GUCGC CU GGG AA GCGAC | 3.06 | 2.49 | -24.80 | -7.26 |
| **CCG** | 1 | CCG | GUCGC CU CGG AA GCGAC | 2.18 | 6.91 | -6.03 | -6.69 |
| | 2 | CCG | GUCGC CU CGG AA GCGAC | 1.95 | 7.66 | -6.23 | -6.63 |
| | 1 | G CCG CCG | GUCGC CU CGG AA GCGAC | / | / | / | / |
| | 2 | G CCG CCG | GUCGC CU CGG AA GCGAC | / | / | / | / |
| **CCU** | 1 | CCU | GUCGG CU AGG AA CCGAC | / | / | / | / |
| | 2 | CCU | GUCGG CU AGG AA CCGAC | / | / | / | / |
| | 1 | CCU U | GUCGG CU AGG AA CCGAC | 1.41 | 19.50 | -9.52 | -6.11 |
| | 2 | CCU U | GUCGG CU AGG AA CCGAC | 1.41 | 18.00 | -9.36 | -6.15 |
| | 1 | CCU CCU UU | GUCGG CU AGG AA CCGAC | 1.13 | 9.68 | -22.90 | -6.50 |
| | 2 | CCU CCU UU | GUCGG CU AGG AA CCGAC | 1.44 | 11.40 | -19.60 | -6.41 |
| **GCA** | 1 | GCA | GUCGG CU UGC AA CCGAC | / | / | / | / |
| | 2 | GCA | GUCGG CU UGC AA CCGAC | / | / | / | / |
| | 1 | GCA GCA UU | GUCGG CU UGC AA CCGAC | / | / | / | / |
| | 2 | GCA GCA UU | GUCGG CU UGC AA CCGAC | / | / | / | / |
| **GCC** | 1 | GCC | GUCGC CU GGC AA GCGAC | 1.42 | 0.66 | -15.80 | -8.02 |
| | 2 | GCC | GUCGC CU GGC AA GCGAC | 1.50 | 0.67 | -15.40 | -8.01 |
| | 1 | GCC GCC C | GUCGC CU GGC AA GCGAC | 2.76 | 0.42 | -12.50 | -8.27 |
| | 2 | GCC GCC C | GUCGC CU GGC AA GCGAC | 2.80 | 0.37 | -12.60 | -8.35 |
| **GCG** | 1 | GCG | GUCGC CU CGC AA GCGAC | 1.04 | 1.23 | -14.30 | -7.66 |
| | 2 | GCG | GUCGC CU CGC AA GCGAC | 0.89 | 0.95 | -15.10 | -7.81 |
| | 1 | G GCG GCG | GUCGC CU CGC AA GCGAC | 1.05 | 0.42 | -9.39 | -8.27 |
| | 2 | G GCG GCG | GUCGC CU CGC AA GCGAC | 1.21 | 0.48 | -9.25 | -8.20 |
| **GCU** | 1 | GCU | GUCGC CU AGC AA GCGAC | 1.34 | 6.76 | -16.20 | -6.70 |
| | 2 | GCU | GUCGC CU AGC AA GCGAC | 1.27 | 6.59 | -16.40 | -6.72 |
| | 1 | GCU GCU G | GUCGC CU AGC AA GCGAC | 1.47 | 0.16 | -24.20 | -8.82 |
| | 2 | GCU GCU G | GUCGC CU AGC AA GCGAC | 1.44 | 0.15 | -21.00 | -8.87 |
| **Loop-Loop** | GUCGC CU GAC AA GCGAC | | GUCGC CU GUU AA GCGAC | 0.63 | 1.28 | -21.3 | -7.64 |
| | GUCGC CU GUU AA GCGAC | | GUCGC CU GAC AA GCGAC | 0.86 | 3.14 | -27.6 | -7.13 |
| | GUCGC CU GGC AA GCGAC | | GUCGC CU GCC AA GCGAC | 0.81 | 0.99 | -16.5 | -7.78 |
| | GUCGC CU GGU AA GCGAC | | GUCGC CU GCC AA GCGAC | 1.38 | 1.73 | -13.5 | -7.47 |

**Table S8** Isothermal titration calorimetry data for anticodon loops binding to a ssRNA containing a single triplet. $K_D$, equilibrium dissociation constant in μM; R, the mole ratio of the loop to ssRNA; dH, enthalpy change in kcal/mol. Blocks in blue highlight the family-box (four-fold degenerate) codons. [a] hairpin forms self-dimer/duplex; [b] ssRNA forms G-quadruplex; [c] ssRNA forms self-dimer.

| | U | | | C | | | A | | | G | | | |
|---|---|---|---|---|---|---|---|---|---|---|---|---|---|
| | $K_D$ | R | dH | $K_D$ | R | dH | $K_D$ | R | dH | $K_D$ | R | dH | |
| U | / | | | / | | | / | | | / | | | U |
| | / | | | / | | | / | | | / | | | C |
| | / | | | / | | | /[a] | | | / | | | A |
| | / | | | / | | | / | | | / | | | G |
| C | / | | | / | | | / | | | /[c] | | | U |
| | / | | | 2.8 | 1.3 | -21 | / | | | /[c] | | | C |
| | / | | | / | | | / | | | /[a] | | | A |
| | / | | | / | | | / | | | 14 | 1.8[a] | -8.2 | G[a] |
| A | / | | | / | | | / | | | / | | | U |
| | / | | | 5.7 | 1.4 | -14 | / | | | / | | | C |
| | /[a] | | | / | | | / | | | / | | | A |
| | / | | | 12 | 1.6 | -11 | / | | | 5.3 | 1.1 | -21 | G |
| G | / | | | 6.7 | 1.3 | -16 | / | | | 4.6 | 1.0 | -22 | U |
| | 5.5 | 1.6 | -16 | 0.7 | 1.5 | -16 | / | | | 1.5 | 1.6 | -8.8 | C[a] |
| | / | | | /[a] | | | / | | | 5.3 | 1.1 | -19 | A |
| | 9.7 | 1.3 | -14 | 1.1 | 1.0 | -15 | 7.2 | 1.3 | -18 | 0.3 | 0.3[b] | -28 | G |

**Table S9** Isothermal titration calorimetry data for anticodon loops binding to a ssRNA containing two contiguous identical triplets. $K_D$ equilibrium dissociation constant in μM; R, the mole ratio of the loop to ssRNA; dH, enthalpy change in kcal/mol. Boxes in blue highlight the family-box (four-fold degenerate) codons. a) hairpin forms self-dimer/duplex; b) ssRNA forms G-quadruplex; c) ssRNA forms self-dimer; d) possible four base pairings.

| | U | | | C | | | A | | | G | | | |
|---|---|---|---|---|---|---|---|---|---|---|---|---|---|
| | $K_D$ | R | dH | $K_D$ | R | dH | $K_D$ | R | H | $K_D$ | R | dH | |
| U | / | | | / | | | / | | | / | | | U[d] |
| | / | | | 9.3 | 1.3 | -21 | / | | | /[c] | | | C |
| | / | | | / | | | /[a] | | | / | | | A |
| | / | | | / | | | / | | | / | | | G |
| C | 22 | 2.2 | -9.9 | 11 | 1.3 | -21 | / | | | 11 | 2.4 | -5.2 | U[d] |
| | 15 | 1.5 | -19 | 0.4 | 1.9 | -25 | / | | | /[c] | | | C |
| | / | | | 22 | 1.9 | -13 | / | | | /[a] | | | A |
| | / | | | / | | | / | | | 6.6 | 2.5[a] | -5.3 | G[a] |
| A[d] | / | | | 13 | 2.0 | -11 | / | | | 6.8 | 1.2 | -21 | U[d] |
| | 8.7 | 1.2 | -17 | 4.3 | 1.9 | -19 | 8.9 | 1.3 | -20 | 3.5 | 1.8 | -7.8 | C |
| | /[a] | | | 19 | 2.2 | -6.6 | / | | | 16 | 1.5 | -15 | A |
| | 18 | 1.2 | -14 | 6.1 | 1.1[c] | -20 | 12 | 1.2 | -22 | 2.3 | 0.6[b] | -35 | G |
| G | 7.2 | 0.6 | -38 | 0.2 | 1.5 | -23 | 12 | 1.2 | -24 | 5.4 | 0.04[b] | -80 | U[d] |
| | 1.2 | 2.0 | -15 | 0.4 | 2.8[c] | -13 | / | | | 1.3 | 2.1 | -4.5 | C[a] |
| | 15 | 0.9 | -26 | /[a] | | | 14 | 1.7 | -12 | 0.6 | 0.2[b] | -30 | A |
| | 3.8 | 0.04[b] | -66 | 0.5 | 1.1[b] | -9.3 | 0.9 | 0.8[b] | -31 | /[b] | | | G |

**Table S10** Biolayer interferometry data for two different anticodon loops binding to biotin-labeled ssRNA. -1 and +1 indicate the residue before and after the codon of interest. s, single loop binding to the ssRNA; sq, sequential binding, the second binding kinetics after the first codon is saturated; m, mixed two anticodon loops binding to ssRNA 5'-A UUC CAU C. $K_D$, equilibrium dissociation constant in μM; $k_{off}$, disociation constant in $s^{-1}$; $k_{on}$, association constant in $M^{-1}s^{-1}$. M, mean; SD, standard deviation; CV, coefficient of variation; N, number of replicates. Data in black indicate family box codons; blue indicate split box codons.

Conditions: 200 μM for the first family-box anticodon loop, 0-400 μM for the second anticodon loop or the single loop, 100 mM NaCl, 100 mM $MgCl_2$, 50 mM HEPES, pH 7.2, 20 °C.

| | Position | | | | | | | $K_D$ | | | $k_{off}$ | | | $k_{on}$ | | | N |
|---|---|---|---|---|---|---|---|---|---|---|---|---|---|---|---|---|---|
| | -1 | 1 | 2 | 3 | +1 | Codon | Aa | M | SD | CV | M | SD | CV | M | SD | CV | |
| s | A | G | C | C | C | GCC | Ala | **15** | 2.4 | 16% | 0.312 | 0.037 | 12% | 21595 | 2269 | 11% | 4 |
| sq | A | G | C | C | C | GCC | | **13** | 0.8 | 6% | 0.055 | 0.005 | 9% | 4154 | 222 | 5% | 5 |
| s | C | C | U | C | C | CUC | Leu | **66** | 5.8 | 9% | 0.322 | 0.028 | 9% | 4861 | 179 | 4% | 4 |
| sq | C | C | U | C | C | CUC | | **49** | 2.8 | 6% | 0.040 | 0.009 | 22% | 805 | 152 | 19% | 5 |
| | | | | | | 5'-A GCC CUC C | | | | | | | | | | | |
| s | A | C | C | C | U | CCC | Pro | **16** | 5.3 | 33% | 0.298 | 0.028 | 9% | 20226 | 5481 | 27% | 5 |
| sq | A | C | C | C | U | CCC | | **16** | 1.7 | 11% | 0.224 | 0.018 | 8% | 14368 | 692 | 5% | 5 |
| s | C | U | C | C | G | UCC | Ser | **73** | 7.6 | 10% | 0.470 | 0.025 | 5% | 6439 | 391 | 6% | 4 |
| sq | C | U | C | C | G | UCC | | **110** | 7.9 | 7% | 0.765 | 0.167 | 22% | 7027 | 1171 | 17% | 5 |
| | | | | | | 5'-A CCC UCC G | | | | | | | | | | | |
| s | A | A | C | G | G | ACG | Thr | **145** | 3.1 | 2% | 0.509 | 0.012 | 2% | 3550 | 146 | 4% | 5 |
| sq | A | A | C | G | G | ACG | | **135** | 11.9 | 9% | 0.17 | 0.014 | 9% | 1252 | 199 | 16% | 4 |
| s | G | G | U | C | C | GUC | Val | **67** | 2.9 | 4% | 0.491 | 0.011 | 2% | 7317 | 430 | 6% | 5 |
| sq | G | G | U | C | C | GUC | | **62** | 9.2 | 15% | 0.699 | 0.049 | 7% | 11506 | 1355 | 12% | 5 |
| | | | | | | 5'-A ACG GUC C | | | | | | | | | | | |
| s | A | C | G | A | G | CGA | Arg | **43** | 11.2 | 26% | 0.401 | 0.053 | 13% | 9968 | 3022 | 30% | 5 |
| sq | A | C | G | A | G | CGA | | **47** | 12.4 | 26% | 0.886 | 0.135 | 15% | 19730 | 5732 | 29% | 5 |
| s | A | G | G | U | C | GGU | Gly | **120** | 13.1 | 11% | 0.624 | 0.018 | 3% | 5290 | 732 | 14% | 5 |
| sq | A | G | G | U | C | GGU | | **105** | 9.1 | 9% | 0.567 | 0.020 | 4% | 5365 | 317 | 6% | 5 |
| | | | | | | 5'-A CGA GGU C | | | | | | | | | | | |
| s | A | G | A | C | G | GAC | Asp | **240** | 27 | 11% | 0.668 | 0.008 | 1% | 2793 | 331 | 12% | 4 |
| sq | A | G | A | C | G | GAC | | **580** | 51 | 9% | 1.650 | 0.012 | 1% | 2852 | 248 | 9% | 4 |
| s | C | G | C | C | C | GCC | Ala | **14** | 1.1 | 8% | 0.134 | 0.016 | 12% | 9418 | 1461 | 16% | 4 |
| sq | C | G | C | C | C | GCC | | **34** | 1.0 | 3% | 0.195 | 0.010 | 5% | 5787 | 227 | 4% | 5 |
| | | | | | | 5'-A GAC GCC C | | | | | | | | | | | |
| s | A | G | U | C | U | GUC | Val | **80** | 0.8 | 1% | 0.806 | 0.008 | 1% | 10021 | 169 | 2% | 4 |
| sq | A | G | U | C | U | GUC | | **240** | 33.6 | 14% | 0.862 | 0.071 | 8% | 3595 | 394 | 11% | 4 |
| s | C | U | A | C | C | UAC | Tyr | **3400** | 578 | 17% | 0.504 | 0.076 | 15% | 153 | 49 | 32% | 5 |
| sq | C | U | A | C | C | UAC | | **/** | / | / | / | / | / | / | / | / | / |
| | | | | | | 5'-A GAC GCC C | | | | | | | | | | | |
| s | C | **U** | **U** | **C** | C | UUC | Phe | **450** | 32 | 7% | 0.635 | 0.047 | 7% | 1404 | 124 | 9% | 3 |
| s | C | **C** | **A** | **U** | C | CAU | His | **180** | 2.8 | 2% | 0.031 | 0.001 | 5% | 169 | 7 | 4% | 3 |
| m | | | | | | mixed | | **190** | 9.0 | 5% | 0.024 | 0.001 | 5% | 127 | 10 | 8% | 5 |
| | | | | | | 5'-A UUC CAU C | | | | | | | | | | | |

**Table 11** Example of (A) a subset of anticodon loops that have some pairs which can bind each other (all ending in C) and (B) an orthogonal subset. A mismatch in the codon:anticodon pair leads to a considerable decrease in $K_D$ (Fig. S17). The codon:anticodon pair do not permit mismatch or wobble (WB) pairing, while loop-loop interaction allows WB pairing (Fig. S12). If the potentially paired anticodons are present in this anticodon set, (indicated as GGU[1], GGC[2], and GCC[3] in the Table A) the set is self-interacting and should be disregarded.
See https://github.com/nieseln/Codon-select.git for code.

(A)

| Aa | Codon | Anticodon | Possibly kissed anticodons 5'-3' | | | | | | |
|---|---|---|---|---|---|---|---|---|---|
| | | 3WC | 3WC | 2WC + 1 WB | | | 1WC + 2WB | | |
| Ser | **UCC** | GGA | UCC | UCU | UUC | | UUU | | |
| Leu | **CUC** | GAG | CUC | CUU | UUC | | UUU | | |
| Pro | **CCC** | GGG | CCC | CCU | CUC | UCC | UUC | UCU | CUU |
| Arg | **CGC** | GCG* | CGC | UGC | CGU | | UGU | | |
| Thr | **ACC** | GGU[1] | ACC | ACU | AUC | GCC[3] | AUU | GUC | GCU |
| Val | **GUC** | GAC | GUC | GUU | | | | | |
| Ala | **GCC** | GGC[2] | GCC[3] | GCU | GUC | | GUU | | |
| Gly | **GGC** | GCC[3] | GGC[2] | GGU[1] | | | | | |

(B)

| Aa | Codon | Anticodon | Possibly kissed anticodons 5'-3' | | | | | | |
|---|---|---|---|---|---|---|---|---|---|
| | | 3WC | 3WC | 2WC + 1 WB | | | 1WC + 2WB | | |
| Ser | **UCC** | GGA | UCC | UCU | UUC | | UUU | | |
| Leu | **CUC** | GAG | CUC | CUU | UUC | | UUU | | |
| Pro | **CCC** | GGG | CCC | CCU | CUC | UCC | UUC | UCU | CUU |
| Arg | **CGA** | UCG | CGA | UGA | CGG | | CGG | | |
| Thr | **ACG** | CGU | ACG | GCG | AUG | | GUG | | |
| Val | **GUC** | GAC | GUC | GUU | | | | | |
| Ala | **GCC** | GGC | GCC | GCU | GUC | | GUU | | |
| Gly | **GGU** | ACC | GGU | | | | | | |

**Table S12** Fourteen orthogonol family-box codon sets from exhaustive search. Codon sets including GGG were excluded to avoid issues due to quadruplex formation. Set 3 was chosen for sequential binding measurement.

| | Ser | Leu | Pro | Arg | Thr | Leu | Ala | Gly |
|---|---|---|---|---|---|---|---|---|
| 1 | UCC | CUC | CCC | CGA | ACG | GUC | GCU | GGU |
| 2 | UCC | CUC | CCC | CGA | ACG | GUC | GCU | GGC |
| **3** | **UCC** | **CUC** | **CCC** | **CGA** | **ACG** | **GUC** | **GCC** | **GGU** |
| 4 | UCC | CUC | CCC | CGA | ACG | GUC | GCA | GGU |
| 5 | UCC | CUC | CCC | CGA | ACG | GUC | GCA | GGC |
| 6 | UCC | CUC | CCC | CGA | ACG | GUC | GCG | GGU |
| 7 | UCC | CUC | CCC | CGA | ACG | GUC | GCG | GGC |
| 8 | UCC | CUC | CCC | CGG | ACG | GUC | GCU | GGU |
| 9 | UCC | CUC | CCC | CGG | ACG | GUC | GCU | GGC |
| 10 | UCC | CUC | CCC | CGG | ACG | GUC | GCC | GGU |
| 11 | UCC | CUC | CCC | CGG | ACG | GUC | GCA | GGU |
| 12 | UCC | CUC | CCC | CGG | ACG | GUC | GCA | GGC |
| 13 | UCC | CUC | CCC | CGG | ACG | GUC | GCG | GGU |
| 14 | UCC | CUC | CCC | CGG | ACG | GUC | GCG | GGC |
